# Multi-modal Latent Factor Exploration of Atrophy, Cognitive and Tau Heterogeneity in Alzheimer’s Disease

**DOI:** 10.1101/462143

**Authors:** Nanbo Sun, Elizabeth C Mormino, Jianzhong Chen, Mert R Sabuncu, BT Thomas Yeo, for the Alzheimer’s Disease Neuroimaging Initiative

**Affiliations:** Department of Electrical and Computer Engineering, ASTAR-NUS Clinical Imaging Research Centre, N.1 Institute for Health and Memory Networks Program, National University of Singapore, Singapore; School of Medicine, Stanford University, Stanford, CA, USA; School of Electrical and Computer Engineering, Nancy E. and Peter C. Meinig School of Biomedical Engineering, Cornell University, Ithaca, NY, USA; Martinos Center for Biomedical Imaging, Massachusetts General Hospital, Charlestown, MA, USA; Center for Cognitive Neuroscience, Duke-NUS Graduate Medical School, Singapore; NUS Graduate School for Integrative Sciences and Engineering, National University of Singapore, Singapore

**Keywords:** Bayesian model, canonical correlation analysis, Executive function, Language, typical late-onset Alzheimer’s Disease

## Abstract

Individuals with Alzheimer’s disease (AD) dementia exhibit significant heterogeneity across clinical symptoms, atrophy patterns, and spatial distribution of Tau deposition. Most previous studies of AD heterogeneity have focused on atypical clinical subtypes, defined subtypes with a single modality, or restricted their analyses to a priori brain regions and cognitive tests. Here, we considered a data-driven hierarchical Bayesian model to identify latent factors from atrophy patterns and cognitive deficits *simultaneously*, thus exploiting the rich dimensionality within each modality. Unlike most previous studies, our model allows each factor to be expressed to varying degrees within an individual, in order to reflect potential multiple co-existing pathologies.

By applying our model to ADNI-GO/2 AD dementia participants, we found three atrophy-cognitive factors. The first factor was associated with medial temporal lobe atrophy, episodic memory deficits and disorientation to time/place (“MTL-Memory”). The second factor was associated with lateral temporal atrophy and language deficits (“Lateral Temporal-Language”). The third factor was associated with atrophy in posterior bilateral cortex, and visuospatial executive function deficits (“Posterior Cortical-Executive”). While the MTL-Memory and Posterior Cortical-Executive factors were discussed in previous literature, the Lateral Temporal-Language factor is novel and emerged only by considering atrophy and cognition jointly. Several analyses were performed to ensure generalizability, replicability and stability of the estimated factors. First, the factors generalized to new participants within a 10-fold cross-validation of ADNI-GO/2 AD dementia participants. Second, the factors were replicated in an independent ADNI-1 AD dementia cohort. Third, factor loadings of ADNI-GO/2 AD dementia participants were longitudinally stable, suggesting that these factors capture heterogeneity across patients, rather than longitudinal disease progression. Fourth, the model outperformed canonical correlation analysis at capturing associations between atrophy patterns and cognitive deficits.

To explore the influence of the factors early in the disease process, factor loadings were estimated in ADNI-GO/2 mild cognitively impaired (MCI) participants. Although the associations between the atrophy patterns and cognitive profiles were weak in MCI compared to AD, we found that factor loadings were associated with inter-individual regional variation in Tau uptake. Taken together, these results suggest that distinct atrophy-cognitive patterns exist in typical Alzheimer’s disease, and are associated with distinct patterns of Tau depositions before clinical dementia emerges.

**Highlights:**

1. Bayesian model reveals 3 atrophy-cognitive factors in typical AD from ADNI-GO/2
2. Replicated in independent ADNI-1 cohort; longitudinally stable within individuals
3. Triple cognitive dissociations among atrophy patterns suggest subtypes, not stages
4. Outperforms canonical correlation analysis
5. Factor loadings associated with spatial patterns of Tau uptake in MCI

## 1 Introduction

Alzheimer’s disease (AD) is a devasting neurodegenerative disease and the most common cause of dementia. Although AD commonly presents as an amnestic syndrome (Scheltens et al., 2016b), there is significant heterogeneity across individuals. For example, there exists multiple AD variants involving atypical clinical symptoms, such as visuospatial or language deficits (Lam et al., 2013; Lehmann et al., 2013). These atypical dysfunctions are often accompanied by corresponding atrophy patterns and tau deposition patterns. For example, participants with “visual variant” (posterior cortical atrophy, PCA) AD dementia might exhibit greater atrophy and tau deposition in posterior cortical regions (Ossenkoppele et al., 2015; 2016; Phillips et al., 2018). On the other hand, participants with “language variant” (logopenic aphasia, LPA) AD dementia might exhibit greater atrophy and tau deposition in temporal or parietal regions (Gorno-Tempini et al., 2011; Ossenkoppele et al., 2016; Phillips et al., 2018).

While the above studies focused on atypical clinically-defined AD subtypes, there is also heterogeneity within typical AD dementia (patients not clearly diagnosed with an atypical AD variant such as PCA or LPA). In cohorts comprising mostly typical AD dementia participants (e.g., ADNI), studies have demonstrated that participants exhibit distinct atrophy patterns or subtypes (Noh et al., 2014; Byun et al., 2015; Ferreira et al., 2017; Risacher et al., 2017; Young et al., 2018). Although details differ across studies, two common atrophy subtypes that have emerged across studies are defined by medial temporal or cortical atrophy (Noh et al., 2014; Byun et al., 2015; Zhang et al., 2016; Ferreira et al., 2017; Young et al., 2018). A less common subtype to be reported involves atrophy of the striatum, thalamus, and cerebellum (Zhang et al., 2016; Young et al., 2018). Unlike atypical clinically-defined AD variants, correspondences between atrophy subtypes and cognitive domains in the context of typical AD remain unclear, with most studies reporting worse overall cognition and faster disease progression in cortical subtypes (Noh et al., 2014; Byun et al, 2015; Risacher et al., 2017; but see Ferreira et al., 2017). An exception is our previous study, which found that AD dementia participants with the medial temporal atrophy subtype suffered worse memory deficits, while AD dementia participants with the cortical atrophy subtype suffered worse executive function deficits (Zhang et al., 2016). However, the subcortical atrophy subtype was not associated with a specific cognitive domain (Zhang et al., 2016).

The relatively weak correspondence between atrophy subtypes and cognitive domains in typical AD dementia highlights a need for improvement in subtyping algorithms. Previous studies have defined subtypes solely based on brain atrophy (Noh et al., 2014; Byun et al., 2015; Zhang et al., 2016; Ferreira et al., 2017; Young et al., 2018) followed by post-hoc associations with cognitive scores. Here, we proposed an extension of our previous model (Zhang et al., 2016) to estimate subtypes using structural MRI and neuropsychological testing scores simultaneously, identifying latent factors associated with distinct patterns of atrophy and cognitive deficits (Figure 1). The advantage of combining both modalities is that the associations between atrophy patterns and cognitive deficits are automatically encouraged by the model. Thus, this approach enabled a more thorough investigation of atrophy and cognitive profiles, rather than restricting analyses to previously defined composite measures of cognition (Zhang et al., 2016). This approach allows the possibility of new atrophy-cognitive factors emerging as a result of considering both atrophy and cognitive deficits simultaneously, thus providing insights into AD heterogeneity.

**Figure 1.**
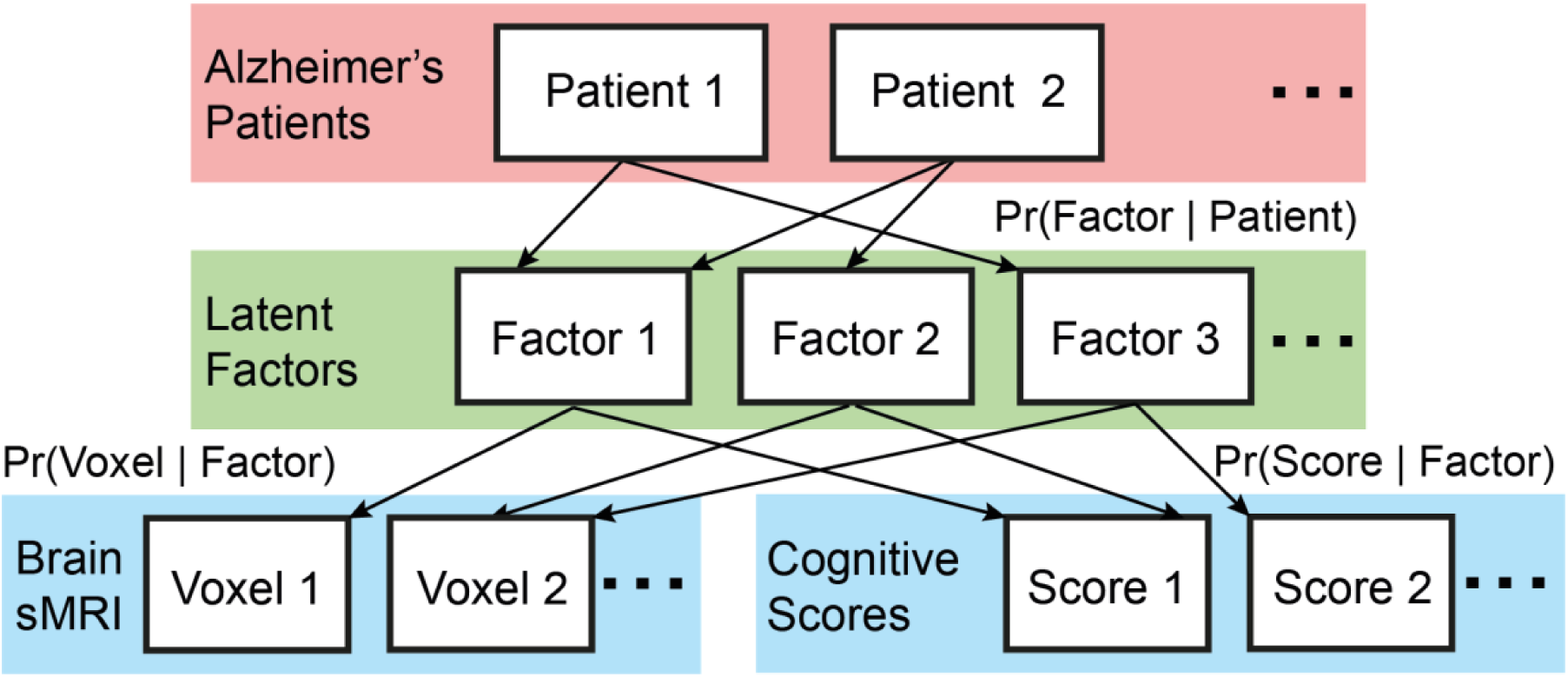
A Bayesian model of AD dementia patients, latent factors, brain structural MRI (sMRI) and cognitive scores. Our model allows each patient to express one or more latent factors. The factors are associated with distinct (but possibly overlapping) patterns of brain atrophy, as well as distinct (but possibly overlapping) profiles of cognitive deficits. The estimated model parameters are the probability that a patient expresses a particular factor (i.e., Pr(Factor | Patient)), the probability that a factor is associated with atrophy at a voxel (i.e., Pr(Voxel | Factor)) and the probability that a factor is associated with a cognitive score (i.e., Pr(Score | Factor)).

Consistent with our previous work (Zhang et al., 2016), a key feature of our modeling approach is that it allows the possibility that multiple latent factors are expressed to varying degrees within an individual. This is motivated by an extensive literature suggesting multiple etiologies underlying disease heterogeneity that are not mutually exclusive (Schneider et al., 2007; Schneider et al., 2009; Dickerson and Wolk, 2011; Murray et al., 2011; Franklin et al., 2015; Coutu et al., 2016; Ossenkoppele et al., 2016). For example, the atrophy-cognitive profiles of a patient might be 80% owing to factor 1, 15% to factor 2, and 5% to factor 3, whereas the atrophy-cognitive profiles of another patient might be 30% owing to factor 1, 65% to factor 2, and 5% to factor 3. This is in contrast to previous “winner-takes-all” subtyping studies, in which a participant is assigned to a single subtype (Noh et al., 2014; Byun et al., 2015; Ferreira et al., 2017). This motivates our use of the term “factors”, rather than “subtypes”.

Because typical AD dementia has a long prodromal stage, we also investigated whether associations between atrophy patterns and cognitive profiles estimated from AD dementia participants (Figure 1) could also be discerned in β-Amyloid positive (Aβ+) MCI participants. Given that brain atrophy and cognitive deficits are thought to occur later in the disease process (Jack et al., 2010; 2013), one might expect weaker associations between atrophy patterns and cognitive profiles in the MCI participants than in AD dementia participants. By contrast, Tau-mediated injury is thought to occur earlier in the disease process (Jack et al., 2010; 2013). The advent of ^18^F-Flortaucipir, a PET tracer with high affinity for paired helical filament Tau, provides an exciting opportunity to investigate regional NFTs *in vivo* (Chien et al., 2013). Thus, we also investigated whether the atrophy-cognitive factors (Figure 1) were associated with spatial patterns of Tau deposition in MCI participants.

Many studies have found strong relationships between tau depositions and cognitive deficits (Murray et al., 2011; Bejanin et al., 2017; Chiotis et al., 2017; Gordon et al., 2019). In the case of atypical clinically-defined AD subtypes, studies have demonstrated strong correspondences between tau deposition patterns and cognitive domains (Ossenkoppele et al., 2016; Phillips et al., 2018). For example, increased ^18^F-Flortaucipir uptake in the hippocampus, bilateral occipital lobe and left temporoparietal regions were associated with worse memory, visuospatial and language functions respectively (Ossenkoppele et al., 2016). However, in cohorts involving predominantly typical AD participants or only elderly nondemented participants, the correspondences between the spatial patterns of Tau deposition and cognitive domains are a lot less clear, with most studies showing greater global tau deposition associated with multiple cognitive domains, or greater regional tau depositions associated with global cognitive deficits (Johnson et al., 2016; Scholl et al., 2016; Maass et al., 2017; Mattsson et al., 2017; Aschenbrenner et al., 2018). Thus, our study seeks to extend these previous studies by clarifying whether the distinct tau deposition patterns are associated with different cognitive domains in MCI participants.

## 2 Materials and methods

### 2.1 Overview

There are three stages of analyses (Figure 2). First, a hierarchical multimodal Bayesian model (Figure 1) was applied to AD dementia participants from the ADNI-GO/2 (ADNI-GO and ADNI-2) database to extract factors based on patterns of atrophy and cognitive performance. The relationships between factor loadings and patient characteristics (e.g., age) were then examined. Second, we tested the generalizability, replicability and stability of the estimated factors, involving 10-fold cross-validation within ADNI-GO/2, independent replication using ADNI-1 and longitudinal data in ADNI-GO/2. We also compared our approach with canonical correlation analysis (CCA), which has been widely used to discover brain-behavior relationships (Smith et al., 2015).

**Figure 2.**
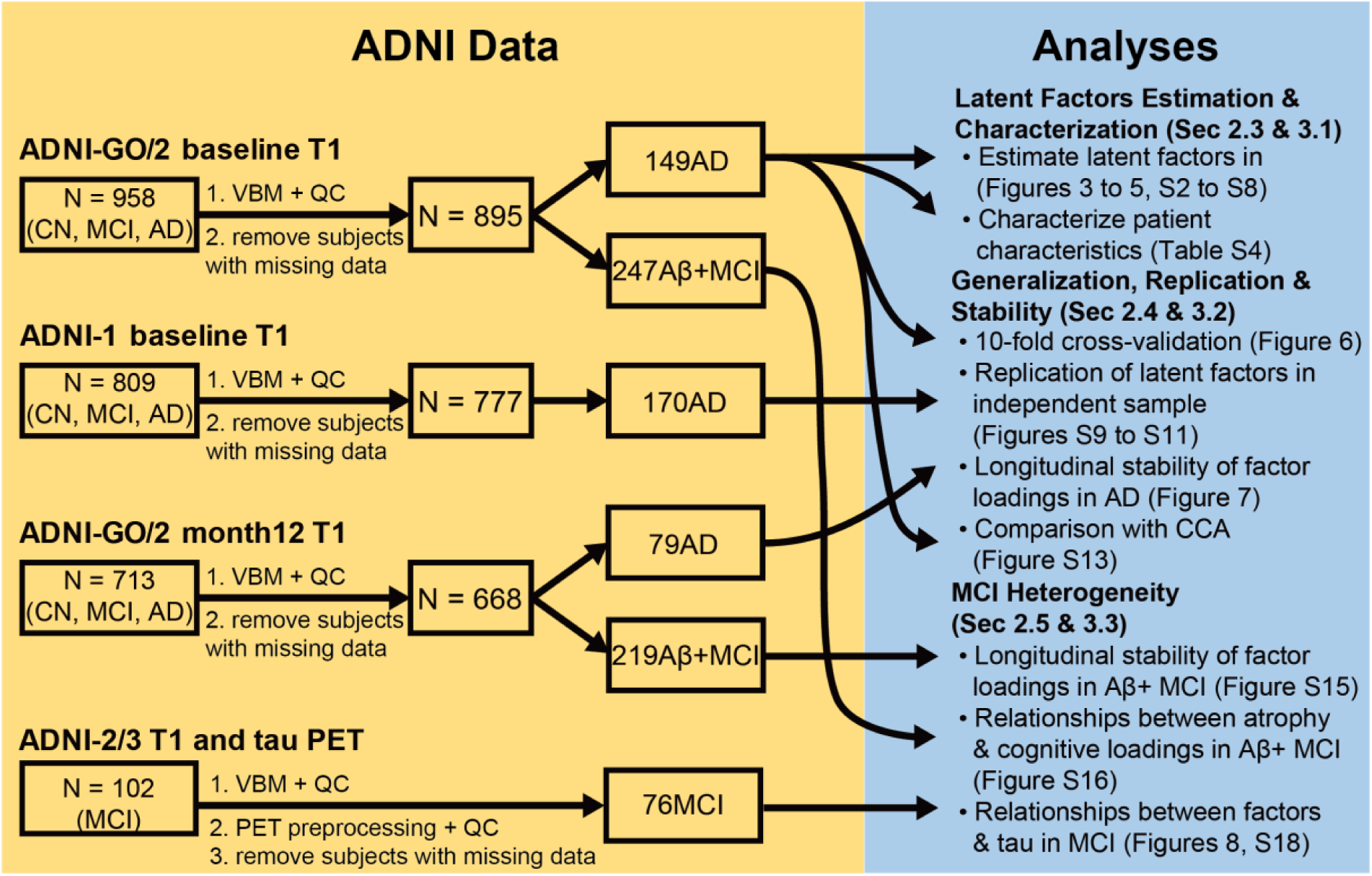
Overview of data and analyses. The yellow panel shows data from different ADNI cohorts and modalities. The blue panel shows the three stages of analyses in this study. Arrows indicate the data used in each analysis. We note that if Aβ+ was not specifically mentioned for a particular cohort, then this implied that the participants were not filtered based on Aβ+. The main reason for not filtering based on Aβ+ was to maximize sample size.

Third, to explore heterogeneity earlier in the disease process, we applied our model to ADNI-GO/2 Aβ+ MCI participants to extract factor loadings. Longitudinal stability of the factor loadings was examined. Relationships between atrophy and cognitive loadings were also explored. Finally, the association between factor loadings and the spatial pattern of Tau was also investigated in ADNI-2/3 MCI participants with available Tau PET. To maximize the number of participants, all MCI participants with available Tau PET were considered regardless of amyloid status^†^.

### 2.2 Data overview

This study utilized data from the ADNI database (adni.loni.usc.edu), which was established as a public-private partnership, led by Dr. Michael W. Weiner. ADNI’s main aim has been to test whether serial MRI, PET, other biological markers, and clinical and neuropsychological assessment can be combined to measure the progression of MCI and early AD. Written consent was obtained from all participants, and the study was approved by the Institutional Review Board at each participating institution. Demographic and clinical characteristics are presented in Table 1.

**Table 1.** Demographic and clinical characteristics of the various ADNI cohorts utilized in this study. Abbreviations: AD=Alzheimer’s disease, MCI=Mild cognitive impairment, CN=Cognitive normal, Aβ+=Amyloid beta positive, APOE=Apolipoprotein E, MMSE=Mini-mental state examination, CDR=Clinical dementia rating. Values represent mean (SD) min−max. The square brackets indicate the number of participants with available data. For example, [N=146] indicates that there were only 146 (out of a total of 149) ADNI-GO/2 AD participants with known amyloid status.

|  | ADNI-GO/2 baseline |  |  | ADNI-1 baseline |  |  |
| --- | --- | --- | --- | --- | --- | --- |
|  | AD | MCI | CN | AD | MCI | CN |
| N | 149 | 454 | 292 | 170 | 385 | 222 |
| Age at MRI acquisition | 74.6 (8.2)<br>55.7–90.3 | 71.7 (7.5)<br>55.0–91.4 | 73.0 (6.0)<br>56.2–90.0 | 75.5 (7.3)<br>55.2–91.0 | 74.9 (7.4)<br>54.8–89.3 | 76.0 (5.1)<br>60.0–89.7 |
| Age at AD onset | – | – | – | 72.0 (7.8)<br>53.0–91.0<br>[N=164] | – | – |
| % baseline age > 70 or<br>% age at AD onset > 65 | 113/149<br>75.84% | – | – | 137/164<br>83.54% | – | – |
| Sex (% male) | 58.4 | 54.2 | 46.2 | 50.6 | 64.2 | 51.4 |
| Education (years) | 15.8 (2.7)<br>9.0–20.0 | 16.1 (2.7)<br>9.0–20.0 | 16.6 (2.5)<br>8.0–20.0 | 14.7 (3.1)<br>6.0–20.0 | 15.7 (2.9)<br>6.0–20.0 | 16.0 (2.9)<br>6.0–20.0 |
| % Aβ+ | 88.4<br>[N=146] | 55.5<br>[N=445] | 33.5<br>[N=284] | 90.1<br>[N=91] | 75.4<br>[N=191] | 37.3<br>[N=110] |
| % APOE ε4 carriers | 67.6<br>[N=145] | 47.6<br>[N=450] | 29.6<br>[N=291] | 64.1<br>[N=170] | 53.5<br>[N=385] | 26.6<br>[N=222] |
| MMSE | 23.1 (2.1)<br>19.0–26.0 | 28.1 (1.7)<br>23.0–30.0 | 29.0 (1.2)<br>24.0–30.0 | 23.5 (2.0)<br>20.0–28.0 | 27.0 (1.8)<br>23.0–30.0 | 29.1 (1.0)<br>25.0–30.0 |
| CDR | 0.8 (0.3)<br>0.5–2.0 | 0.5 (0.0)<br>0.0–1.0 | 0.0 (0.0)<br>0.0–0.5 | 0.7 (0.3)<br>0.5–1.0 | 0.5 (0.0)<br>0.0–0.5 | 0.0 (0.0)<br>0.0–0.0 |
|  | ADNI-GO/2 month 12 |  |  | ADNI-2/3 Tau PET |  |  |
|  | AD | MCI | CN | AD | MCI | CN |
| N | 114 | 353 | 201 | – | 76 | – |
| Age at MRI acquisition | 75.2 (7.7)<br>56.4–91.6 | 73.1 (7.3)<br>56.7–92.5 | 73.7 (6.2)<br>56.0–90.2 |  | 76.4 (7.4)<br>57.8–92.2 |  |
| Sex (% male) | 60.5 | 56.4 | 46.8 |  | 56.6 |  |
| Education (years) | 15.9 (2.6)<br>9.0–20.0<br>[N=114] | 16.2 (2.7)<br>10.0–20.0<br>[N=353] | 16.6 (2.5)<br>12.0–20.0<br>[N=201] |  | 16.2 (2.9)<br>8.0–20.0<br>[N=64] |  |
| % Aβ+ | – | – | – | | 51.6<br>[N=62] | |
| % APOE ε4 carriers | 75.4<br>[N=114] | 44.2<br>[N=353] | 29.4<br>[N=201] | | 32.8<br>[N=64] | |
| MMSE | 22.7 (3.8)<br>11.0–30.0 | 27.8 (2.0)<br>20.0–30.0 | 28.8 (1.4)<br>20.0–30.0 |  | 27.9 (2.1)<br>22.0–30.0 |  |
| CDR | 0.9 (0.4)<br>0.5–2.0<br>[N=113] | 0.5 (0.1)<br>0.0–1.0<br>[N=349] | 0.1 (0.2)<br>0.0–0.5<br>[N=196] |  | 0.4 (0.2)<br>0.0–1.0<br>[N=76] |  |

According to ADNI’s inclusion criteria (page 27 of https://adni.loni.usc.edu/wp-content/uploads/2008/07/adni2-procedures-manual.pdf), participants must be at least 55 years old when diagnosed with AD dementia. ADNI also excluded participants whose MMSE were less than 20. In practice, AD dementia participants were on average 75 years old and 85% of the AD dementia participants were above 65 years old (Table 1). Given that the cognitive tests in ADNI were not designed to detect atypical clinically-defined AD subtypes (e.g., logopenic subtypes), we could not rule out the presence of atypical clinically-defined AD subtypes. However, atypical AD variants usually have faster disease progression, early disease onset and are relatively rare. Therefore, we expect the occurrence of atypical AD variants in the ADNI cohort to be relatively low given the age range of ADNI AD dementia participants and the exclusion of participants with MMSE less than 20. Thus, we interpret the ADNI cohort as comprising predominantly typical late-onset AD dementia participants, consistent with the interpretation of some previous work with ADNI (Meda et al., 2012; Iturria-Medina et al., 2016). It is also worth noting that late-onset AD is typically defined as being diagnosed with AD dementia at 65 years or older (American Psychiatric Association, 2013). Unfortunately, the age of onset was not available in ADNI-GO/2, while amyloid status of many participants was not available in ADNI-1. Since the gap between the age of onset and baseline age was around 3.5 years in ADNI-1 (Table 1), Section 3.1 will include control analyses involving ADNI-GO/2 Aβ+ AD dementia participants, who were at least 70 years old at baseline. Thus, the sensitivity analysis in Section 3.1 was performed to ensure that factors derived with the larger sample of AD participants were not driven by heterogeneity within Aβ- AD and/or the subset of younger ADNI participants with AD that could reflect traditional atypical presentations.

### 2.3 Latent Factors Estimation & Characterization

#### 2.3.1 Structural MRI

Baseline MRI data from 158 AD dementia patients, 489 MCI participants, and 311 cognitively normal (CN) participants from ADNI-GO/2 were used to define a study specific template for voxel-based morphometry (VBM; Figure 2). All structural MRI (T1-weighted, 3.0T) were acquired with Sagittal MP-RAGE/IR-SPGR sequence and followed a standardized preprocessing protocol: Gradwarp, B1 non-uniformity and N3 correction (http://adni.loni.usc.edu/methods/mri-analysis/mri-pre-processing/).

The structural MRI were analyzed using the CAT12 VBM software (http://www.neuro.uni-jena.de/cat/), yielding subject-specific gray-matter (GM) density maps in MNI space. We then applied log_10_ transformation to the GM density images and regressed the effects of age, sex and intracranial volume (ICV) from all participants. Regression coefficients were estimated from only CN participants to retain any interaction between AD and participants’ characteristics (e.g., interaction between AD and age). The regression coefficients were then utilized to compute residuals for all participants. For more details about the VBM and regression procedures, see Supplementary Methods S1.

#### 2.3.2 Cognitive scores

Cognitive scores from the Alzheimer’s Disease Assessment Scale-Cognitive (ADAS-Cog13), Mini Mental State Exam (MMSE), Boston Naming Test, animal category of Category Fluency Test, Clock Drawing Test, Clock Copying Test, Logical Memory, Rey Auditory Verbal Learning Test and Trail Making Test were considered. See https://adni.loni.usc.edu/wp-content/uploads/2008/07/adni2-procedures-manual.pdf for detailed descriptions of these tests. In the case of MMSE, subsets of the 30 item scores were summed together into 8 domains (Table S1; Folstein et al., 1975): “MMSE: Orientation to time”, “MMSE: Orientation to place”, “MMSE: Immediate Recall”, “MMSE: Attention”, “MMSE: Delayed Recall”, “MMSE: Language”, “MMSE: Repetition” and “MMSE: Complex Commands”. This resulted in a total of 32 scores (Table S1).

Table S5 shows the standard deviation of the cognitive scores across CN participants within the ADNI-1 and ADNI-GO/2 cohorts. The five cognitive scores with lowest variance were consistent across both cohorts and corresponded to “ADAS: Recall Instructions”, “MMSE: Language”, “ADAS: Comprehension”, “MMSE: Immediate Recall” and “ADAS: Spoken Language”. Among these five cognitive scores with limited range, the following scores were combined since they reflect similar cognitive functions: (1) “MMSE: Language” and “ADAS: Naming”, (2) “MMSE: Immediate Recall” and “MMSE: Delayed Recall” and (3) “ADAS: Spoken Language” and “ADAS: Comprehension”. We did not think that the remaining cognitive score “ADAS: Recall Instructions” was tapping into similar cognitive function as other scores, so we simply remove the score from consideration. The variance of the combined scores are shown in Table S6. Thus, we ended up with 28 scores (Table S3).

Among the 158 ADNI-GO/2 AD dementia participants, 9 were excluded because more than 10 cognitive scores were missing. Of the remaining 149 participants, 124 had all scores, while 25 participants had less than five missing scores. The general linear model (GLM) was used to impute the missing scores in the 25 participants (Section 2.7 of Enders, 2010). Briefly, for each participant with missing scores, we fitted a GLM to all participants (CN, MCI and AD) without missing scores, where the observed scores were independent variables and the missing scores were dependent variables. The estimated regression coefficients were used to fill in the missing scores of the participant with missing scores.

Consistent with the structural MRI processing, we regressed out effects of age, sex and ICV from all scores using regression coefficients estimated from the CN participants. See Supplementary Methods S1 for details.

#### 2.3.3 Multimodal Bayesian model

We have previously utilized a hierarchical Bayesian model, latent Dirichlet allocation (LDA; Blei et al., 2003), to encode the premise that each AD patient expresses one or more latent factors, associated with distinct patterns of brain atrophy (Zhang et al., 2016). LDA (and its variants) have also been successfully used to extract overlapping brain networks from functional MRI (Yeo et al., 2014) and meta-analytic data (Bertolero et al., 2015; Yeo et al., 2015). In the current analysis, we considered an extension of the LDA model to incorporate two modalities (gray matter atrophy and neuropsychological testing scores), herein referred to as multi-modality LDA (MMLDA). In this framework, each AD patient expresses one or more latent factors, each of which is associated with distinct (but possibly overlapping) atrophy patterns and distinct (but possibly overlapping) cognitive deficits (Figure 1). The model is mathematically equivalent to that proposed by Putthividhya and colleagues (2007). Details are found in Supplementary Methods S2 and S3.

We considered VBM and cognitive scores of all ADNI-GO/2 participants with AD dementia (N = 149). Since most participants were Aβ+ (88%; Table 1), we did not exclude the small number of Aβ- participants, in order to maximize the number of participants. The VBM and cognitive scores of 149 ADNI-GO/2 participants with AD dementia were first z-normalized with respect to the CN participants (see Supplementary Methods S4 for details). Given the normalized VBM and cognitive scores of the 149 participants, as well as a predefined number of latent factors *K*, a variational expectation-maximization (VEM) algorithm was used to estimate the probability of a patient expressing a latent factor [Pr(Factor | Patient)], probability that a factor was associated with atrophy at a voxel [Pr(Voxel | Factor)] and the probability that a factor was associated with a cognitive score [Pr(Score | Factor)]. See Table 2 for a summary of the terminologies.

**Table 2.**
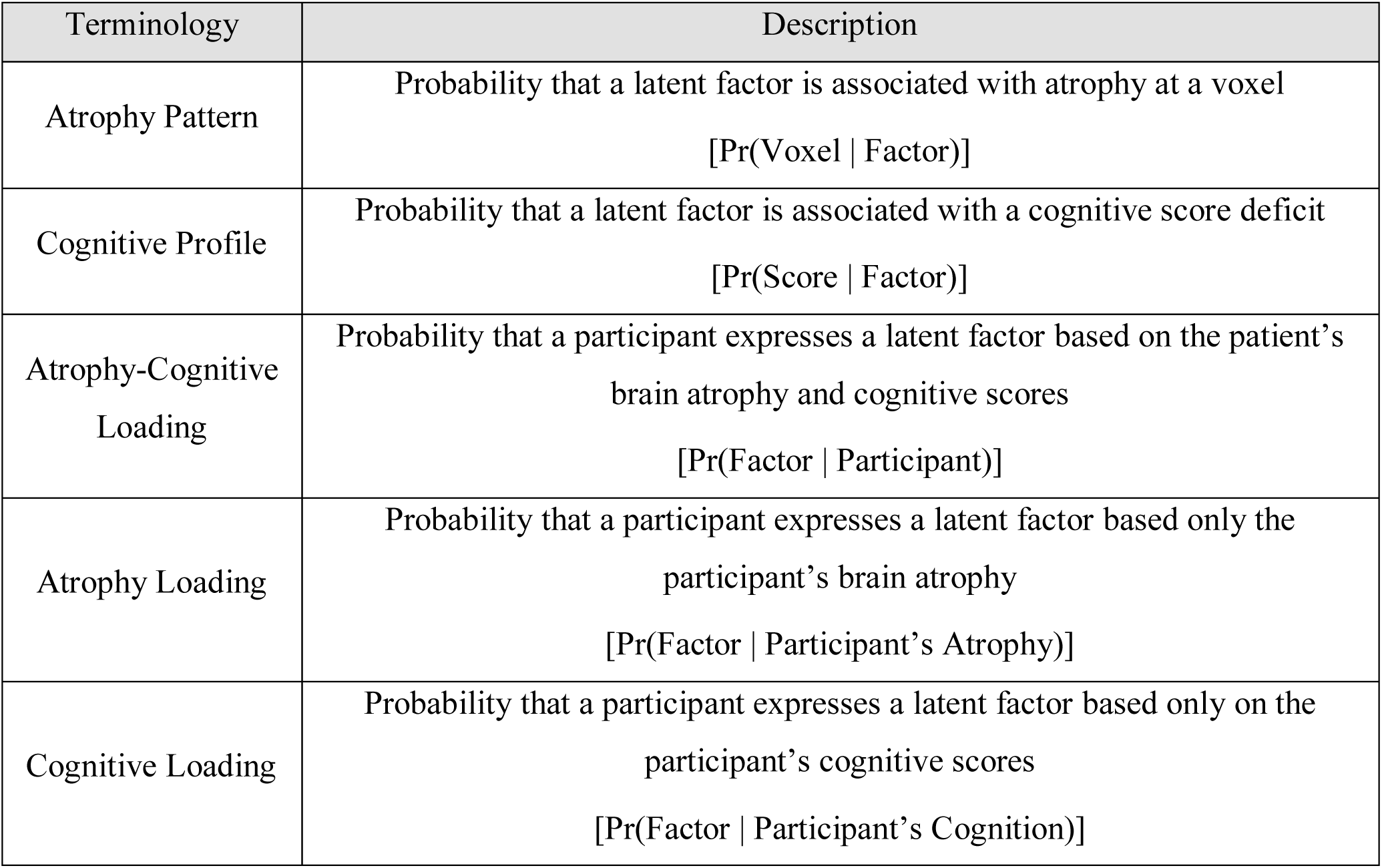
Summary of terminologies and their descriptions.

An important model parameter is the number of latent factors *K*. We considered *K* = 2, 3 or 4. As will be explained in the Results, the four-factor solution was not interpretable, and so we did not explore beyond four factors. For the remainder of this paper, we will largely focus on the three-factor solution.

#### 2.3.4 Visualization

The probability of a voxel being associated with a factor (Pr(Voxel | Factor)) can be thought of as a brain map. For visualization, the brain map is projected from MNI152 (Figure S2 and S10) to fsaverage space (Figure 3 and S9; Wu et al., 2018). The fsaverage space is the official FreeSurfer surface coordinate system obtained by averaging the cortical surfaces of 40 participants (Dale et al., 1999; Fischl et al., 1999a). Subcortical structures are illustrated using coronal slices. Multiple coronal slices are also shown in supplemental figures.

**Figure 3.**
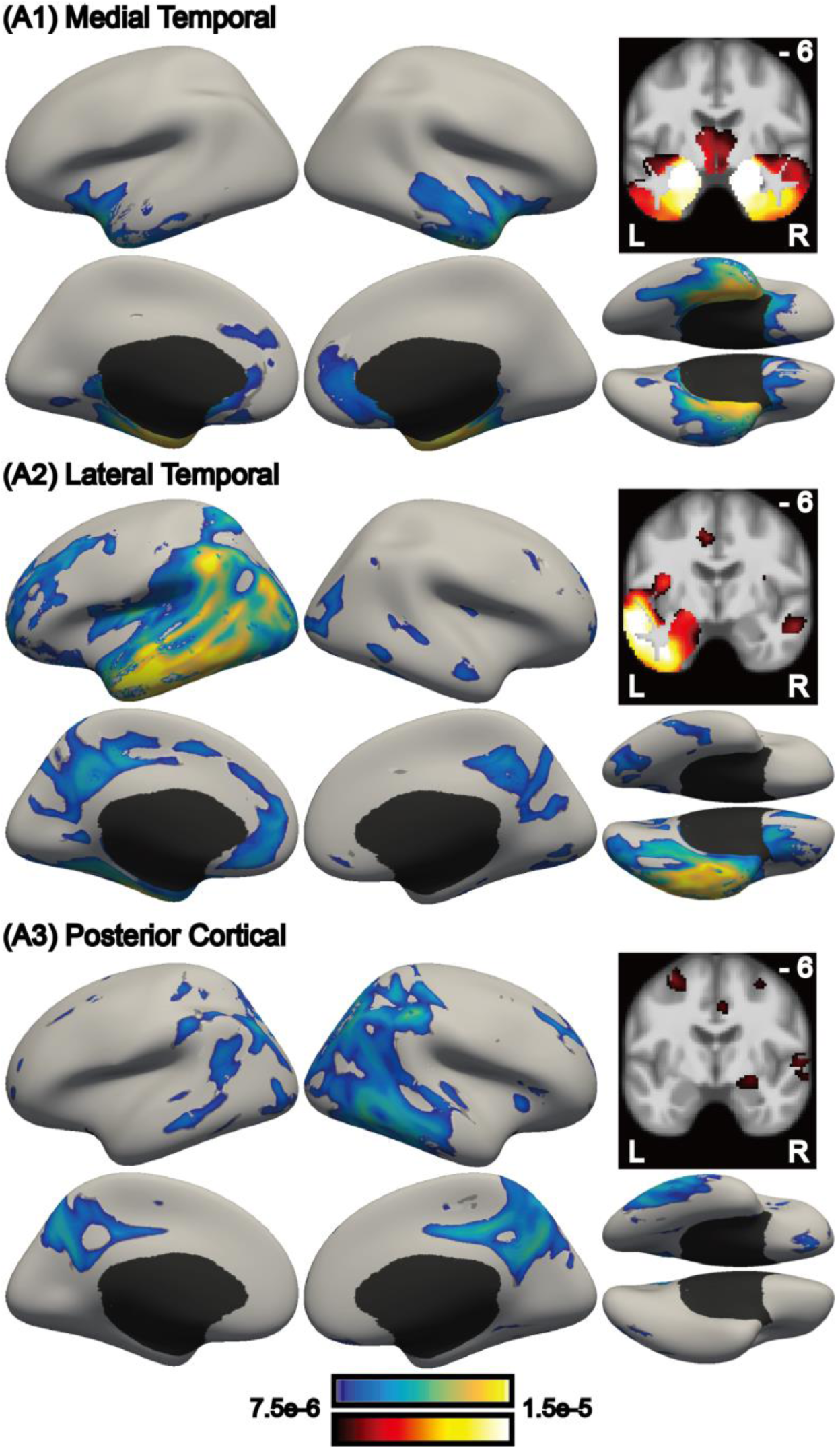
Probabilistic atrophy maps of three latent factors in ADNI-GO/2. Brighter color indicates higher probability of atrophy at that voxel for a particular factor (i.e., Pr(Voxel | Factor)). Each of the three factors was associated with a distinct pattern of brain atrophy. Coronal slices are found in Figure S2.

#### 2.3.5 Relationship between factors & patient characteristics

We explored how patient characteristics (age at time of structural MRI scan, education, Amyloid, and APOE genotype) varied across the three latent factors using GLMs for continuous variables and logistic regression for binary variables. In all cases, patient characteristics were the independent variables, while factor loadings were the dependent variables. Statistical tests were then performed to determine if there were overall differences across all factors, and whether there were differences between pairs of factors. See Supplementary Methods S5 for details.

### 2.4 Generalization, Replication & Stability

#### 2.4.1 Inference of atrophy loading, cognitive loading or atrophy-cognitive loading

After the probabilistic atrophy maps Pr(Voxel | Factor) and probabilistic cognitive deficits Pr(Score | Factor) were estimated, they could be used to infer the factor composition (or loading) of a new participant using only atrophy [“atrophy loading” or Pr(Factor | Atrophy of Participant)], only cognitive scores [“cognitive loading” or Pr(Factor | Cognition of Participant)] or both atrophy and cognitive scores [“atrophy-cognitive loading” or Pr(Factor | Participant)]. See Table 2 for a summary of the terminologies.

#### 2.4.2 10-fold cross-validation

To test whether the association between cognitive profiles and atrophy patterns can generalize to new participants within the same dataset, we performed 10-fold cross-validation among the 149 ADNI-GO/2 AD dementia participants. Briefly, this involves randomly splitting the 149 participants into ten folds (groups) of roughly the same size. For each test fold, the multimodal Bayesian model parameters were estimated from the remaining nine training folds, and then used to infer factor loadings in the test fold participants using only atrophy (“atrophy loading”) or only cognitive scores (“cognitive loading”). The prediction quality is measured by the correlation between atrophy and cognitive loadings across the test fold participants.

#### 2.4.3 Replication in ADNI-1

The multimodal Bayesian model was also applied to an independent cohort of 170 ADNI-1 patients with AD dementia to see if the resulting factors were similar across ADNI cohorts. The model parameters were estimated completely independently from the previous analyses involving ADNI-GO/2 participants.

#### 2.4.4 Longitudinal stability of factor loadings in AD dementia

To ensure the factors were not simply reflecting disease stages, we examined longitudinal factor stability. More specifically, we considered the subset of 149 AD dementia (from the ADNI-GO/2 cohort) with one-year follow-up structural MRI and the full set of cognitive scores, yielding 79 dementia participants. Factor loadings of the participants were inferred using only structural MRI (i.e., atrophy loading), only cognitive scores (i.e., cognitive loading) or both structural MRI and cognitive scores (i.e., atrophy-cognitive loading) at both time points separately. If the factors mostly reflected disease stages (rather than subtypes), then we expected a big longitudinal shift in the factor loadings (e.g., factor 3 loading might increase over time).

#### 2.4.5 Comparison with canonical correlation analysis (CCA)

CCA is a multivariate data-driven statistical technique that maximizes the correlations between two data modalities by deriving canonical components, which are optimal linear combinations of the original data. Because CCA has been widely used to discover brain-behavior relationships (Smith et al., 2015), we applied CCA to the ADNI-GO/2 data to investigate the atrophy-cognitive modes extracted with this alternative method (Figure 1). For more details, see Supplementary Methods S6.

### 2.5 MCI Heterogeneity

AD has a long prodromal stage before overt dementia. Therefore, we were interested in exploring whether the atrophy-cognitive factors estimated from the AD dementia participants were relevant in an earlier disease stage (i.e., MCI).

#### 2.5.1 Longitudinal stability of factor loadings in Aβ+ MCI

Similar to Section 2.4.4, we examined longitudinal factor stability to ensure the factors were not simply reflecting disease stages. Of the 454 ADNI-GO/2 MCI participants, 445 had Amyloid PET data available, and 247 of 445 were Aβ+ (see Supplementary Methods S9 for details). We further considered the subset of participants with one-year follow-up structural MRI and the full set of cognitive scores, yielding 219 Aβ+ MCI participants. The multimodal Bayesian model parameters estimated from the ADNI-GO/2 AD dementia participants were used to infer factor loadings of the participants using only structural MRI (i.e., atrophy loading), only cognitive scores (i.e., cognitive loading) or both structural MRI and cognitive scores (i.e., atrophy-cognitive) at both time points separately. If the factors mostly reflected disease stages (rather than subtypes), then we expect a longitudinal shift in the factor loadings over time.

#### 2.5.2 Relationships between atrophy & cognitive loadings in MCI

To explore the associations between atrophy patterns and cognitive deficits in MCI participants, the multimodal Bayesian model parameters estimated from the ADNI-GO/2 AD dementia participants were used to infer factor loadings in the 247 Aβ+ MCI participants using only atrophy (“atrophy loading”) or only cognitive scores (“cognitive loading”). To examine the relationships between cognitive and atrophy loadings, GLM was utilized. For example, factor 1 cognitive loading would be the dependent variable, while the atrophy loadings would be the independent variables. Age and sex were included as nuisance variables. To be more explicit, let us denote memory loading by *h,* medial temporal loading by *m*, lateral temporal loading by *l*, and posterior cortical loading by c. Age *x*_1_and sex *x*_2_ were included as nuisance variables, so the final GLM was h = β_0_ + β_*l*_*l* + β_*c*_*c* + β_1_*x*_1_ + β_2_*x*_2_ + 𝜖, where β’s denote the regression coefficients, and 𝜖 is the residual. The medial temporal loading *m* was implicitly modeled because *m* + *l* + *c* = 1. The GLM was repeated with factor 2 cognitive loading and then factor 3 cognitive loading as the dependent variables. For each GLM, statistical tests were performed to determine if there were overall differences across all factors, and whether there were differences between pairs of factors. See Supplementary Methods S7 for details.

#### 2.5.3 Relationships between factors & Tau in MCI

There were 102 participants with MCI from ADNI-2/3 (ADNI-2 and ADNI-3), who had accompanying ^18^F-Flortaucipir PET scans. Participants were scanned beginning at 75 min post-injection, for 30 min (6 × 5 min frames) and each scan underwent the following preprocessing steps: Co-registered Dynamic, Averaged, Standardized Image and Voxel Size and Uniform Resolution (http://adni.loni.usc.edu/methods/pet-analysis/pre-processing/).

For ADNI-2 participants, the PET scan was not acquired at baseline, but during a follow-up visit. Since structural MRI and cognitive tests were not collected for every follow-up visit, this resulted in substantial missing data. In total, 26 participants were excluded because of missing structural MRI or more than five missing cognitive scores. Of the remaining 76 participants, 29 participants had all cognitive scores, while 47 participants had three or less missing scores. Like before, missing scores were imputed using the GLM based on all ADNI-GO/2 participants (CN, MCI and AD) without missing scores. The participants were not filtered based on Amyloid status because including only Aβ+ participants will lead to too few participants (Table 1).

All structural MR images were processed using FreeSurfer 6.0 (https://surfer.nmr.mgh.harvard.edu/) to obtain the parcellation of the cerebellum in native T1 space (Dale et al., 1999; Fischl et al., 1999a, b; Fischl et al., 2001; Ségonne et al., 2007; Greve and Fischl, 2009). All PET images were then co-registered to the corresponding T1 images. Voxelwise standardized uptake value ratio (SUVR) images were created by dividing each voxel by the mean value in the cerebellar gray matter. Finally, the PET images were transformed to the same template space as the VBM analysis and downsampled to 2mm (which was the same resolution as the GM density images).

To investigate the relationships between the spatial heterogeneity of Tau depositions and factor loadings, each probabilistic atrophy map (Pr(Voxel | Factor)) was thresholded to obtain a mask containing the top 5% of the voxels (Figure S1). The SUVR signal was averaged across voxels within each mask to obtain the regional Tau deposition for each factor. Similar results were obtained if we utilized a 2.5% threshold or a 10% threshold.

To determine if the regional Tau deposition were associated with the factor loadings, the GLM was applied for each factor loading. For example, the atrophy-cognitive loading of factor 1 (Pr(Factor 1 | Patient)) would be the dependent variable, while the Tau deposition in each of the three atrophied regions would be the independent variables using the 5%-thresholded masks described above. Age and sex were included as nuisance variables. For each GLM, statistical tests were performed to determine if there were overall differences across all factors, and whether there were differences between pairs of factors. See Supplementary Methods S8 for details.

Finally, for the purpose of visualization, at each voxel, we regressed out age and sex from the Tau PET signal and then correlated the residual with the atrophy-cognitive factor loadings across participants, resulting in three Tau-atrophy-cognitive maps. For visualization, the maps were projected to the FreeSurfer fsaverage surface (Wu et al., 2018). Furthermore, we correlated the Tau-atrophy-cognitive maps (Figure S18) with the atrophy patterns (Figure 3) and utilized permutation test to determine the significance of the correlations.

### 2.6 Statistical Analyses

In the case of logistic regression, the likelihood ratio test was employed to test for differences between factors. In the case of GLMs, the F-test was employed. In the case of Pearson’s correlations (e.g., 10-fold cross-validation), p values were computed using the Student’s t distribution. All tests were two-sided. Furthermore, since multiple statistical tests (patients’ characteristics analyses, 10-fold cross-validation, associations between atrophy and cognitive loadings among Aβ+ MCI participants, and associations between Tau deposition and factor loadings among MCI participants) were performed in this study, all p values were corrected using a false discovery rate (FDR) of q = 0.05.

### 2.7 Code and data availability

The code used in this paper, study-specific VBM templates (ADNI-1 and ADNI-GO/2), atrophy-cognitive factors and factor loadings of participants are publicly available at https://github.com/ThomasYeoLab/CBIG/tree/master/stable_projects/disorder_subtypes/Sun2019_ADJointFactors. Researchers can estimate factor loadings of new participants based on our estimated model or re-estimate the model based on their own subjects. The paper utilized data from the publicly available ADNI database (http://adni.loni.usc.edu/data-samples/access-data/).

## 3 Results

### 3.1 Latent Factor Estimation & Characterization

#### 3.1.1 Latent atrophy-cognitive factors in AD dementia

Using a multimodal Bayesian model, we estimated three latent factors from 149 ADNI-GO/2 AD dementia participants that captured covariance between patterns of atrophy and cognitive testing scores. The three latent factors involved distinct atrophy patterns (Figure 3; Figure S2) and corresponding cognitive testing profiles (Figure 4). The first factor was associated with atrophy in the medial temporal lobe (MTL) and was associated with episodic memory tests (delayed and immediate recall measures), as well as orientation measures (“MTL-Memory”). The second factor was associated with atrophy in left lateral temporal cortex and diffuse atrophy across the default network, and was associated with multiple tests assessing language (“Lateral Temporal-Language”). Finally, the third factor was associated with atrophy in the posterior bilateral cortex (including parietal and lateral temporal regions) and portions of frontal cortex, and was associated with tests of executive function and visuospatial function (“Posterior Cortical-Executive”).

**Figure 4.**
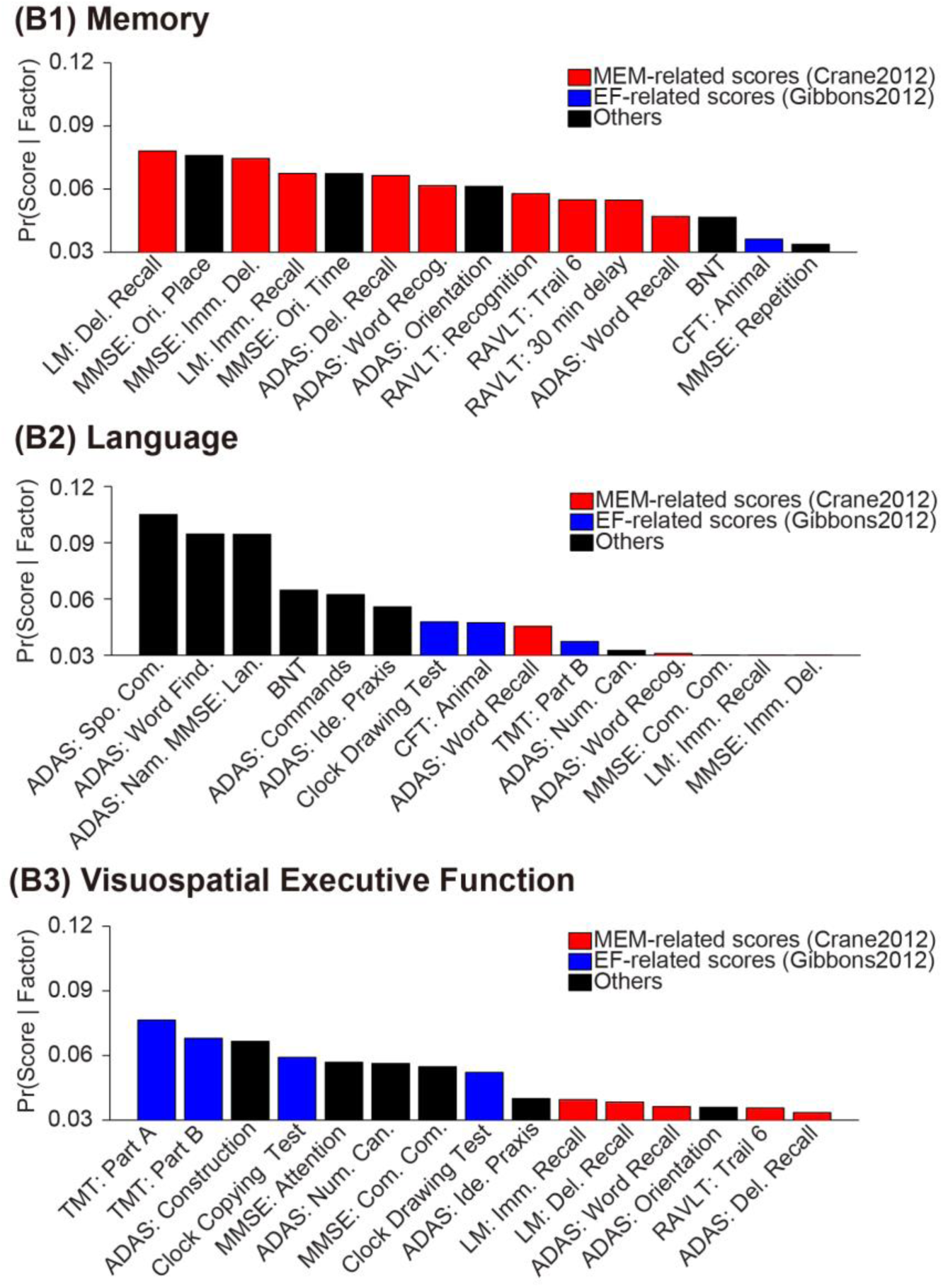
Probabilistic cognitive deficits of three latent factors in ADNI-GO/2. Red bars indicate cognitive scores associated with memory (as defined by Crane et al., 2012). Blue bars indicate cognitive scores associated with executive function (as defined by Gibbons et al., 2012). The remaining cognitive scores were colored black. Vertical axis corresponds to the probability of a cognitive score being associated with a factor (i.e., Pr(Score | Factor)). Only the top 15 scores are shown. ADAS: Alzheimer’s Disease Assessment Scale; MMSE: Mini Mental State Exam; CFT: Category Fluency Test; LM: Logical Memory; RAVLT: Rey Auditory Verbal Learning Test; TMT: Trail Making Test.

As a control analysis, we re-estimated the three factors using a subset of 97 Aβ+ AD dementia participants greater than 70 years old. Correlations between atrophy patterns from ADNI-GO/2 AD (Figure S2) and ADNI-GO/2 Aβ+ AD participants with age > 70 (Figure S3) were 0.93, 0.87 and 0.77 for factors 1, 2 and 3 respectively. Correlations between corresponding cognitive profiles from ADNI-GO/2 AD (Figure 4) and ADNI-GO/2 Aβ+ AD (Figure S4) participants with age > 70 were 0.99, 0.91 and 0.90 for factors 1, 2 and 3 respectively. Given the strong agreement between the original and control analyses, we chose to use the factors estimated from all 149 ADNI-GO/2 AD dementia participants in the following analyses.

It is worth pointing out that Pr(Score | Factor) must sum to one over all scores because it is a probability distribution. Therefore, if certain cognitive tests from a particular cognitive domain were over-represented in the dataset, then on average Pr(Score | Factor) for the particular cognitive domain would be lower. Here, there were more cognitive tests in ADNI involving memory and visuospatial executive function, so Pr(Score | Factor) was generally lower for the Posterior Cortical-Executive and MTL-Memory factors compared with the Lateral Temporal-Language factor (Figure 4).

#### 3.1.2 Number of factors

An important model parameter is the number of latent factors. This is an unsolved problem in machine learning with no consensus on the best approach for estimating the number of factors. In this paper, we first estimated the two-factor model and then continued to increase the number of estimated factors as long as two criteria were satisfied. First, we preferred more factors as long as the relationships between the atrophy pattern and cognitive profiles remained biologically plausible. Because biological plausibility was subjective, we considered a second quantitative criterion, which was that the atrophy-cognitive relationships generalized to new participants within a 10-fold cross-validation procedure (Section 3.2.1).

The two-factor model revealed two factors that were very similar to the first and third factors of the three-factor model. More specifically, there was one factor associated with medial temporal atrophy (Figure S5A1) and memory-related cognitive deficits (Figure S6B1), whereas the second factor was associated with posterior cortical atrophy (Figure S5A2) and visuospatial/executive function cognitive deficits (Figure S6B2). Therefore, the two-factor estimates were biologically plausible. However, according to the first criterion, we preferred the three-factor model to the two-factor model because of the additional Lateral Temporal – Language factor. The three-factor model also generalized to new participants (Section 3.2.1), which satisfied the second criterion.

On the other hand, the four-factor model revealed three factors (Figures S7A1-3, S8B1-3), which were similar to factors in three-factor model. However, the fourth factor was associated with medial temporal atrophy in addition to subcortical atrophy in the basal ganglia and thalamus (Figure S7A4). Thus, the atrophy pattern was not constrained to a known brain system. The fourth factor was also associated with a mixture of deficits that appeared to cut across multiple cognitive domains (Figure S8B4). Therefore, overall, we felt that the fourth atrophy-cognitive factor did not seem biologically plausible. The fourth factor also did not generalize to new participants (Section 3.2.1). Therefore, we preferred the three-factor model to the four-factor model, and we did not estimate the five-factor model.

#### 3.1.3 Characteristics of factor loadings

Examination of factor loadings revealed that the majority of the 149 AD dementia participants expressed multiple latent atrophy factors rather than predominantly expressing a single atrophy factor (Figure 5). Furthermore, there was no significant association between the factor loading and education, sex, or APOE (Table S4). However, the third factor (Posterior Cortical-Executive) was associated with younger age (at time of scan) than the other two factors (p = 2e-4; Table S4).

**Figure 5.**
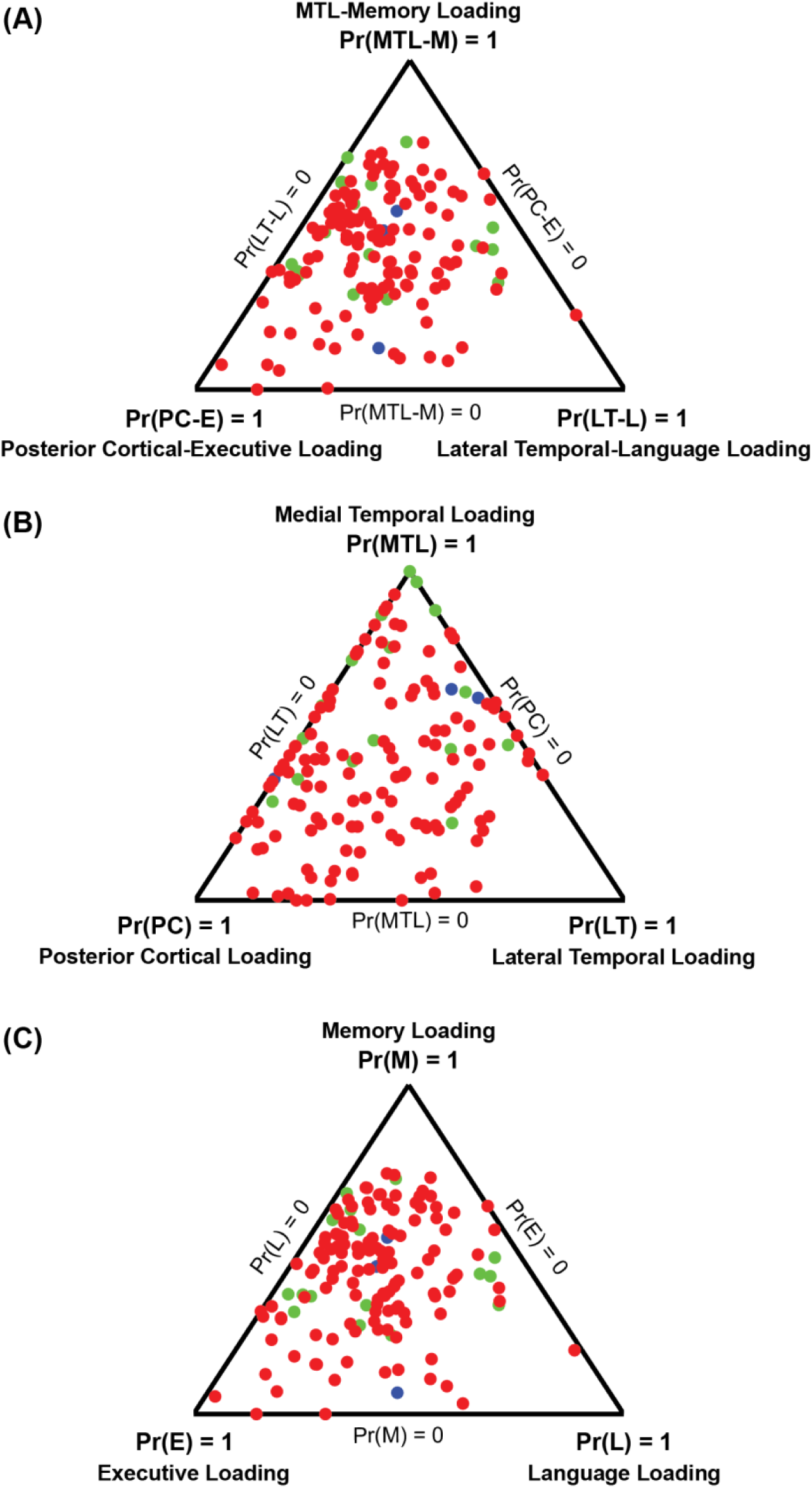
Factor loadings of 149 AD dementia patients. Each patient corresponds to a dot, with location (in barycentric coordinates) representing the factor loading. Color indicates Amyloid status: red for Aβ+, green for Aβ-, and blue for unknown. Corners of the triangle represent pure factors; closer distance to the respective corner indicates higher probability for the respective factor. (A) Atrophy-cognitive loading estimated using both brain atrophy and cognitive scores. “MTL-M” indicates Medial Temporal Lobe-Memory. “LT-L” indicates Lateral Temporal-Language. “PC-E” indicates Posterior Cortical-Executive. (B) Atrophy loading estimated using only structural MRI. (C) Cognitive loading estimated using only cognitive scores. Most dots are far from the corners, suggesting that most patients expressed multiple factors.

### 3.2 Generalization, Replication & Stability

#### 3.2.1 Atrophy pattern can be used to predict cognitive deficits with modest accuracies

To test whether the atrophy pattern of an out-of-sample participant could be used to predict his/her cognitive deficit profile, we performed 10-fold cross-validation among the AD dementia participants in ADNI-GO/2. We found that greater medial temporal atrophy predicted worse memory function with modest accuracy (Figure 6; r = 0.34, p = 2e-5), greater lateral temporal atrophy predicted worse language performance with modest accuracy (Figure 6; r=0.30, p=2e-4), and greater posterior cortical atrophy predicted worse visuospatial executive function with modest accuracy (Figure 6; r=0.30, p=3e-4). For 10-fold cross-validation results for the four-factor model, see Supplementary Results S1.

**Figure 6.**
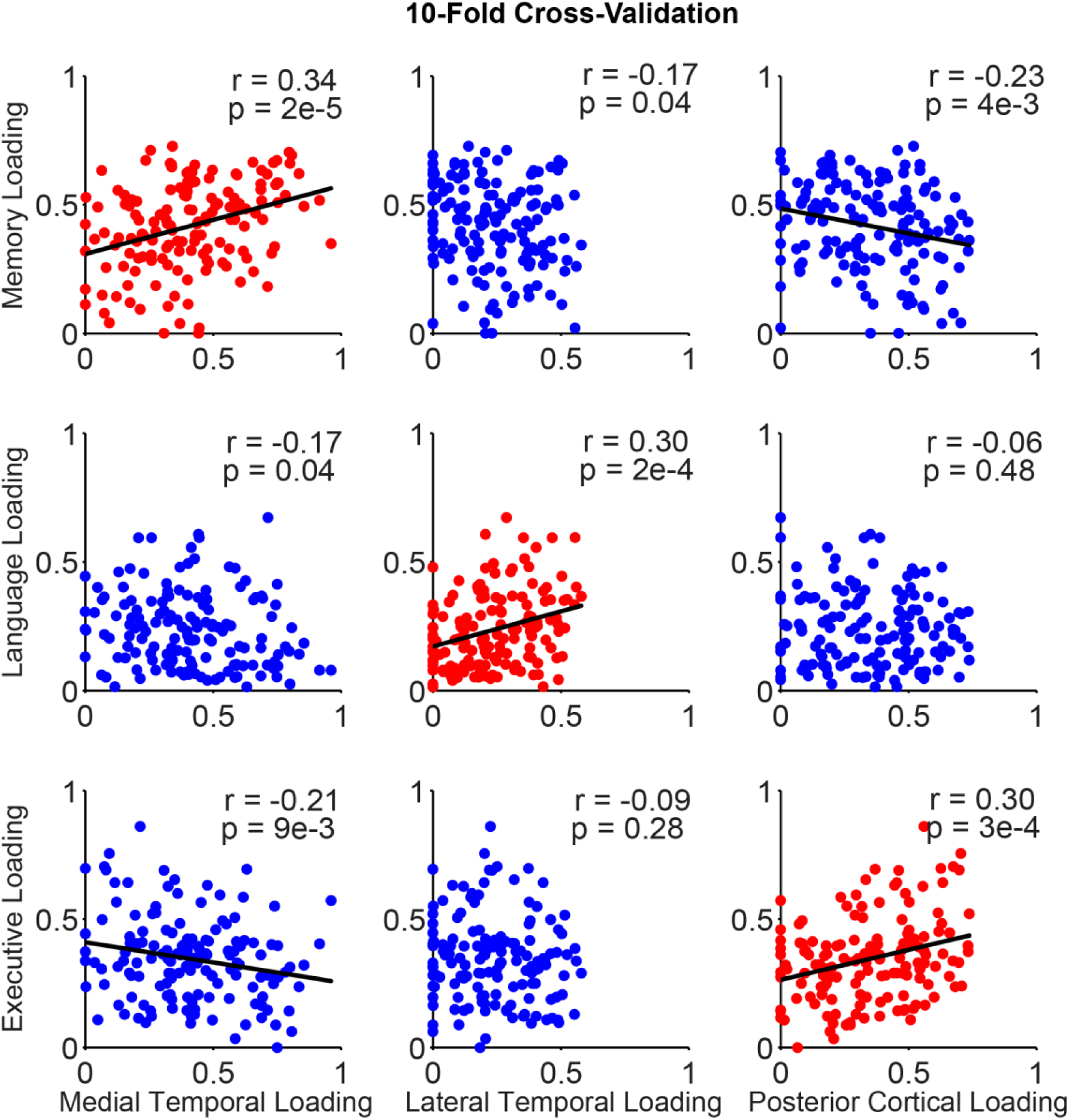
10-fold cross-validation among AD dementia patients in ADNI-GO/2. For each test fold, the multimodal Bayesian model parameters were estimated from the remaining nine training folds, and then used to infer factor loadings in the test fold subjects using only atrophy (“atrophy loading”) or only cognitive scores (“cognitive loading”). The prediction quality is measured by the correlation of atrophy and cognitive loadings across the test fold subjects, shown above for all test folds. The x axis represents atrophy loading and the y axis represents cognitive loading. Each dot represents an AD patient. Red dots indicate scatterplots for corresponding atrophy and cognitive loadings. Blue dots indicate scatterplots for non-corresponding atrophy and cognitive loadings.

#### 3.2.2 Replication in ADNI-1

We investigated whether the factors found in ADNI-GO/2 could be reproduced in an independent cohort of patients with AD from ADNI-1 (N=170). We found that the cognitive profiles and atrophy patterns in ADNI-1 (Figures S9 and S11) were similar to ADNI-GO/2 (Figures 3 and 4). Although the replication was not perfect, the discrepancies likely reflected cohort differences, rather than an artifact of our approach (see Supplementary Results S2).

#### 3.2.3 Longitudinal stability of factor loadings in AD dementia

To determine whether factor expression remained stable over time, we examined 79 dementia from ADNI-GO/2, who had both structural MRI and the full set of cognitive scores in the one-year follow-up visit. The factor loadings were highly correlated between the baseline and one-year follow up, and linear fits were close to the y = x line, suggesting that the factors were not merely reflecting disease progression (Figure 7). Interestingly, the atrophy loading (computed only using structural MRI) were more reliable (greater Pearson’s correlations) than the cognitive loading (computed only using cognitive scores).

**Figure 7.**
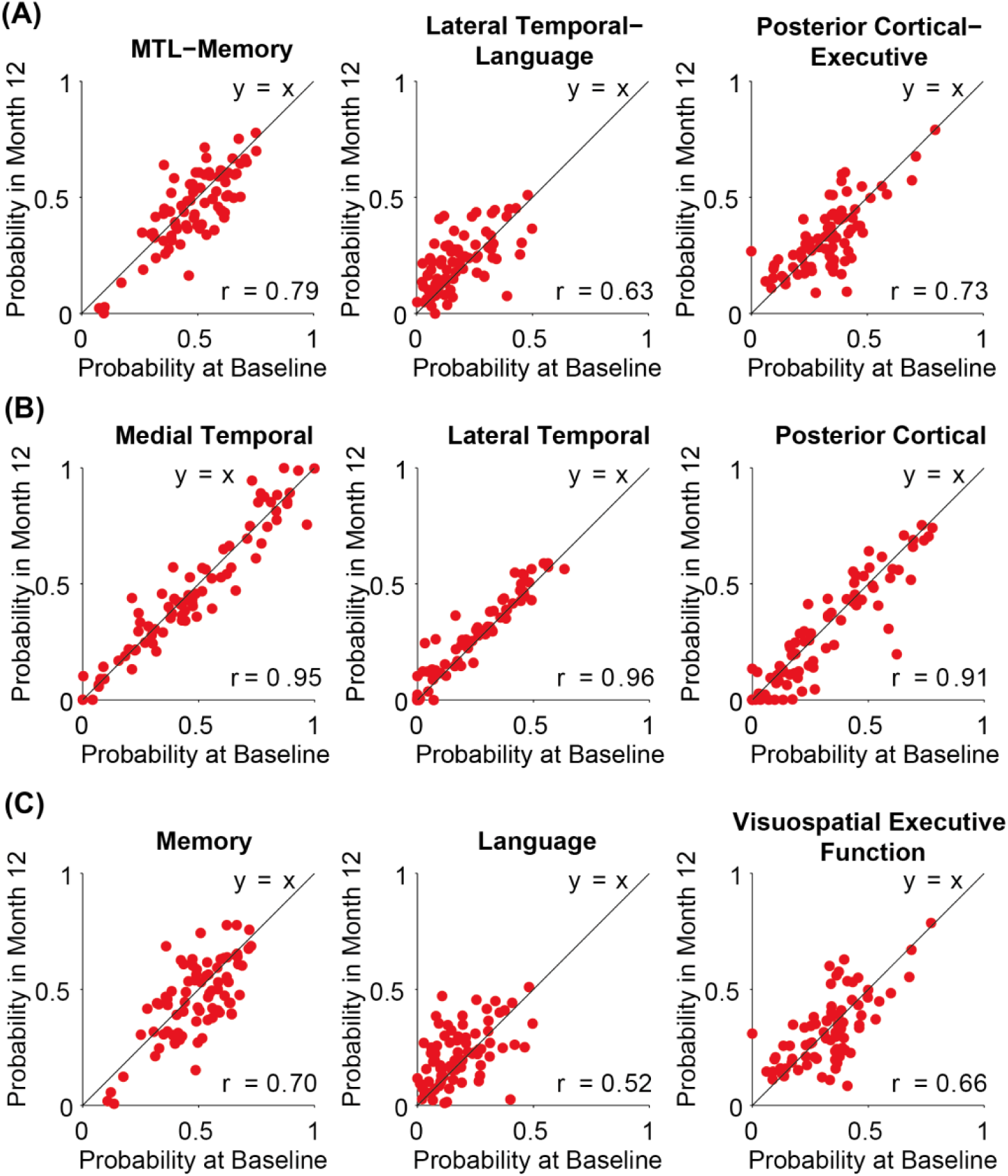
Stability of factor loadings after one year. Each red dot corresponds to an AD dementia participant. The x axis represents the probability of each factor at baseline and the y axis represents the probability of each factor after one year. (A) Atrophy-cognitive loading estimated using both structural MRI and cognitive scores. (B) Atrophy loading estimated using only structural MRI. (C) Cognitive loading estimated using cognitive scores. Factor loadings were highly correlated between the baseline and one-year follow up, and linear fits were close to the y = x line (black) suggesting that the factors were not merely reflecting disease stages.

#### 3.2.4 Comparison with Canonical Correlation Analysis (CCA)

We applied CCA to the ADNI-GO/2 data to investigate if meaningful atrophy-cognitive modes could be obtained. Permutation testing did not reveal any significant CCA mode (p > 0.13). Nevertheless, we considered the top three components and performed 10-fold cross-validation. None of the CCA mode yielded successful prediction in the out-of-sample data (Figure S13).

### 3.3 MCI Heterogeneity

#### 3.3.1 Longitudinal stability of factor loadings in Aβ+ MCI

Similar to Section 3.2.3, we examined 219 Aβ+ MCI participants from ADNI-GO/2, who had both structural MRI and the full set of cognitive scores in the one-year follow-up visit. The factor loadings were highly correlated between the baseline and one-year follow up, and linear fits were close to the y = x line, suggesting that the factors were not merely reflecting disease progression (Figure S15).

#### 3.3.2 Atrophy and cognition in Aβ+ MCI

Using the model defined in the ADNI-GO/2 participants with AD dementia, we extracted factor compositions for the 247 Aβ+ participants with MCI. Like the AD dementia participants, the majority of individuals with MCI also expressed multiple factors (Figure S14).

The model parameters estimated from the AD dementia cohort were used to infer factor loading in the Aβ+ MCI participants, using only atrophy (“atrophy loading”) or only cognitive scores (“cognitive loading”). GLMs were then utilized to investigate the relationship between atrophy loading and cognitive loading in the Aβ+ MCI participants (see Methods). We found that greater atrophy in posterior cortical regions (compared with lateral temporal cortex) was associated with worse visuospatial executive function (p = 1e-3 corrected with FDR < 0.05, Figure S16C). There was a trend that greater atrophy in posterior cortical regions (compared with MTL regions) was associated with worse visuospatial executive function (p = 0.04; Figure S16C). There was no significant association for memory and language deficits (Figure S16A-B).

#### 3.3.3 Relationship between Tau deposition and factor loading

The model parameters estimated from the AD dementia cohort were used to infer atrophy-cognitive factor loading in 76 MCI participants, using both atrophy and cognitive scores. GLMs were then utilized to investigate the relationship between the factor loading and the spatial distribution of Tau deposition (see Methods). Figure 8A shows that participants with higher Tau deposits in the medial temporal regions (compared with lateral temporal and posterior cortical regions) exhibited greater loading on the MTL-memory factor. On the other hand, participants with higher Tau signal in the lateral temporal and posterior cortical regions (compared with medial temporal regions) exhibited higher loading on the Posterior Cortical-Executive factor. However, there was no association between Tau deposits and Lateral Temporal-Language factor loadings.

**Figure 8.**
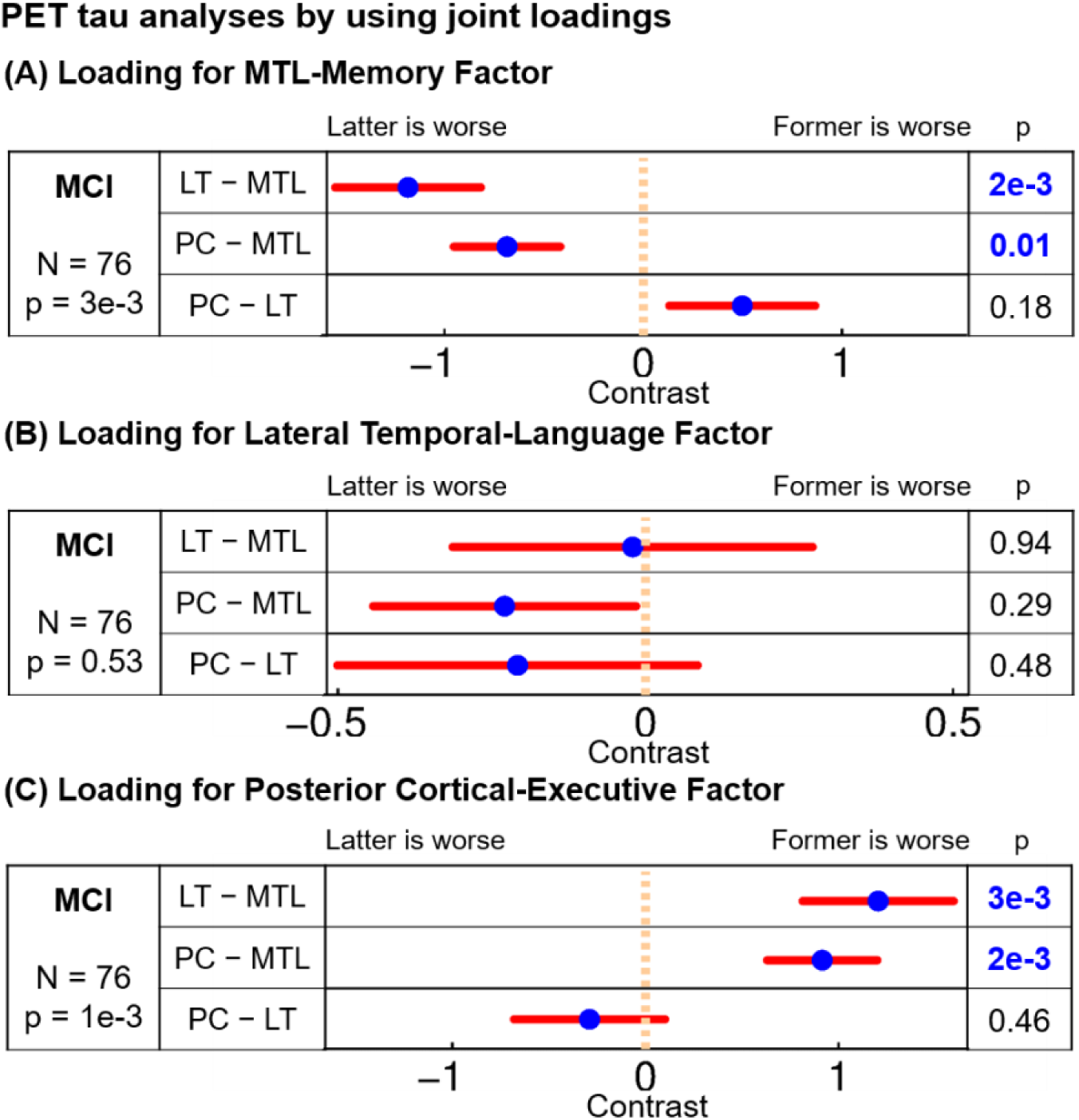
Associations between factor loadings and Tau deposits in atrophied regions among ADNI-2/3 MCI participants. All p values remaining significant after FDR (q < 0.05) are highlighted in blue. MTL, LT, and PC indicate Tau deposition in the medial temporal lobe, lateral temporal, and posterior cortical atrophied regions respectively. Blue dots are estimated differences in regression coefficients between Tau deposits in different atrophied regions and red bars show standard errors. For example, in the bottom plot, the blue dot of “PC – MTL” is on the right side (p = 2e-3), which means that participants with greater Tau deposits in the posterior cortical regions (compared with the medial temporal regions) exhibited greater factor loading on the Posterior Cortical-Executive factor.

When restricting our analysis to Aβ+ MCI participants (N = 32), the directionalities of the results were consistent with the full sample (Figure S17). However, most results were no longer significant because of the much smaller sample size. It is noteworthy that Tau signals within the factors’ masks were significantly higher in Aβ+ MCI than Aβ- MCI (p = 0.009). Thus, restricting analyses to the Aβ+ MCI group may have limited the range of Tau PET values in this analysis. For the purpose of visualization, Figure S18 shows the across-participant correlations between the atrophy-cognitive factor loadings and Tau deposition at each voxel.

## 4 Discussion

In a predominantly typical late-onset Alzheimer’s Disease cohort, we identified three latent factors, capturing (1) medial temporal lobe atrophy associated with memory and orientation (“MTL-Memory”), (2) lateral temporal lobe atrophy associated with language (“Lateral Temporal-Language”), and (3) posterior cortical atrophy associated with executive function (“Posterior Cortical-Executive”). Although the associations between the atrophy patterns and cognitive profiles were weak in MCI participants, we identified associations between some atrophy-cognitive factors and Tau PET. Specifically, greater Tau deposition in MTL regions were associated with greater loading on the MTL-Memory factor, but lower loading on the Posterior Cortical-Executive factor.

The contributions of this study are three-fold. First, we introduced a multimodal Bayesian model that allowed the joint estimation of atrophy-cognitive factors from VBM and cognitive scores simultaneously. We demonstrated that our model was better than CCA in capturing the associations between atrophy patterns and cognitive profiles. Our model might potentially be applied to other neurodegenerative or psychiatric disorders that are known to demonstrate disease heterogeneity.

Second, unlike atypical clinically-defined AD subtypes, correspondences between atrophy subtypes and cognitive domains are less clear in typical AD dementia with most studies reporting that the cortical atrophy subtype is associated with worse overall cognition and faster disease progression. An exception is our previous study, which demonstrated that the medial temporal atrophy factor was associated worse memory deficits, while the cortical atrophy factor was associated with worse executive function deficits (Zhang et al., 2016). Here, we found a novel Lateral Temporal-Language factor, which emerged only by considering atrophy and cognition jointly.

Finally, a number of studies have found strong correspondences between the spatial patterns of tau depositions and cognitive domains in cohorts mixing atypical clinically-defined AD subtypes and typical AD dementia. However, in cohorts involving predominantly typical AD or only MCI participants, the correspondences between the spatial patterns of Tau deposition and cognitive domains are less clear. Our study is therefore one of the first to demonstrate that the spatial pattern of tau heterogeneity is associated with different atrophy-cognitive factors in MCI participants.

### 4.1 Atrophy patterns

Our model revealed three factors with distinct atrophy patterns in AD patients (Figure 3). Two of these atrophy patterns were similar to results from other studies (Whitwell et al., 2012; Noh et al., 2014; Byun et al., 2015; Zhang et al., 2016; Young et al., 2018), while the third (lateral temporal) atrophy pattern was a novel insight from the current modeling approach. More specifically, the first factor was associated with preferential atrophy of the medial temporal lobe; the second factor was associated with preferential atrophy of the lateral temporal cortex; and the third factor was associated with preferential atrophy of the posterior cortex. Our medial temporal atrophy pattern was similar to the “medial temporal” subtype and our posterior cortical atrophy pattern was similar to the “parietal dominant” subtype previously reported by Noh and colleagues (Noh et al., 2014).

Besides the anatomical subtypes, our medial temporal atrophy pattern is similar to the “limbic predominant” Tau subtype, whereas the lateral temporal and posterior cortical atrophy factors is similar to the “hippocampal-sparing” Tau subtype (Murray et al., 2011; Whitwell et al., 2012). Interestingly, the “hippocampal-sparing” subtype was associated with younger age of dementia onset (Murray et al., 2011; Risacher et al., 2017), and we found a similar younger age in our posterior cortical-executive factor (Table S4). Overall, this seems to suggest that cortical presentation is associated with younger age, whereas an MTL presentation is associated with older age.

It is worth noting that Murray and colleagues likely examined a mixture of early-onset and late-onset AD participants. For example, the age of dementia onset for the hippocampal-sparing subtype was 63±10, suggesting a majority of those participants were likely early-onset AD. Consistent with Risacher et al. (2017) and our previous study (Zhang et al., 2016), our current results show that differentiation in participants’ age exists even within the limited age range found in ADNI. However, similar to our previous study (Zhang et al., 2016), we did not find an association between factor loadings and APOE ε4, while Risacher and colleagues found increased prevalence of APOE ε4 positivity in participants belonging to the hippocampal atrophy subtype (consistent with Murray et al., 2011). Since Risacher et al. (2017), Zhang et al. (2016) and our current study all utilized the ADNI database, the discrepancy is likely a result of how the subtypes were defined, rather than cohort differences.

### 4.2 Atrophy patterns and cognitive deficits

Our previous work exploring latent atrophy factors utilizing structural MRI alone identified “Temporal”, “Subcortical” and “Cortical” patterns (Zhang et al., 2016). Although post-hoc analyses revealed that the temporal atrophy factor was associated with worse memory performance and the cortical atrophy factor was associated with worse executive function performance, the subcortical atrophy factor was not associated with any behavioral performance (Zhang et al., 2016). Our current study extends our previous work by flexibly examining cognitive profiles using a data-driven approach, rather than restricting associations with pre-defined summary measures of cognitive domains.

Indeed, each cognitive test can be thought of as an imperfect measurement of some underlying cognitive construct. The test might be imperfect in that it is only measuring part of the cognitive construct or it might involve cognitive processes beyond the construct of interest, random noise and measurement error. Here, the data-driven fusion of multiple related cognitive tests seeks to better capture underlying cognitive constructs, compared with using individual imperfect cognitive tests.

In doing so, the current approach shed new insight into the link between atrophy and cognition compared to our previous work (Zhang et al., 2016), For example, the MTL-Memory factor included not just episodic memory measures (Crane et al., 2012), but also orientation to time and place, which are also well-established functions associated with the medial temporal lobe and hippocampus (O’Keefe and Dostrovsky, 1971; Squire et al., 2004; Hafting et al., 2005). Similarly, the Posterior Cortical-Executive factor included various executive function measures (e.g., “MMSE: attention” and “ADAS: number cancellation”), which were not included in the ADNI executive function composite score (Gibbons et al., 2012), but have been previously shown to related to posterior parietal regions (Crutch et al., 2012; Wu et al., 2016).

Furthermore, the current analysis identified a new Lateral Temporal-Language factor associated with lateral temporal atrophy and language deficits. Interestingly, the distribution of scores across participants (Figure 5) revealed that the range of expression of the lateral temporal factor was reduced compared to the medial temporal and posterior cortical factors, suggesting that very few participants predominately express this latent pattern. This restricted expression may explain why we did not identify this pattern when exploring MRI alone (Zhang et al., 2016). However, this factor emerged when considering joint correspondence between atrophy and cognition. Importantly, a previous study (Domoto-Reilly et al., 2012) has shown that among participants with AD dementia, deficits in the Boston naming test was associated with atrophy of the anterior temporal lobe, which overlapped significantly with our lateral temporal atrophy pattern.

In addition to implicating the lateral temporal cortex, the Lateral Temporal-Language factor was also associated with atrophy in multiple regions within the default network. A meta-analysis by Binder and colleagues has previously suggested that the left lateralized default network is involved in language processing and semantic memory (Binder et al., 2009). Traditionally, the default network has been thought to include temporal lobe, precuneus and posterior cingulate cortex, medial prefrontal cortex, dorsal prefrontal cortex and the inferior parietal cortex (Buckner et al., 2008; Yeo et al., 2011). However, previous work has also suggested that the default network can be fractionated into different functional subsystems (Laird et al., 2009; Andrews-Hanna et al., 2010; Yeo et al., 2014). Our first and second factors appeared to fractionate the default network in an interesting new way, where the medial temporal lobe was associated with the first factor, while the lateral temporal lobe and other default network regions were associated with the second factor.

Also noteworthy is that both the Lateral Temporal-Language and Posterior Cortical-Executive factors were associated with different portions of precuneus, which is consistent with the current understanding of the functional heterogeneity of this region (Margulies et al., 2009; Fornito et al., 2012). More specifically, the Posterior Cortical-Executive factor appeared to be associated with portions of precuneus that has been previously associated with the frontoparietal control network (Figure 1 of Fornito et al., 2012 and Figures 11 and 13 of Yeo et al., 2011). On the other hand, the Lateral Temporal-Language factor appeared to be associated with the precuneus/posterior cingulate core of the default network (Figure 1 of Fornito et al., 2012 and Figures 11 and 13 of Yeo et al., 2011). Thus, although default network regions (in particular the precuneus) are consistently implicated in AD, the specific pattern of regions involved might vary across individual patients.

Interestingly, focal and pronounced atrophy in these factors have been consistently implicated in atypical variants of AD. Specifically, Logopenic Aphasia (LPA) is associated with atrophy in regions found within our Lateral Temporal-Language factor whereas patients with posterior cortical atrophy (PCA) show atrophy in regions within our Posterior Cortical-Executive factor (Gorno-Tempini et al., 2011; Crutch et al., 2012). Thus, the emergence of similar patterns in the context of typical late-onset AD in the current study suggests that vulnerability in these brain networks exists along a continuum rather than being a specific feature to atypical presentations. It is possible that atypical clinical variants represent the extreme end of a continuous distribution of deficits across different cognitive domains.

### 4.3 Subtypes and stages

Due to the cross-sectional data used in this study, it is possible that our factors reflect different disease stages along the continuum of AD progression rather than heterogeneous disease subtypes (Ritchie and Touchon, 1992). However, there are two reasons why our factors likely reflected disease subtypes, rather than simply disease stages (Fonteijn et al., 2012; Young et al., 2014, 2018). First, our factors reflected the impairment of distinct cognitive domains, such as memory, language and visuospatial executive function (Figure 4). Second, one-year follow-up analyses revealed that the atrophy and cognitive loadings of the three factors were stable (Figure 7). However, we note that the one-year stability analysis is short and we cannot rule out the evolution of atrophy patterns and cognitive profiles within an individual. See further discussion in the next section.

### 4.4 Association with Tau pathology

Tau aggregations into neurofibrillary tangles are a hallmark pathological feature of Alzheimer’s disease (Braak and Braak, 1991), and are consistently shown to correlate more strongly with cognitive functions than Amyloid-β plaques (Nelson et al., 2012; Rolstad et al., 2013). Excitingly, the advent of PET ligands that bind to Tau aggregations, such as ^18^F-Flortaucipir (Chien et al., 2013), allows us to investigate the relationship between the spatial distribution of Tau *in vivo* throughout the course of Alzheimer’s disease (James et al., 2015; LaPoint et al., 2017; Lowe et al., 2017; Schöll et al., 2017; Xia et al., 2017), and has been shown to relate to clinical symptoms associated with atypical AD such as Logopenic Aphasia and Posterior Cortical Atrophy (Ossenkoppele et al., 2016).

Here, we extended the work of Ossenkoppele and colleagues by examining Tau heterogeneity in a cohort comprising predominantly of typical late-onset AD dementia participants (a population with a more limited range of clinical deficits across non-memory domains). In the context of typical late-onset AD, associations between regional ^18^F-Flortaucipir and memory (or global cognition) have been identified earlier in the “typical” AD trajectory, among participants with MCI as well as clinically normal older controls (Maass, et al., 2017a, b; Vogel et al., 2018). This is consistent with our findings that participants with MCI, who had higher Tau deposits in medial temporal and hippocampus regions, exhibited greater loadings on the MTL-Memory factor (Figure 8A). Furthermore, we extended these previous studies by showing that participants with MCI, who had higher Tau deposits in posterior cortical regions, were associated with greater loadings on the Posterior Cortical-Executive factor (Figure 8C). To the best of our knowledge, this is the first study showing that distinct atrophy-cognitive factors are associated with distinct patterns of Tau depositions in participants with MCI. However, we did not find an association between Lateral Temporal-Language factor loadings and tau PET in lateral temporal regions (Figure 8B). One might speculate that this factor should correlate with measures of language such as semantic memory. Although we did not find direct evidence for this, tests assessing language are sparse in the ADNI neuropsychological battery. It is therefore possible that the current analyses are limited in extracting a factor that robustly correlates with language performance.

Unlike the Tau analysis, the cross-sectional associations between atrophy patterns and cognitive profiles were weaker in MCI participants compared with AD dementia participants in both the current study (Figure S16) and our previous study (Zhang et al., 2016). Furthermore, while we could reliably estimate very similar atrophy factors in MCI and AD dementia participants independently (Zhang et al., 2016), we could not reliably estimate atrophy-cognitive factors when restricted to MCI participants (results not shown). It is possible that the coupling between the atrophy patterns and cognitive profiles might emerge later in the disease process, which would explain the weak correlations among MCI. Future studies that address the association between atrophy factors and longitudinal cognition during the stages preceding overt dementia may provide insight into this possibility, and provide more direct insights into the time course of these changes. This is consistent with the perspective that Tau-mediated injury is thought to occur earlier in the disease process, followed by atrophy and cognitive decline (Jack et al., 2010; 2013).

### 4.5 Limitations

Our study has several limitations. First, our Tau analysis utilized a relatively small sample of 76 MCI participants. Second, to maximize sample size, our Tau analysis included all MCI participants with Tau PET data, regardless of amyloid status. Our control analyses using a smaller sample of Aβ+ MCI participants showed results consistent with the full sample, but were generally not statistically significant. Consequently, this affects the interpretability of our results.

We note that a recent paper demonstrated that the spatial patterns of tau deposits were associated with different cognitive domains in a sample comprising 25 Aβ+ MCI and 48 Aβ+ typical late-onset AD dementia participants (Ossenkopple et al., 2019). By restricting the sample to Aβ+ participants, the study provided a cleaner link to typical AD. However, by mixing both MCI and AD dementia participants, the temporal staging of the observations became less clear. Indeed, when the analysis was restricted to the 25 Aβ+ MCI participants, the results were no longer significant (Ossenkopple et al., 2019), similar to our control analysis of Aβ+ MCI participants (N = 32). Therefore, our study and Ossenkopple et al. (2019) together suggest that a minimum sample size of about 70 Aβ+ MCI participants might be necessary to demonstrate clear relationships between Tau deposition patterns and cognitive domains during the MCI stage of typical AD.

Finally, we note that higher tau deposition in the hippocampus may be affected by the known off-target binding of ^18^F-flortaucipir to the choroid plexus (Lee, et al., 2018). Future studies should apply these approaches to second-generation Tau tracers (e.g., Lois, et al., 2018) that may show reduced choroid plexus binding.

### 4.6 Future work

Study-specific VBM templates for both ADNI-1 and ADNI-GO/2 have been made publicly available, so other researchers can potentially utilize the templates to estimate factor loadings in their own participants. However, further studies are necessary to determine if the application of ADNI templates to participants outside of ADNI might result in sub-optimal results or biases.

The code for the MMLDA model has also been made publicly available. The model can potentially be applied to other heterogeneous neurodegenerative disorders, such as Parkinson’s disease. Furthermore, even though we have only utilized the model for two modalities, the model can be easily extended to handle more than two modalities, e.g., atrophy, cognitive scores and Tau. However, this requires sufficient number of participants with all modalities.

## 5 Conclusion

By utilizing the proposed multi-modal latent Dirichlet allocation (MMLDA) model, our study revealed three latent AD factors with distinct atrophy patterns and corresponding profiles of cognitive deficits. The first factor was associated with medial temporal atrophy and deficits in memory and orientation; the second factor was associated with lateral temporal atrophy and language deficits; the third factor was associated with posterior cortical atrophy and visuospatial executive function deficits. Our approach allowed each individual to express multiple factors to various degrees, rather than assigning the individual to a single subtype. This is biologically more plausible than non-overlapping subtypes, given that multiple non-mutually exclusive factors likely influence this heterogeneity (such as age, co-morbid pathologies, genetics, exposures throughout the lifespan, etc). Therefore, each participant exhibited his or her own unique factor composition, which might potentially be exploited to predict individual-specific cognitive profile that may improve disease monitoring. Finally, our study suggested that these atrophy-cognitive profiles were associated with distinct patterns of Tau deposition in mild cognitively impaired participants, highlighting the emergence of these subtypes early in the AD trajectory.

## 6 Acknowledgements

We are currently supported by Singapore MOE Tier 2 (MOE2014-T2-2-016), NUS Strategic Research (DPRT/944/09/14), NUS SOM Aspiration Fund (R185000271720), Singapore NMRC (CBRG/0088/2015), NUS YIA and the Singapore National Research Foundation (NRF) Fellowship (Class of 2017). Our research also utilized resources provided by the Center for Functional Neuroimaging Technologies, P41EB015896 and instruments supported by 1S10RR023401, 1S10RR019307, and 1S10RR023043 from the Athinoula A. Martinos Center for Biomedical Imaging at the Massachusetts General Hospital. Our computational work was partially performed on resources of the National Supercomputing Centre, Singapore (https://www.nscc.sg). Data collection and sharing for this project was funded by the Alzheimer’s Disease Neuroimaging Initiative (ADNI) (National Institutes of Health Grant U01 AG024904) and DOD ADNI (Department of Defense award number W81XWH-12-2-0012). ADNI is funded by the National Institute on Aging, the National Institute of Biomedical Imaging and Bioengineering, and through generous contributions from the following: AbbVie, Alzheimer’s Association; Alzheimer’s Drug Discovery Foundation; Araclon Biotech; BioClinica, Inc.; Biogen; Bristol-Myers Squibb Company; CereSpir, Inc.; Cogstate; Eisai Inc.; Elan Pharmaceuticals, Inc.; Eli Lilly and Company; EuroImmun; F. Hoffmann-La Roche Ltd and its affiliated company Genentech, Inc.; Fujirebio; GE Healthcare; IXICO Ltd.; Janssen Alzheimer Immunotherapy Research & Development, LLC.; Johnson & Johnson Pharmaceutical Research & Development LLC.; Lumosity; Lundbeck; Merck & Co., Inc.; Meso Scale Diagnostics, LLC.; NeuroRx Research; Neurotrack Technologies; Novartis Pharmaceuticals Corporation; Pfizer Inc.; Piramal Imaging; Servier; Takeda Pharmaceutical Company; and Transition Therapeutics. The Canadian Institutes of Health Research is providing funds to support ADNI clinical sites in Canada. Private sector contributions are facilitated by the Foundation for the National Institutes of Health (www.fnih.org). The grantee organization is the Northern California Institute for Research and Education, and the study is coordinated by the Alzheimer’s Therapeutic Research Institute at the University of Southern California. ADNI data are disseminated by the Laboratory for Neuro Imaging at the University of Southern California.

## Footnotes

† Unfortunately, there were only 21 AD dementia participants with Tau PET, so we did not perform an analysis on this subgroup because of the small sample size.

