## Supplementary Material for "Multi-modal Latent Factor Exploration of Atrophy, Cognitive and Tau Heterogeneity in Alzheimer’s Disease"

This supplementary material is divided into **Supplementary Methods, Supplementary Results, Supplementary Figures and Tables.**

#### Supplementary Methods

This section provides additional information about processing, mathematical modeling and statistical procedures. Section S1 provides more details for structural MRI and cognitive scores processing. Section S2 provides mathematical details about the extension of latent dirichlet allocation (LDA) from unimodal to multimodal data. Section S3 explains how the multimodal Bayesian model parameters are estimated. Section S4 explains how structural MRI and cognitive scores can be mapped as inputs to the multimodal Bayesian model. Section S5 provides more information about the setup and statistical tests for general linear modeling (GLM) and logistic regression of patient characteristics. Section S6 provides more information about the setup for CCA. Section S7 provides more information about the setup and statistical tests for general linear modeling (GLM) of atrophy and cognitive loadings. Finally, section S8 provides more information about the setup and statistical tests for general linear modeling (GLM) of tau deposition with atrophy-cognitive factor loading. Section S9 explains how the amyloid status of ADNI cohorts are defined.

##### S1. Structural MRI & cognitive scores processing

The structural MRI (T1-weighted) were analyzed using the CAT12 VBM software. Briefly, the T1 images were first normalized to a template space and segmented into GM, white matter and cerebrospinal fluid. Second, the gray and white matter images were used to create a study-specific template. Third, the template was transformed to the MNI standard space. Fourth, the T1 images were renormalized to the study-specific template (in MNI space) and re-segmented into GM density maps. The resulting GM density images were downsampled from 1.5mm to 2mm and smoothed with a Gaussian kernel of 10-mm FWHM. The 10-mm FWHM smoothing was chosen to be consistent with our previous work (Zhang et al., 2016) and was within the typical range of 8mm to 12mm FWHM smoothing used in VBM. The downsampling reduced computational time, but did not substantially affect the resulting factors. Using the CAT12 default 8-mm FWHM smoothing (http://www.neuro.uni-jena.de/cat12/CAT12-Manual.pdf) without downsampling yielded very similar factors: mean correlation between factors with 8-mm (without downsampling) and 10-mm smoothing (with downsampling) was r = 0.95.

Having obtained the GM density maps, we first applied $\log_{10}$ transformation to each voxel of the maps. For each voxel, we then regressed the effects of age, sex and intracranial volume (ICV) from all participants. The regression coefficients were estimated from CN participants to retain any interaction between AD and participants' characteristics (e.g., interaction between AD and age). More specifically, for each voxel, we estimated $\boldsymbol{\beta}_{CN}$ from CN participants via the following GLM: $voxel\_value_{CN} = \boldsymbol{\beta}_{CN} \times (1 + age_{CN} + sex_{CN} + ICV_{CN}) + \epsilon$. Once we have estimated $\boldsymbol{\beta}_{CN}$, we then regressed the effects of age, sex and ICV from all participants (CN, MCI, AD) with the following $residual_{ALL} = voxel\_value_{ALL} - \boldsymbol{\beta}_{CN} \times (1 + age_{ALL} + sex_{ALL} + ICV_{ALL})$.

The same procedure was applied to the cognitive scores. More specifically, for each cognitive score, we estimated $\boldsymbol{\beta}_{CN}$ from CN participants via the following GLM: $cognitive\_score_{CN} = \boldsymbol{\beta}_{CN} \times (1 + age_{CN} + sex_{CN} + ICV_{CN}) + \epsilon$. Once we have estimated $\boldsymbol{\beta}_{CN}$, we then regressed the effects of age, sex and ICV from all participants (CN, MCI, AD) with the following $residual_{ALL} = cognitive\_score_{ALL} - \boldsymbol{\beta}_{CN} \times (1 + age_{ALL} + sex_{ALL} + ICV_{ALL})$.

To further illustrate the goal of this regression procedure, suppose we find that across CN participants, there is a drop of 0.1 unit of test score per year *cross-sectionally*. Among AD dementia participants, there is a drop of 0.3 unit of test score per year *cross-sectionally*. After computing the regression coefficients based on CN participants and apply the regression coefficients to AD dementia participants, we might find that among AD participants, there is a drop of 0.2 unit of test score per year cross-sectionally. Thus, the (linear) effects of "healthy aging" have been removed, but the effects of "unhealthy aging" remains.

S2. Extension of LDA to multimodal data.

The LDA model was originally developed to automatically discover latent topics from a corpus of text documents. The model assumes that each document is an unordered collection of words associated with a subset of *K* latent topics. Each topic is represented by a probability distribution over a dictionary of words. Given a collection of documents, there exist algorithms to estimate the probability of a dictionary word given a topic Pr(Word | Topic) and the probability that a topic is associated with a particular document Pr(Topic | Document). The LDA model is useful, because it allows a document to be associated with multiple topics (which can be shared across documents) and each topic to be associated with multiple words (which can be shared across topics). In our previous work, we applied LDA to discover atrophy factors (Zhang et al., 2016). To achieve this, one can think of AD patients as text documents, atrophy factors as topics, and MNI152 voxels as dictionary words. Correspondingly, each patient expresses one or more latent atrophy factors to different extents [Pr(Factor | Patient)], and each factor is associated with atrophy at multiple voxels to different extents [Pr(Voxel | Factor)].

Here, an extension the LDA model was utilized to incorporate two modalities (MRI gray matter atrophy and neuropsychological testing scores), herein referred to as multi-modality LDA (MMLDA). Different extensions of LDA have been proposed to handle multiple modalities (Barnard et al., 2003; Blei and Jordan, 2003). Our extension is basically the same as that of Putthividhya and colleagues (Putthividhya et al., 2007), although the following exposition has been devised to (hopefully) be more intuitive for the likely readers of this work. To this end, we discuss the extension of LDA to multimodal data by first staying within the text analysis domain.

More specifically, to extend LDA to handle multiple modalities, we can think of each text document being written in multiple languages with imperfect translations, so that the information contents are not identical across languages. We can think of each language as one modality and for simplicity, we will only assume two languages (or modality), e.g., English and Spanish. Given a corpus of text documents written in two languages (English and Mandarin) and for a fixed number of topics $K$, multimodal LDA seeks to estimate the probability a text document involves a topic ($\theta$, i.e., Pr(topic | document)), the probability that a topic is associated with an English word ($\beta^1$, i.e., Pr(English word | Topic)) and the probability that a topic is associated a Spanish word ($\beta^2$, i.e., Pr(Spanish word | Topic)).

Suppose there are $K$ topics and $D$ documents. Then $\theta$ is a $D \times K$ matrix, where each row sums to one. $\theta_d$ is the $d$-th row of the $\theta$ matrix and the $k$-th element indicates the probability that the $d$-th document involves the $k$-th topic. Suppose there are $V$ words in the English vocabulary. Then $\beta^1$ is a $K \times V$ matrix, where each row sums to one. $\beta_k^1$ is the $k$-th row of the $\beta^1$ matrix and the $v$-th element ($\beta_{k,v}^1$) indicates the probability that the $k$-th topic is associated with atrophy at the $v$-th English word. On the other hand, suppose there are $C$ words in the Spanish dictionary. Then $\beta^2$ is a $K \times C$ matrix, where each row sums to one. $\beta_k^1$ is the $k$-th row of the $\beta^1$ matrix and the $c$-th element ($\beta_{k,c}^1$) indicates the probability that the $k$-th topic is associated with the $c$-th Spanish word.

The model (Fig. S13) assumed a prior (Dirichlet($\alpha$)) on the topic composition ($\theta_d$) for each document. Given the topic composition ($\theta_d$), the total word count ($N_d^1$) associated with the English translation of the $d$-th document, and the total word count ($N_d^2$) associated with the Spanish translation of the $d$-th document, the two translations of the $d$-th document are generated as follows.

1. Each word in the English translated document is generated independently. For the $n$-th English word
   a. Sample a topic $z_{d,n}^1$ using $\theta_d$ ($d$-th row of the $\theta$ matrix)
   b. Sample the word $w_{d,n}^1$ using the distribution specified in the $z_{d,n}^1$-th row of the $\beta^1$ matrix.
2. Each word in the Spanish translated document is generated independently. For the $n$-th Spanish word
   a. Sample a topic $z_{d,n}^2$ using $\theta_d$ ($d$-th row of the $\theta$ matrix)
   b. Sample the word $w_{d,n}^2$ using the distribution specified in the $z_{d,n}^2$-th row of the $\beta^2$ matrix.

The above generative process specifies the mathematical model exactly. The following section will describe how the model parameters are estimated.

S3. Estimation and inference of the multimodal Bayesian model

Given the text documents translated into both English and Spanish, and a pre-defined number of topics $K$, the goal is to estimate the probability that a text document involves a topic ($\theta$, i.e., Pr(topic | document)), the probability that a topic is associated with an English word ($\beta^1$, i.e., Pr(English word | Topic)) and the probability that a topic is associated a Spanish word ($\beta^2$, i.e., Pr(Spanish word | Topic)). Like the original LDA paper (Blei et al., 2003), we utilized the variational expectation-maximization (VEM) algorithm to estimate the parameters.

The algorithm involves iterating between the variational E-step and M-step until convergence. In the E-step, $\alpha, \beta^1, \beta^2$ are assumed fixed and the following variational parameters are updated by iterating the following two equations until convergence.

$$\phi_{d,n,k}^l \propto \beta_{k,v(n_{ld})}^l exp\left(\psi(\gamma_{d,k}) - \psi\left(\sum_{i=1}^{K} \gamma_{d,i}\right)\right) \quad (1)$$

$$\gamma_{d,k} = \alpha + \sum_{l=1}^{2}\sum_{n=1}^{N_d^l} \phi_{d,n,k}^{l}\, , \ (2)$$

where $\phi$ is the variational parameter for $z$, so we can interpret $\phi_{d,n,k}^1$ as the posterior probability that the $n$-th word of the English translation of the $d$-th document belonged to the $k$-th topic. Similarly, we can interpret $\phi_{d,n,k}^2$ as the posterior probability that the $n$-th word of the Spanish translation of the $d$-th document belonged to the $k$-th topic. $\gamma$ is the variational parameter for $\theta$, so we can interpret $\gamma_{d,k}$ as the posterior probability of the $d$-th document exhibiting the $k$-th topic. $v(n_{1d})$ indexes the word (in the English dictionary) that the n-th word of the English translation of the $d$-th document corresponded to, while $v(n_{2d})$ indexes the word (in the Spanish dictionary) that the n-th word of the Spanish translation of the $d$-th document corresponded to. $\psi$ is the digamma function (the first derivative of the log Gamma function).

In the M-step, $\phi$ and $\gamma$ are assumed fixed. The hyperparameter $\alpha$ is updated using the Newton-Raphson algorithm (Blei et al., 2003) and $\beta$ is updated as follows

$$\beta_{k,v}^{l} \propto \sum_{d=1}^{D}\sum_{n=1}^{N_d^l} \phi_{d,n,k}^{l}\, w_{d,n}^{l,v} \quad (3)$$

where $w_{d,n}^{1,v}$ is 1 if the $n$-th English word in the $d$-th document corresponded to the $v$-th dictionary word and 0 otherwise. $w_{d,n}^{2,v}$ is 1 if the $n$-th Spanish word in the $d$-th document corresponded to the $v$-th dictionary word and 0 otherwise. Looking at Eq. (1) to Eq. (3), we observe that each document can be summarized by the number of times each English word occurs and the number of times each Spanish word occurs.

Suppose we are given a new document that has been translated into both English and Spanish. Assuming that we have already estimated $\alpha, \beta^1, \beta^2$, then we can simply apply the E-step (Eq. (1) and Eq. (2)) till convergence to estimate the topic composition of the new document.

We can also infer the topic composition of a new document based on a single language (modality). This can be accomplished by applying the E-step with the following slight modification:

$$\gamma_{d,k}^{l} = \alpha + \sum_{n=1}^{N_d^l} \phi_{d,n,k}^{l}. \quad (4)$$

Comparing Eq. (2) and Eq. (4), we note that Eq. (2) summed over both languages, while Eq. (4) does not. Upon convergence, we can interpret $\gamma_{d,k}^1$ as the topic loading of the document based only on the English translation, and $\gamma_{d,k}^2$ as the topic loading of the document based only on the Spanish translation.

S4. Application of Bayesian model to structural MRI and cognitive scores

To map the multimodal LDA to VBM and cognitive scores, we can think of English dictionary words as corresponding to MNI voxels and Spanish dictionary words as correspond to the cognitive scores. Furthermore, multimodal LDA assumes that a document is summarized by the number of times each English dictionary word appears in the English translation and the number of times each Spanish dictionary word appears in the Spanish translation. Because word count is discrete and non-negative, so the log-transformed GM density images and cognitive scores have to be thresholded and discretized.

More specifically, for each voxel of the log-transformed GM density images, z transformation (using the mean of CN participants and variance of all participants) was applied. The resulting z-scores were flipped in sign, so that a positive z-score at a particular voxel of an individual would indicate atrophy at the voxel relative to CN participants. The same z-normalization procedure was applied to cognitive scores, such that a positive cognitive z-score would indicate poorer performance than CN participants.

Finally, the atrophy and cognitive z-scores were thresholded and discretized. If the threshold was set at zero (negative values set to zero), too much statistical power would be lost in the case of the cognitive scores (Fig. S14). Therefore, we added an offset of one to all z-scores before thresholding at zero. Varying this offset (e.g., adding two instead of one) did not significantly modify the results.

Furthermore, there were hundreds of thousands of voxels, but only 28 cognitive scores. To balance the two modalities, the cognitive z-scores were multiplied by the ratio between number of voxels and number of cognitive scores. Finally, both atrophy and cognitive z scores were multiplied by 10 and rounded to the nearest integer.

To summarize, for each participant, the brain atrophy pattern is represented by a non-negative integer at each voxel and the cognitive deficit profile is represented by a non-negative integer for each cognitive test. A more positive integer at a voxel would indicate greater atrophy, while a more positive integer for a cognitive score would indicate worse cognitive performance. The correspondence among various terminologies (e.g. atrophy pattern), description (e.g. the probability that a factor is associated with a voxel [Pr(Voxel | Factor)]) and Bayesian model symbol (e.g. $\beta^1$) can be found in Table S7.

To estimate the model parameters, the VEM algorithm was run with 20 different random initializations and the result with highest log likelihood was chosen as the final estimate.

#### S5. GLM and logistic regression for patient characteristics

We apply GLM to continuous patient characteristics (e.g., age) and logistic regression to binary patient characteristics (e.g., sex) to evaluate their relationships with factor loadings. To illustrate the GLM, let's denote age by $y$, factor 1 loading by $f_1$, factor 2 loading by $f_2$, and factor 3 loading by $f_3$. Then the GLM was $y = \beta_0 + \beta_2 f_2 + \beta_3 f_3 + \epsilon$, where $\beta_0, \beta_2, \beta_3$ denote regression coefficients and $\epsilon$ is the residual. Factor 1 loading $f_1$ was implicitly modeled because $f_1 + f_2 + f_3 = 1$.

Statistical tests involved null hypotheses of the form $H\beta = 0$, where $\beta = [\beta_0,\ \beta_2,\ \beta_3]^T$, and $H$ is the linear contrast. First, we performed an omnibus test of overall differences across all factor loadings with $H = [0, 1, 0; 0, 0, 1]$. We then tested for pairwise differences between factor loadings. For example, $H = [0, 1, 0]$ tested the difference between factor 2 and factor 1 loadings; $H = [0, 0, 1, 0, 0]$ tested the difference between factor 3 and factor 1 loadings; $H = [0, -1, 1, 0, 0]$ compared factor 3 and factor 2 loadings.

To illustrate the logistic regression, let's denote sex by $y$, factor 1 loading by $f_1$, factor 2 loading by $f_2$, and factor 3 loading by $f_3$. The regression model was $\log(\mu/(1-\mu)) = \beta_0 + \beta_2 f_2 + \beta_3 f_3 + \epsilon$, where $\mu$ is the probability of female, $\beta_0, \beta_2, \beta_3$ denote regression coefficients and $\epsilon$ is the residual. Intuitively, the linear combination $\beta_0 + \beta_2 f_2 + \beta_3 f_3$ predicts the probability of female ($y = 1$); $\exp(\beta_0)$ reflects the odds ratio for factor 1, $\exp(\beta_2)$ reflects the ratio of odds ratio between factors 3 and 1, and $\exp(\beta_3)$ reflects the ratio of odds ratio between factors 2 and 1.

The likelihood ratio test was utilized to determine whether the probabilities of sex were different across factors. More specifically, the test involved comparing the likelihood of an appropriate restricted model to the original model. We first performed a statistical test of overall differences across factors. In this case, the restricted model $\log\big(\mu/(1-\mu)\big) = \beta_0 + \epsilon$ was fitted to the data, and the resulting likelihood was compared with that of the original model. We then tested for differences between the factors. For example, to compare factor 2 and factor 3 loadings, the restricted model was specified as $\log\big(\mu/(1-\mu)\big) = \beta_0 + \beta_{23}(f_2 + f_3) + \epsilon$ because $\beta_2 = \beta_3$ under the null hypothesis. To compare factor 1 and 2 loadings, the restricted model was $\log\big(\mu/(1-\mu)\big) = \beta_0 + \beta_3 f_3 + \epsilon$, because $\beta_2 = 0$ under the null hypothesis.

S6. Comparisons with canonical correlation analysis (CCA)

To apply CCA to the 149 ADNI-GO/2 participants, for each voxel of the log-transformed GM density images, z transformation (using the mean of CN participants and variance of all participants) was applied. The same procedure was applied to the cognitive scores. To avoid overfitting the CCA, we followed Smith and colleagues' (Smith et al., 2015) procedure of applying PCA to reduce the dimensionality of both atrophy and cognitive scores, retaining 95% of the variance. Varying the number of PCA components did not significantly affect the results. Statistical significance of the CCA modes was established using permutation testing (Smith et al., 2015). In addition, we also performed 10-fold cross-validation, just like in MMLDA.

S7. GLM of atrophy and cognitive loadings in Aβ+ MCI participants

To evaluate the associations between atrophy patterns and cognitive deficits in MCI participants, we applied the GLM for each cognitive loading. For example, let's denote memory loading by $h$, medial temporal loading by $m$, lateral temporal loading by $l$, and posterior cortical loading by c. Age $x_1$ and sex $x_2$ were included as nuisance variables, so the final GLM was $\mathrm{h} = \beta_0 + \beta_l l + \beta_c c + \beta_1 x_1 + \beta_2 x_2 + \epsilon$, where β's denote the regression coefficients, and $\epsilon$ is the residual. The medial temporal loading $m$ was implicitly modeled because $m + l + c = 1$.

Similar to the previous section, statistical tests involved null hypotheses of the form $H\beta = 0$, where $\beta = [\beta_0, \beta_l, \beta_c, \beta_1, \beta_2]^T$, and $H$ is the linear contrast. First, we performed an test of overall differences across all atrophy factor loadings with $H = [0,1,0,0,0; 0,0,1,0,0]$. We then tested for pairwise differences between atrophy loadings. For example, $H = [0,1,0,0,0]$ tested the differences between lateral temporal and medial temporal atrophy loadings; $H = [0,0,1,0,0]$ compared posterior cortical and medial temporal atrophy loadings; $H = [0,-1,1,0,0]$ compared posterior cortical and lateral temporal atrophy loadings.

S8. GLM of tau deposition and factor loading

To evaluate the association between tau deposits and factor loadings, we applied the GLM for each factor loading. For example, let's denote MTL-Memory loading by $h$, medial temporal tau deposition $\tau_m$, lateral temporal tau deposition $\tau_l$ and posterior cortical tau deposition $\tau_c$. The nuisance variables included age $x_1$ and sex $x_2$, which was obtained by averaging tau SUVR within the gray matter of each subject. Hence, the GLM was $h = \beta_0 + \beta_m \tau_m + \beta_l \tau_l + \beta_c \tau_c + \beta_1 x_1 + \beta_2 x_2 + \epsilon$, where β's denote the regression coefficients, and $\epsilon$ is the residual. In this case, we note that $\tau_m + \tau_l + \tau_c$ is not necessarily a constant, so all three tau deposits are included in the GLM.

Statistical tests included null hypotheses of the form $H\beta = 0$, where $\beta = [\beta_0, \beta_m, \beta_l, \beta_c, \beta_1, \beta_2]^T$, and $H$ is the linear contrast. First, we performed an omnibus test of overall differences across all tau deposits with $H = [0, -1, 1, 0, 0, 0; 0, -1, 0, 1, 0, 0]$. We then tested for pairwise differences between tau deposits. For example, $H = [0, -1, 1, 0, 0, 0]$ tested the differences between lateral temporal tau and medial temporal tau deposits; $H = [0, -1, 0, 1, 0, 0]$ compared posterior cortical tau and medial temporal tau deposits; $H = [0, 0, -1, 1, 0, 0]$ compared posterior cortical tau and lateral temporal tau deposits.

S9. Amyloid Status

To generate the amyloid statistics in Table 1 for ADNI-GO/2 baseline, and ADNI-GO/2 month 12 cohorts, participants were categorized as abnormal (+) or normal (-) on florbetapir using a cortical retention ratio cutoff value of 1.11 (Landau et al., 2012). This value is equivalent to the upper 95% confidence interval above the mean of a group of young normal controls (Landau et al., 2013). In the case of ADNI-1 baseline, since there were no florbetapir PET imaging available, participants were categorized as abnormal (+) if the CSF amyloid concentration was less than 192 pg/mL (Shaw et al., 2009). In the case of ADNI-2/3 Tau PET cohort, we utilized CSF amyloid status with dates before and closest to the Tau PET scan date. If the participants did not have CSF amyloid status, we used florbetapir PET amyloid status with dates before and closest to the Tau PET scan date.

### Supplementary Results

S1. 10-fold cross-validation for the four-factor model

In section 3.2.1, we showed that the 10-fold cross-validation results for the three-factor model were significant for all factors. When the 10-fold cross-validation was repeated for the four-factor solution, the first three atrophy patterns (which corresponded to the three-factor solution) successfully predicted individuals' cognitive deficit profiles with r = 0.31, 0.29 and 0.35 respectively. However, the fourth atrophy pattern failed to predict the corresponding cognitive deficit profile (r = 0.09). This further motivates our decision to focus on the three-factor model in this paper.

S2. Replication in ADNI-1

In section 3.2.2, we showed that atrophy patterns in ADNI-1 were similar to ADNI-GO/2. To quantify the similarity between ADNI-1 and ADNI-GO/2, we computed correlations between corresponding atrophy patterns from the two cohorts, which were 0.83, 0.65 and 0.63 for factors 1, 2 and 3 respectively.

The profiles of cognition deficits were also similar between ADNI-GO/2 and ADNI-1 (Figure S11 compared to Figure 4). For example, the top 6 cognitive scores for the MTL-Memory factor in ADNI-GO/2 were among the top 7 cognitive scores (with different ordering) for the corresponding factor in ADNI-1. Similarly, the top 5 cognitive scores for the Lateral Temporal-Language factor in ADNI-GO/2 were among the top 6 cognitive scores (with different ordering) for the corresponding factor in ADNI-1. Correlations between corresponding cognitive profiles from the two cohorts were 0.98, 0.58 and 0.83. These results suggest that although all factors were replicable, the medial temporal lobe - memory (MTL-M) factor was the most replicable, which might not be surprising given that we are focusing on AD dementia in this study.

Interestingly, the correlations of cognitive profiles between the two cohorts were generally stronger than that of atrophy patterns. One major difference between ADNI-GO/2 and ADNI-1 is that the lateral temporal factor was left lateralized in ADNI-GO/2, but more bilateral in ADNI-1. To ensure that discrepancy in the lateralization was not due to artifacts of the MMLDA algorithm, for each participant, we compute composite memory, language and visuospatial executive function scores by simply averaging the top five cognitive scores associated with each factor (Figure 4). This overall score was then correlated against voxelwise VBM data across the ADNI-1 and ADNI-GO/2 participants.

The resulting correlation maps (Figure S12) were consistent with the atrophy patterns (Figures S2 and S10). Of particular importance is that the language composite score was correlated with atrophy in only the left temporal lobe in ADNI-GO/2, but correlated with atrophy in both left and right temporal lobes in ADNI-1. Furthermore, correlations between the ADNI-1 and ADNI-GO/2 correlation maps (Figure S12) were 0.31 and 0.39 for the first two correlation maps, which were significantly lower than the replicability of the atrophy patterns (0.83 and 0.65). For the third correlation maps, correlation between ADNI-1 and ADNI-GO/2 was 0.68, which was slightly better than the atrophy pattern' replicability (0.63).

Overall, this suggests that discrepancies between ADNI-1 and ADNI-GO/2 were not an artifact of the MMLDA algorithm, but inevitable cohort differences (e.g., scanner upgrade, etc) in a long-term study like ADNI.

#### Supplementary Figures and Tables

**Table S1. Chosen cognitive scores from ADAS, MMSE and NEUROBAT spreadsheets.** The left column lists the behavioral description and the right column corresponds to the fieldname of the behavior in the spreadsheet. For the MMSE, the blocks show how we merge the item scores into subscores. For example, “MMSE: Orientation to time” was obtained by summing “MMDATE”, “MMYEAR”, “MMMONTH”, “MMDAY” and "MMSEASON”. ADAS = Alzheimer’s Disease Assessment Scale; MMSE = Mini-Mental State Examination; NEUROBAT = Neuropsychological Battery; BNT = Boston Naming Test; CFT = Category Fluency Test; LM = Logical Memory; RAVLT = Rey Auditory Verbal Learning Test; TMT = Trail Making Test.

| **Behavioral Description** | **Fieldname** | **Behavioral Description** | **Fieldname** |
|---|---|---|---|
| **ADAS** | | **MMSE** | |
| ADAS: Word Recall | Q1SCORE | MMSE: Orientation to time | MMDATE |
| ADAS: Commands | Q2SCORE | | MMYEAR |
| ADAS: Construction | Q3SCORE | | MMMONTH |
| ADAS: Delayed Word Recall | Q4SCORE | | MMDAY |
| ADAS: Naming | Q5SCORE | | MMSEASON |
| ADAS: Ideational Praxis | Q6SCORE | MMSE: Orientation to place | MMHOSPIT |
| ADAS: Orientation | Q7SCORE | | MMFLOOR |
| ADAS: Word Recognition | Q8SCORE | | MMCITY |
| ADAS: Recall Instructions | Q9SCORE | | MMAREA |
| ADAS: Spoken Language | Q10SCORE | | MMSTATE |
| ADAS: Word Finding | Q11SCORE | MMSE: Immediate Recall | MMBALL |
| ADAS: Comprehension | Q12SCORE | | MMFLAG |
| ADAS: Number Cancellation | Q13SCORE | | MMTREE |
| **NEUROBAT** | | MMSE: Attention | MMD |
| BNT: without cue and semantic cue | BNTTOTAL | | MML |
| CFT: animal | CATANIMSC | | MMO |
| Clock Drawing Test | CLOCKSCOR | | MMW |
| Clock Copying Test | COPYSCOR | MMSE: Delayed Recall | MMBALLDL |
| LM: Immediate Recall | LIMMTOTAL | | MMFLAGDL |
| LM: Delayed Recall | LDELTOTAL | | MMTREEDL |
| RAVLT: Trial 6 | AVTOT6 | MMSE: Language | MMWATCH |
| RAVLT: 30 min delay | AVDEL30MIN | | MMPENCIL |
| RAVLT: recognition | AVDELTOT | MMSE: Repetition | MMREPEAT |
| TMT: Part A | TRAASCOR | MMSE: Complex commands | MMHAND |
| TMT: Part B | TRABSCOR | | MMFOLD |
| | | | MMONFLR |
| | | | MMREAD |
| | | | MMWRITE |
| | | | MMDRAW |

**Table S2. Range of cognitive scores in ADNI-GO/2 baseline cohort.** Values represent mean (SD) min−max.

| | ADNI-GO/2 baseline | | |
|---|---|---|---|
| | AD | MCI | CN |
| N | 149 | 454 | 292 |
| ADAS: Word Recall | 6.3(1.4) 3−10 | 4.1(1.5) 0−9 | 2.9(1.3) 0−7 |
| ADAS: Commands | 0.3(0.7) 0−4 | 0.1(0.4) 0−5 | 0.1(0.4) 0−5 |
| ADAS: Construction | 0.9(0.8) 0−4 | 0.5(0.5) 0−2 | 0.4(0.5) 0−2 |
| ADAS: Delayed Word Recall | 8.6(1.5) 3−10 | 4.9(2.6) 0−10 | 2.9(1.9) 0−10 |
| ADAS: Naming | 0.6(0.8) 0−4 | 0.2(0.4) 0−2 | 0.1(0.2) 0−1 |
| ADAS: Ideational Praxis | 0.5(0.9) 0−5 | 0.1(0.3) 0−5 | 0.1(0.3) 0−2 |
| ADAS: Orientation | 2.4(1.8) 0−7 | 0.4(0.8) 0−8 | 0.1(0.4) 0−3 |
| ADAS: Word Recognition | 7.8(2.9) 0−12 | 3.4(2.6) 0−12 | 1.9(1.8) 0−12 |
| ADAS: Recall Instructions | 0.4(1) 0−5 | 0(0.2) 0−3 | 0(0.2) 0−2 |
| ADAS: Spoken Language | 0.6(0.8) 0−3 | 0.1(0.3) 0−3 | 0(0.1) 0−1 |
| ADAS: Word Finding | 0.7(0.9) 0−3 | 0.2(0.5) 0−2 | 0.1(0.4) 0−3 |
| ADAS: Comprehension | 0.3(0.7) 0−3 | 0.1(0.3) 0−2 | 0(0.2) 0−2 |
| ADAS: Number Cancellation | 1.7(1.3) 0−5 | 0.6(0.9) 0−5 | 0.4(0.6) 0−3 |
| MMSE: Orientation to time | 3.3(1.2) 0−5 | 4.8(0.5) 2−5 | 4.9(0.3) 4−5 |
| MMSE: Orientation to place | 3.9(1) 1−5 | 4.7(0.5) 2−5 | 4.8(0.4) 3−5 |
| MMSE: Immediate Recall | 2.9(0.2) 2−3 | 3(0.1) 2−3 | 3(0) 3−3 |
| MMSE: Attention | 4.1(1.3) 0−5 | 4.7(0.8) 1−5 | 4.8(0.6) 2−5 |
| MMSE: Delayed Recall | 0.6(0.9) 0−3 | 2.3(0.9) 0−3 | 2.7(0.5) 0−3 |
| MMSE: Language | 2(0.1) 1−2 | 2(0.1) 0−2 | 2(0.1) 1−2 |
| MMSE: Repetition | 0.8(0.4) 0−1 | 0.9(0.4) 0−1 | 0.9(0.3) 0−1 |
| MMSE: Complex commands | 5.4(0.8) 3−6 | 5.8(0.5) 4−6 | 5.9(0.4) 4−6 |
| BNT: without cue and semantic cue | 22(5.9) 3−30 | 27(3.6) 12−30 | 28.2(2.1) 17−30 |
| CFT: animal | 12.1(5.1) 2−26 | 18.1(5.1) 5−35 | 20.9(5.3) 8−38 |
| Clock Drawing Test | 3.4(1.4) 0−5 | 4.5(0.8) 1−5 | 4.7(0.6) 2−5 |
| Clock Copying Test | 4.4(1) 0−5 | 4.7(0.6) 0−5 | 4.9(0.4) 3−5 |
| LM: Immediate Recall | 4.1(2.7) 0−12 | 9.6(3.3) 0−22 | 14.3(3) 6−23 |
| LM: Delayed Recall | 1.5(1.9) 0−8 | 7.2(3.2) 0−15 | 13.3(3) 6−23 |
| RAVLT: Trial 6 | 1.8(1.8) 0−8 | 6.1(3.9) 0−15 | 9.1(3.5) 0−15 |
| RAVLT: 30 min delay | 0.7(1.4) 0−10 | 4.8(4.1) 0−15 | 7.6(4) 0−15 |
| RAVLT: recognition | 6.6(3.9) 0−15 | 11.3(3.1) 0−15 | 12.7(2.6) 1−15 |
| TMT: Part A | 61.7(33.4) 19−150 | 38.9(16.4) 13−150 | 33.7(11.3) 14−83 |
| TMT: Part B | 195.8(86.7) 55−300 | 106.2(58.1) 33−300 | 83.5(42.7) 32−300 |

**Table S2 (continued). Range of cognitive scores in ADNI-1 baseline cohort.** Values represent mean (SD) min−max.

| | ADNI-1 baseline | | |
|---|---|---|---|
| | AD | MCI | CN |
| N | 170 | 385 | 222 |
| ADAS: Word Recall | 6(1.4) 2.7−10 | 4.6(1.4) 1−8.3 | 2.8(1.1) 0−5.7 |
| ADAS: Commands | 0.4(0.7) 0−3 | 0.2(0.5) 0−4 | 0.1(0.4) 0−4 |
| ADAS: Construction | 0.8(0.6) 0−2 | 0.5(0.6) 0−3 | 0.4(0.5) 0−2 |
| ADAS: Delayed Word Recall | 8.6(1.5) 4−10 | 6.2(2.3) 0−10 | 2.8(1.7) 0−8 |
| ADAS: Naming | 0.5(0.7) 0−5 | 0.3(0.5) 0−3 | 0.1(0.3) 0−1 |
| ADAS: Ideational Praxis | 0.3(0.7) 0−5 | 0.1(0.4) 0−2 | 0(0.2) 0−1 |
| ADAS: Orientation | 2.1(1.7) 0−7 | 0.6(0.9) 0−5 | 0.1(0.3) 0−2 |
| ADAS: Word Recognition | 6.6(2.8) 1−12 | 4.6(2.7) 0−12 | 2.5(2.3) 0−12 |
| ADAS: Recall Instructions | 0.3(0.8) 0−5 | 0.1(0.4) 0−4 | 0(0) 0−0 |
| ADAS: Spoken Language | 0.3(0.6) 0−2 | 0.1(0.3) 0−3 | 0(0.1) 0−1 |
| ADAS: Word Finding | 0.5(0.8) 0−4 | 0.3(0.6) 0−4 | 0.1(0.3) 0−2 |
| ADAS: Comprehension | 0.2(0.5) 0−2 | 0.1(0.3) 0−3 | 0(0.1) 0−1 |
| ADAS: Number Cancellation | 1.7(1.2) 0−5 | 1(0.9) 0−5 | 0.5(0.7) 0−4 |
| MMSE: Orientation to time | 3.4(1.3) 1−5 | 4.5(0.7) 1−5 | 4.9(0.3) 3−5 |
| MMSE: Orientation to place | 3.8(1) 1−5 | 4.5(0.7) 2−5 | 4.9(0.3) 3−5 |
| MMSE: Immediate Recall | 3(0.2) 1−3 | 3(0.2) 1−3 | 3(0.1) 2−3 |
| MMSE: Attention | 4.3(1.1) 1−5 | 4.7(0.8) 1−5 | 4.9(0.5) 1−5 |
| MMSE: Delayed Recall | 0.8(0.9) 0−3 | 1.8(1.1) 0−3 | 2.7(0.6) 1−3 |
| MMSE: Language | 2(0.1) 1−2 | 2(0.1) 1−2 | 2(0) 2−2 |
| MMSE: Repetition | 0.8(0.4) 0−1 | 0.8(0.4) 0−1 | 0.9(0.3) 0−1 |
| MMSE: Complex commands | 5.4(0.7) 3−6 | 5.7(0.5) 4−6 | 5.8(0.4) 4−6 |
| BNT: without cue and semantic cue | 22.7(5.9) 2−30 | 25.5(4.1) 4−30 | 27.9(2.3) 18−30 |
| CFT: animal | 12.7(4.9) 2−27 | 15.9(4.9) 5−30 | 20(5.6) 6−38 |
| Clock Drawing Test | 3.5(1.2) 0−5 | 4.2(1) 0−5 | 4.7(0.7) 1−5 |
| Clock Copying Test | 4.4(0.8) 1−5 | 4.6(0.7) 0−5 | 4.9(0.4) 2−5 |
| LM: Immediate Recall | 4.2(2.9) 0−14 | 7.1(3.1) 0−18 | 13.8(3.5) 3−22 |
| LM: Delayed Recall | 1.3(1.9) 0−8 | 3.8(2.7) 0−12 | 13(3.6) 5−22 |
| RAVLT: Trial 6 | 1.7(1.8) 0−9 | 3.8(3.2) 0−15 | 8.1(3.4) 0−15 |
| RAVLT: 30 min delay | 0.8(1.7) 0−9 | 2.8(3.3) 0−15 | 7.4(3.7) 0−15 |
| RAVLT: recognition | 7.3(4) 0−15 | 9.7(3.6) 0−15 | 12.9(2.5) 2−15 |
| TMT: Part A | 64.2(33.5) 18−150 | 44.4(22.1) 17−150 | 36.5(13.1) 17−102 |
| TMT: Part B | 197.9(86.7) 35−300 | 129.8(72.8) 0−300 | 89.4(44.6) 34−300 |

**Table S2 (continued). Range of cognitive scores in ADNI-GO/2 month 12 cohort.** Values represent mean (SD) min−max.

| | ADNI-GO/2 month 12 | | |
|---|---|---|---|
| | AD | MCI | CN |
| N | 114 | 353 | 201 |
| ADAS: Word Recall | 6.2(1.6) 1−9 | 3.7(1.6) 0−9 | 2.5(1.3) 0−5 |
| ADAS: Commands | 0.5(0.7) 0−3 | 0.1(0.3) 0−2 | 0.1(0.5) 0−5 |
| ADAS: Construction | 0.8(0.7) 0−3 | 0.5(0.6) 0−3 | 0.4(0.5) 0−2 |
| ADAS: Delayed Word Recall | 8.6(1.8) 3−10 | 4.5(2.7) 0−10 | 2.4(1.7) 0−9 |
| ADAS: Naming | 0.6(0.7) 0−3 | 0.1(0.3) 0−2 | 0(0.2) 0−1 |
| ADAS: Ideational Praxis | 0.4(0.7) 0−5 | 0.1(0.3) 0−3 | 0(0.4) 0−5 |
| ADAS: Orientation | 2.6(1.9) 0−7 | 0.4(0.7) 0−4 | 0.1(0.4) 0−2 |
| ADAS: Word Recognition | 7.5(3) 0−12 | 3.2(2.7) 0−12 | 1.9(1.9) 0−11 |
| ADAS: Recall Instructions | 0.5(1) 0−3 | 0(0.1) 0−1 | 0(0.1) 0−1 |
| ADAS: Spoken Language | 0.5(0.8) 0−3 | 0.1(0.3) 0−2 | 0(0.1) 0−1 |
| ADAS: Word Finding | 0.9(1.1) 0−4 | 0.3(0.5) 0−3 | 0.1(0.4) 0−2 |
| ADAS: Comprehension | 0.4(0.8) 0−3 | 0.1(0.3) 0−2 | 0(0.1) 0−1 |
| ADAS: Number Cancellation | 1.7(1.4) 0−5 | 0.5(0.8) 0−4 | 0.3(0.6) 0−4 |
| MMSE: Orientation to time | 3.1(1.6) 0−5 | 4.6(0.7) 1−5 | 4.9(0.3) 3−5 |
| MMSE: Orientation to place | 3.7(1.2) 0−5 | 4.8(0.5) 3−5 | 4.9(0.3) 3−5 |
| MMSE: Immediate Recall | 2.9(0.3) 1−3 | 3(0.1) 2−3 | 3(0.1) 2−3 |
| MMSE: Attention | 4.2(1.3) 0−5 | 4.8(0.7) 1−5 | 4.9(0.6) 0−5 |
| MMSE: Delayed Recall | 0.6(0.9) 0−3 | 1.9(1.2) 0−3 | 2.5(0.8) 0−3 |
| MMSE: Language | 2(0.1) 1−2 | 2(0.1) 1−2 | 2(0) 2−2 |
| MMSE: Repetition | 0.7(0.5) 0−1 | 0.9(0.3) 0−1 | 0.9(0.3) 0−1 |
| MMSE: Complex commands | 5.5(0.8) 1−6 | 5.8(0.5) 4−6 | 5.8(0.4) 4−6 |
| BNT: without cue and semantic cue | 22.3(6.2) 5−30 | 27.6(3) 12−30 | 28.7(1.8) 20−30 |
| CFT: animal | 11.9(4.7) 2−25 | 18(5.2) 7−34 | 21.7(5.2) 11−34 |
| Clock Drawing Test | 3.6(1.3) 1−5 | 4.6(0.7) 1−5 | 4.7(0.6) 2−5 |
| Clock Copying Test | 4.4(1.2) 0−5 | 4.8(0.4) 2−5 | 4.9(0.4) 3−5 |
| LM: Immediate Recall | 4.6(3.3) 0−16 | 10.8(4.3) 1−21 | 15.2(3.1) 7−24 |
| LM: Delayed Recall | 1.6(2.8) 0−12 | 8.5(4.9) 0−22 | 13.8(3.6) 3−23 |
| RAVLT: Trial 6 | 1.7(1.9) 0−8 | 6.2(4.1) 0−15 | 9.3(3.6) 0−15 |
| RAVLT: 30 min delay | 0.5(1.4) 0−9 | 4.7(4.4) 0−15 | 8(4.3) 0−15 |
| RAVLT: recognition | 6.4(3.8) 0−15 | 11.3(3.3) 0−15 | 13.1(2.3) 2−15 |
| TMT: Part A | 60.2(30.8) 23−151 | 38(15.8) 13−117 | 32(11.6) 15−101 |
| TMT: Part B | 204.3(90.3) 41−306 | 102.9(57.5) 31−300 | 79.8(43.1) 31−300 |

**Table S2 (continued). Range of cognitive scores in ADNI-2/3 Tau PET cohort.** Values represent mean (SD) min−max.

| | ADNI-2/3 Tau PET | | |
|---|---|---|---|
| | AD | MCI | CN |
| N | – | 76 | – |
| ADAS: Word Recall | | 3.9(1.3) 1−6.7 | |
| ADAS: Commands | | 0.9(0.8) 0−5 | |
| ADAS: Construction | | 0.5(0.7) 0−3 | |
| ADAS: Delayed Word Recall | | 5(2.4) 0−10 | |
| ADAS: Naming | | 0.2(0.4) 0−2 | |
| ADAS: Ideational Praxis | | 0(0.1) 0−1 | |
| ADAS: Orientation | | 0.5(0.9) 0−5 | |
| ADAS: Word Recognition | | 5.4(2.8) 0−12 | |
| ADAS: Recall Instructions | | 0(0.2) 0−1 | |
| ADAS: Spoken Language | | 0.1(0.2) 0−1 | |
| ADAS: Word Finding | | 0.1(0.3) 0−1 | |
| ADAS: Comprehension | | 0(0) 0−0 | |
| ADAS: Number Cancellation | | 1.3(1.1) 0−4 | |
| MMSE: Orientation to time | | 4.6(0.7) 2−5 | |
| MMSE: Orientation to place | | 4.9(0.4) 4−5 | |
| MMSE: Immediate Recall | | 3(0) 3−3 | |
| MMSE: Attention | | 4.7(0.9) 1−5 | |
| MMSE: Delayed Recall | | 2.1(1.3) 0−3 | |
| MMSE: Language | | 2(0) 2−2 | |
| MMSE: Repetition | | 0.8(0.4) 0−1 | |
| MMSE: Complex commands | | 5.7(0.5) 4−6 | |
| BNT: without cue and semantic cue | | 27.9(2.4) 23−30 | |
| CFT: animal | | 17.7(5.9) 6−32 | |
| Clock Drawing Test | | 4.5(0.9) 2−5 | |
| Clock Copying Test | | 4.8(0.5) 3−5 | |
| LM: Immediate Recall | | 11.4(4.7) 0−19 | |
| LM: Delayed Recall | | 9(4.9) 0−20 | |
| RAVLT: Trial 6 | | 6.1(4) 0−15 | |
| RAVLT: 30 min delay | | 4.4(4.1) 0−15 | |
| RAVLT: recognition | | 11.3(3.7) 0−15 | |
| TMT: Part A | | 37.8(14.8) 18−106 | |
| TMT: Part B | | 108(70.2) 34−300 | |

**Table S3. 28 cognitive scores after combination and corresponding abbreviations.** The left column lists combined scores from Table S1 and right column corresponds to abbreviations of these scores shown in Fig. 4.

| **Behavioral Description** | **Behavioral Abbreviation** |
|---|---|
| ADAS: Word Recall | ADAS: Word Recall |
| ADAS: Commands | ADAS: Commands |
| ADAS: Construction | ADAS: Construction |
| ADAS: Delayed Word Recall | ADAS: Del. Recall |
| ADAS: Naming + MMSE: Language | ADAS: Nam. + MMSE: Lan. |
| ADAS: Ideational Praxis | ADAS: Ide. Praxis |
| ADAS: Orientation | ADAS: Orientation |
| ADAS: Word Recognition | ADAS: Word Recog. |
| ADAS: Spoken Language + Comprehension | ADAS: Spo. Com. |
| ADAS: Word Finding | ADAS: Word Find. |
| ADAS: Number Cancellation | ADAS: Num. Can. |
| MMSE: Orientation to time | MMSE: Ori. Time |
| MMSE: Orientation to place | MMSE: Ori. Place |
| MMSE: Immediate Recall + Delayed Recall | MMSE: Imm. Del. |
| MMSE: Attention | MMSE: Attention |
| MMSE: Repetition | MMSE: Repetition |
| MMSE: Complex Commands | MMSE: Com. Com. |
| BNT: without cue and semantic cue | BNT |
| CFT: animal | CFT: Animal |
| Clock Drawing Test | Clock Drawing Test |
| Clock Copying Test | Clock Copying Test |
| LM: Immediate Recall | LM: Imm. Recall |
| LM: Delayed Recall | LM: Del. Recall |
| RAVLT: Trail 6 | RAVLT: Trail 6 |
| RAVLT: 30 min delay | RAVLT: 30 min delay |
| RAVLT: Recognition | RAVLT: Recognition |
| TMT: Part A | TMT: Part A |
| TMT: Part B | TMT: Part B |

**Table S4. Characteristics of 149 AD dementia patients by factor.** Data are weighted averages (weighted standard deviation) with weights corresponding to factor probabilities. Highlighted p values (blue) are characteristics significantly different across factors. *Computed by linear hypothesis test on GLM or likelihood ratio test on logistic regression for sex (see Methods). ‡Only available for 146 patients. §The original counts were 0, 1 or 2. Only available for 145 patients.

| | MTL-Memory | Lateral Temporal-Language | Posterior Cortical-Executive | Overall p* |
|---|---|---|---|---|
| Baseline age (years) | 75.4 (7.7) | 74.9 (7.6) | 73.4 (8.8) | **2e-4** |
| Education (years) | 15.9 (2.7) | 15.8 (2.7) | 15.6 (2.6) | 0.44 |
| Sex (0 for male) | 0.4 (0.5) | 0.4 (0.5) | 0.4 (0.5) | 0.91 |
| Amyloid (SUVR)‡ | 1.4 (0.2) | 1.4 (0.2) | 1.4 (0.2) | 0.16 |
| APOE ε2§ | 0.06 (0.3) | 0.07 (0.3) | 0.06 (0.3) | 0.97 |
| APOE ε4§ | 0.87 (0.7) | 0.86 (0.7) | 0.89 (0.7) | 0.82 |

**Table S5. Standard deviation of cognitive scores in ADNI-1 and ADNI-GO/2 CN participants.**

| ADNI-1 CN (N = 222) | | ADNI-GO/2 CN (N = 292) | |
|---|---|---|---|
| Cognitive Scores | Standard Deviation | Cognitive Scores | Standard Deviation |
| ADAS: Recall Instructions | 0 | MMSE: Immediate Recall | 0 |
| MMSE: Language | 0 | MMSE: Language | 0.058521 |
| ADAS: Comprehension | 0.067116 | ADAS: Spoken Language | 0.14211 |
| MMSE: Immediate Recall | 0.094701 | ADAS: Recall Instructions | 0.15415 |
| ADAS: Spoken Language | 0.13332 | ADAS: Comprehension | 0.21664 |
| ADAS: Ideational Praxis | 0.2175 | ADAS: Naming | 0.24707 |
| ADAS: Naming | 0.25157 | ADAS: Ideational Praxis | 0.27887 |
| MMSE: Repetition | 0.26652 | MMSE: Orientation to time | 0.31291 |
| ADAS: Word Finding | 0.30836 | MMSE: Repetition | 0.31291 |
| MMSE: Orientation to time | 0.30836 | ADAS: Word Finding | 0.35018 |
| ADAS: Orientation | 0.33905 | ADAS: Orientation | 0.35193 |
| MMSE: Orientation to place | 0.34607 | Clock Copying Test | 0.36978 |
| ADAS: Commands | 0.36457 | MMSE: Orientation to place | 0.3802 |
| MMSE: Complex commands | 0.38148 | MMSE: Complex commands | 0.38225 |
| Clock Copying Test | 0.42899 | ADAS: Commands | 0.38745 |
| ADAS: Construction | 0.50089 | ADAS: Construction | 0.52695 |
| MMSE: Attention | 0.51436 | MMSE: Delayed Recall | 0.52812 |
| MMSE: Delayed Recall | 0.55441 | Clock Drawing Test | 0.601 |
| Clock Drawing Test | 0.66811 | MMSE: Attention | 0.64344 |
| ADAS: Number Cancellation | 0.68254 | ADAS: Number Cancellation | 0.64931 |
| ADAS: Word Recall | 1.1443 | ADAS: Word Recall | 1.3187 |
| ADAS: Delayed Word Recall | 1.7116 | ADAS: Word Recognition | 1.7797 |
| ADAS: Word Recognition | 2.2864 | ADAS: Delayed Word Recall | 1.8982 |
| BNT: without cue and semantic cue | 2.3108 | BNT: without cue and semantic cue | 2.0933 |
| RAVLT: recognition | 2.526 | RAVLT: recognition | 2.6186 |
| RAVLT: Trial 6 | 3.4189 | LM: Immediate Recall | 3.0363 |
| LM: Immediate Recall | 3.4926 | LM: Delayed Recall | 3.0429 |
| LM: Delayed Recall | 3.5898 | RAVLT: Trial 6 | 3.4539 |
| RAVLT: 30 min delay | 3.6879 | RAVLT: 30 min delay | 4.0232 |
| CFT: animal | 5.5864 | CFT: animal | 5.2792 |
| TMT: Part A | 13.0895 | TMT: Part A | 11.283 |
| TMT: Part B | 44.611 | TMT: Part B | 42.7338 |

**Table S6. Standard deviation of combined cognitive scores in ADNI-1 and ADNI-GO/2 CN participants.**

| ADNI-1 CN (N = 222) | | ADNI-GO/2 CN (N = 292) | |
|---|---|---|---|
| Cognitive Scores | Standard Deviation | Cognitive Scores | Standard Deviation |
| ADAS:Spoken Language+Comprehension | 0.14871 | ADAS:Naming+MMSE:Language | 0.25302 |
| ADAS:Naming+MMSE:Language | 0.25157 | ADAS:Spoken Language+Comprehension | 0.28189 |
| MMSE:Immediate Recall+Delayed Recall | 0.5654 | MMSE:Immediate Recall+Delayed Recall | 0.52812 |

**Table S7. Summary of terminologies and their descriptions.**

| Terminology | Description | Bayesian Model Symbol |
|---|---|---|
| Atrophy Pattern | Probability that a latent factor is associated with atrophy at a voxel [Pr(Voxel \| Factor)] | $\beta^1$ |
| Cognitive Profile | Probability that a latent factor is associated with a cognitive score deficit [Pr(Score \| Factor)] | $\beta^2$ |
| Atrophy-Cognitive Loading | Probability that a participant expresses a latent factor based on the patient's brain atrophy and cognitive scores [Pr(Factor \| Participant)] | $\gamma$ |
| Atrophy Loading | Probability that a participant expresses a latent factor based only the participant's brain atrophy [Pr(Factor \| Participant's Atrophy)] | $\gamma^1$ |
| Cognitive Loading | Probability that a participant expresses a latent factor based only on the participant's cognitive scores [Pr(Factor \| Participant's Cognition)] | $\gamma^2$ |

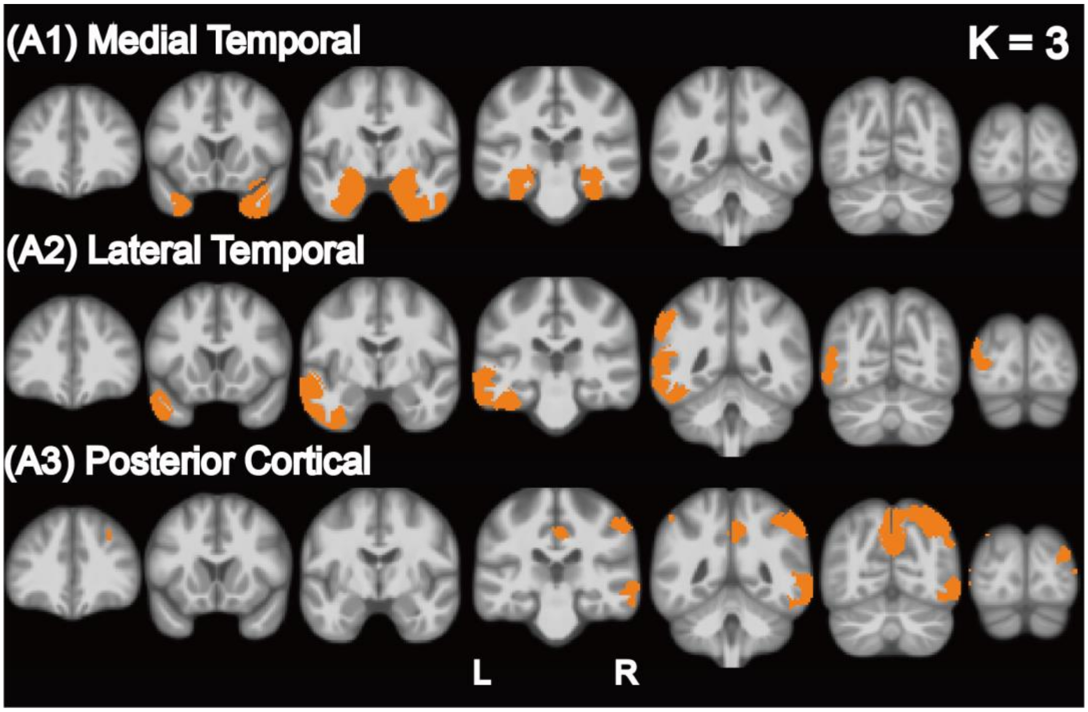


**Figure S1. Masks containing top 5% voxels for each latent factor in ADNI-GO/2.** The SUVR signal was averaged across voxels within each mask to obtain the regional Tau deposition for each factor.

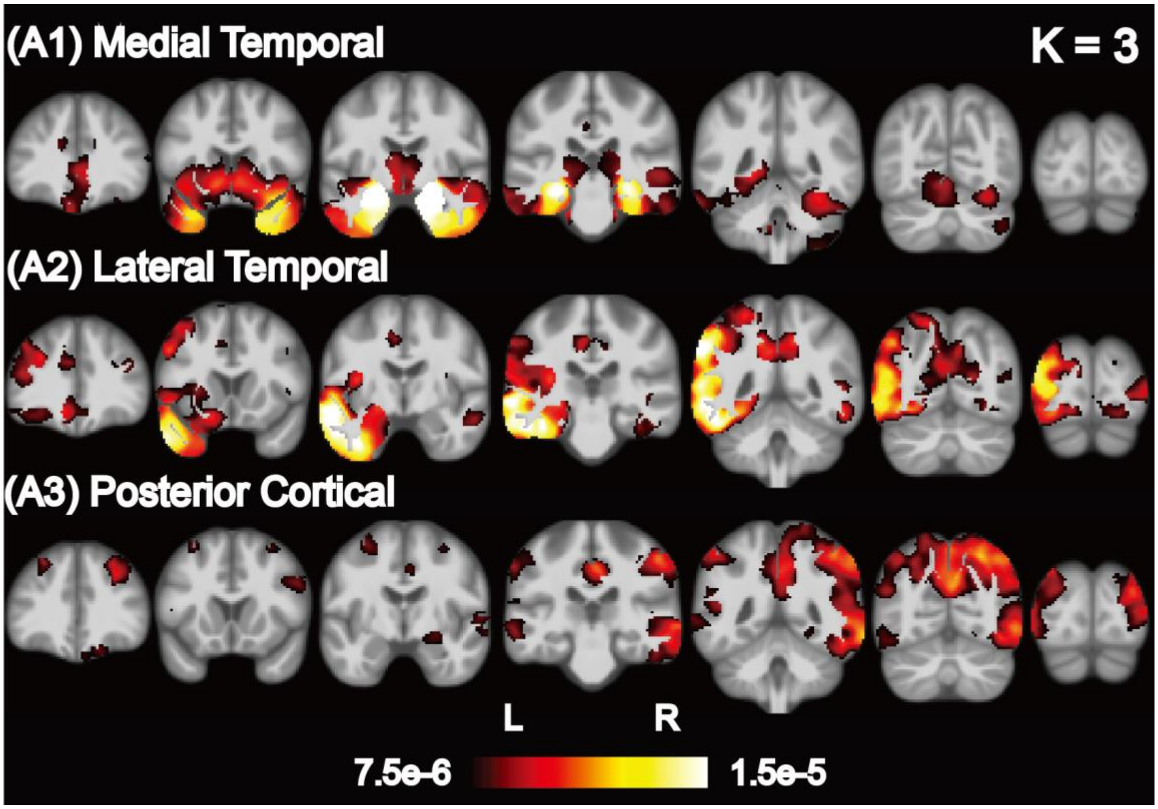


**Figure S2. Probabilistic atrophy maps of K = 3 latent factors in ADNI-GO/2.** Brighter color indicates higher probability of atrophy at that voxel for a particular factor (i.e., Pr(Voxel | Factor)). Each of the three factors was associated with a distinct pattern of brain atrophy.

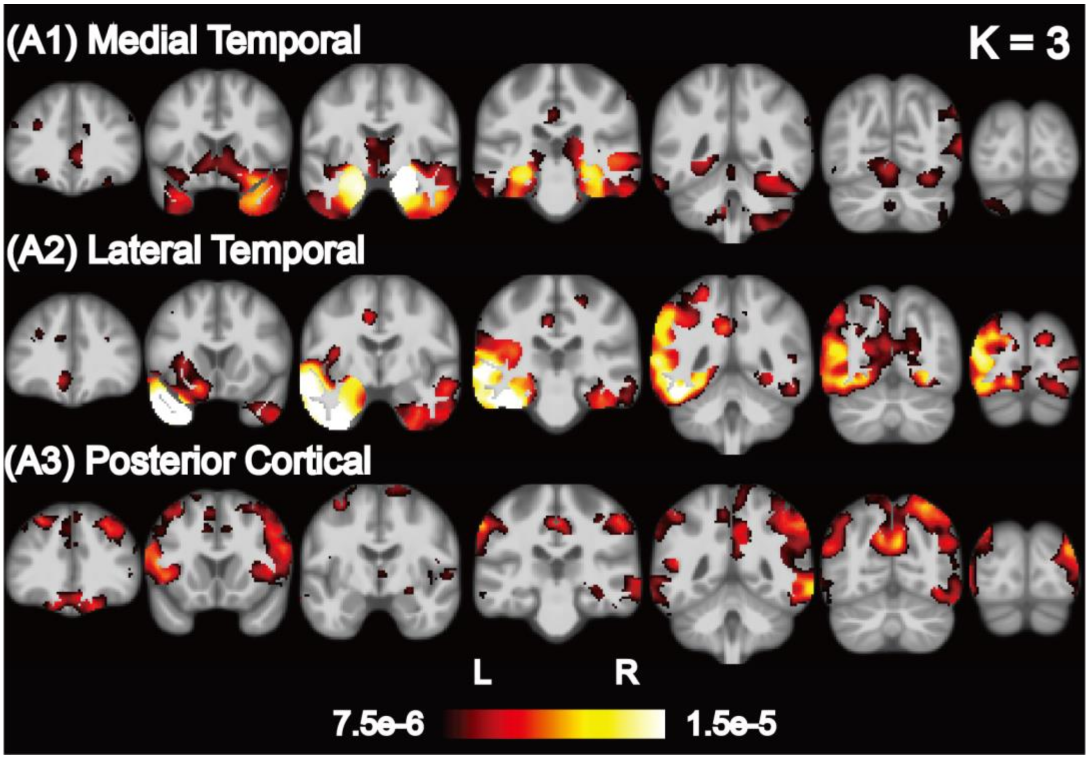


**Figure S3. Probabilistic atrophy maps for K=3 latent factors in ADNI-GO/2 <u>A+ AD dementia participants with age > 70</u>.** Bright color indicates higher probability of atrophy at that spatial location for a particular atrophy factor (i.e., Pr(Voxel | Factor)). Each of the three factors was associated with a distinct pattern of brain atrophy.

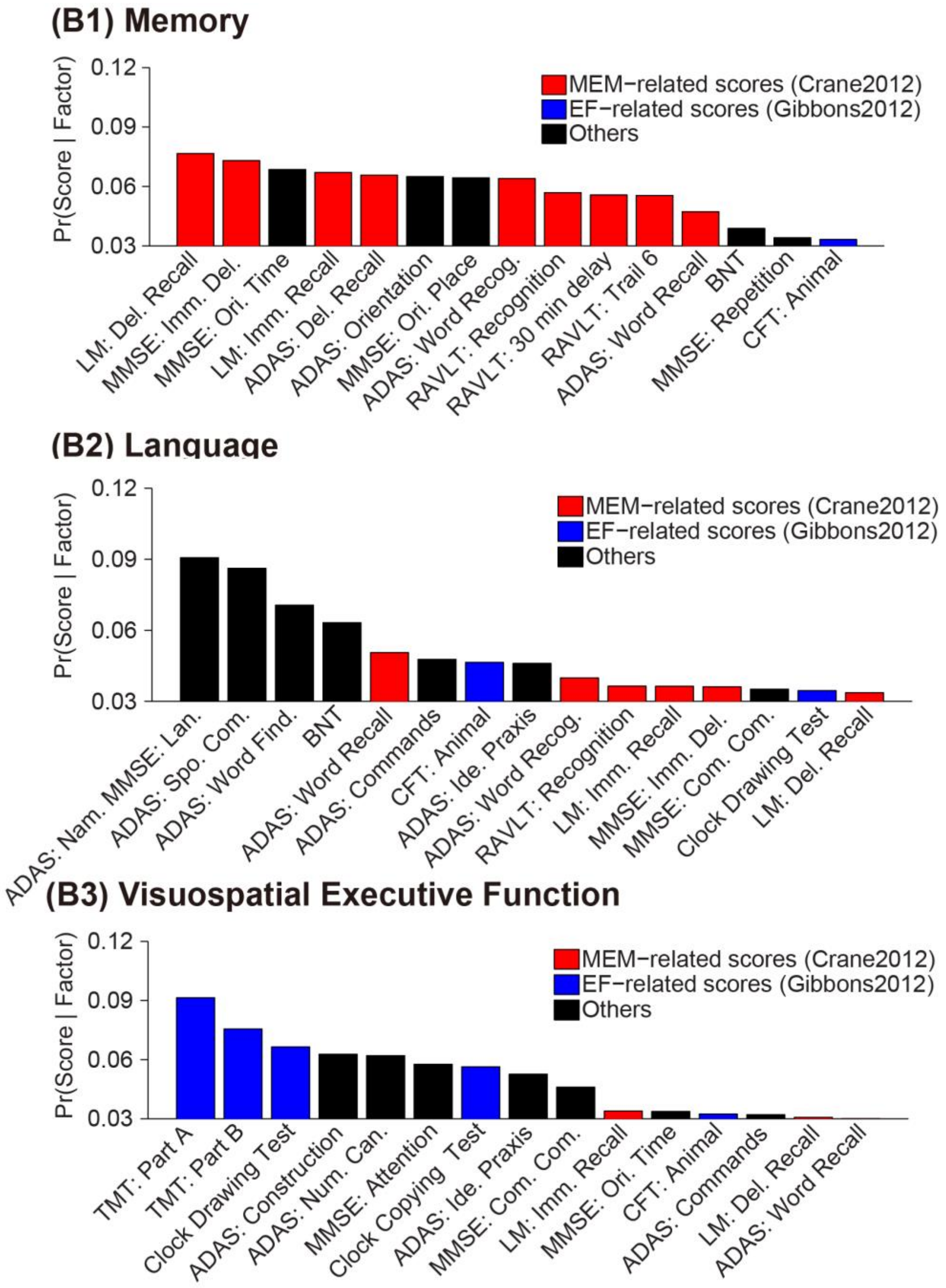


**Figure S4. Probabilistic cognitive deficits for K=3 latent factors in ADNI-GO/2 A+ AD dementia participants with age > 70.** Red bars indicate scores associated with memory, blue bars indicate scores associated with executive function and black bars indicate scores that do not belong to either memory or executive function. The probabilities of cognitive scores associated with a factor (i.e., Pr(Score | Factor)) are sorted from high to low and only top 15/28 cognitive scores are shown in the plot.

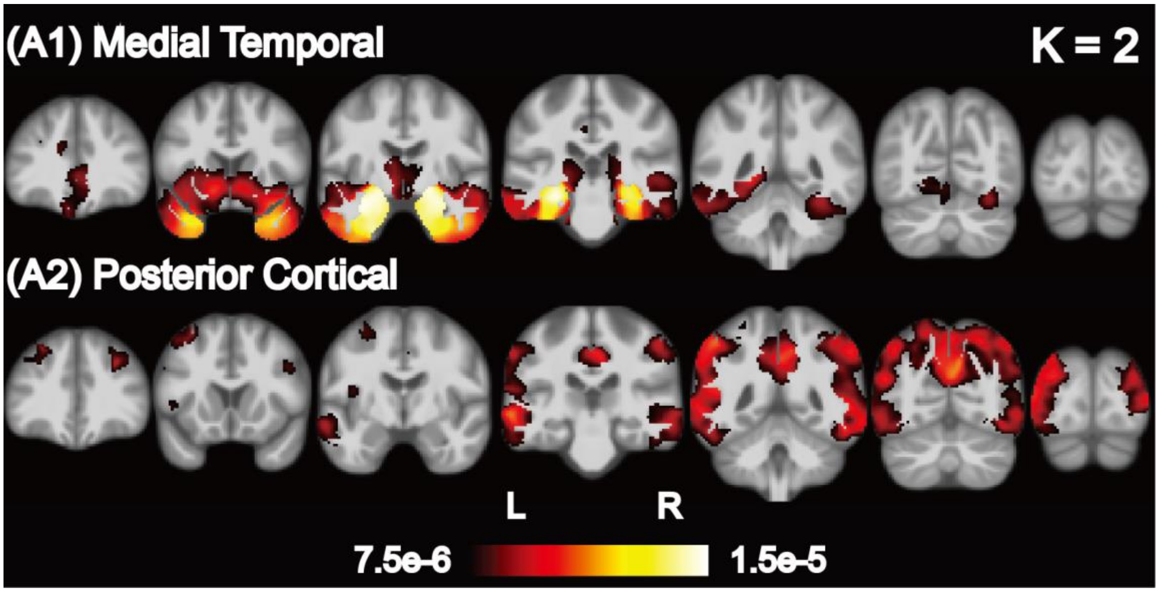


**Figure S5. Probabilistic atrophy maps for K = 2 latent factors in ADNI-GO/2.** Brighter color indicates higher probability of atrophy at that voxel for a particular factor (i.e., Pr(Voxel | Factor)).

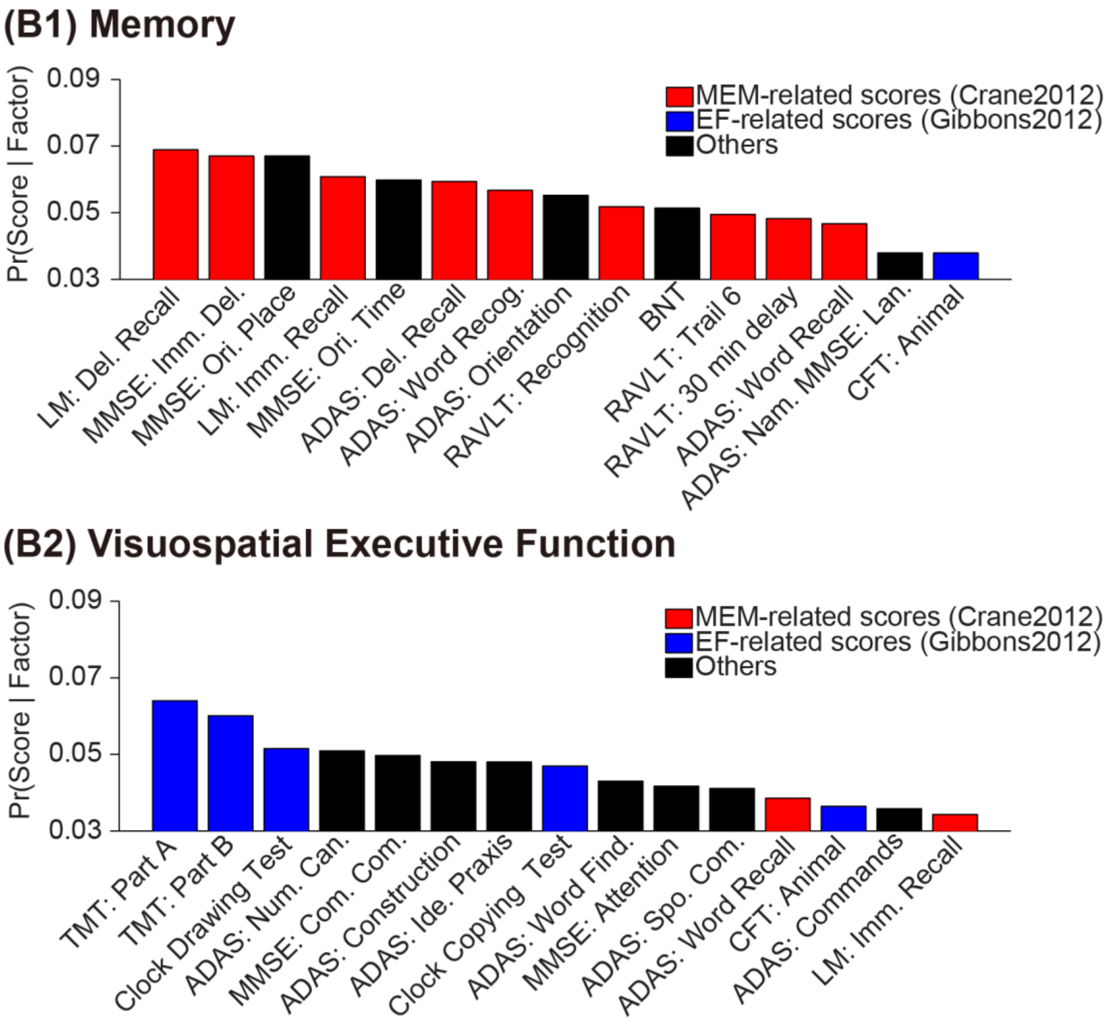


**Figure S6. Probabilistic cognitive deficits for K = 2 latent factors in ADNI-GO/2.** Red bars indicate scores associated with memory, blue bars indicate scores associated with executive function and black bars indicate scores that do not belong to either memory or executive function. The probabilities of cognitive scores associated with a factor (i.e., Pr(Score | Factor)) are sorted from high to low and only top 15/28 cognitive scores are shown in the plot.

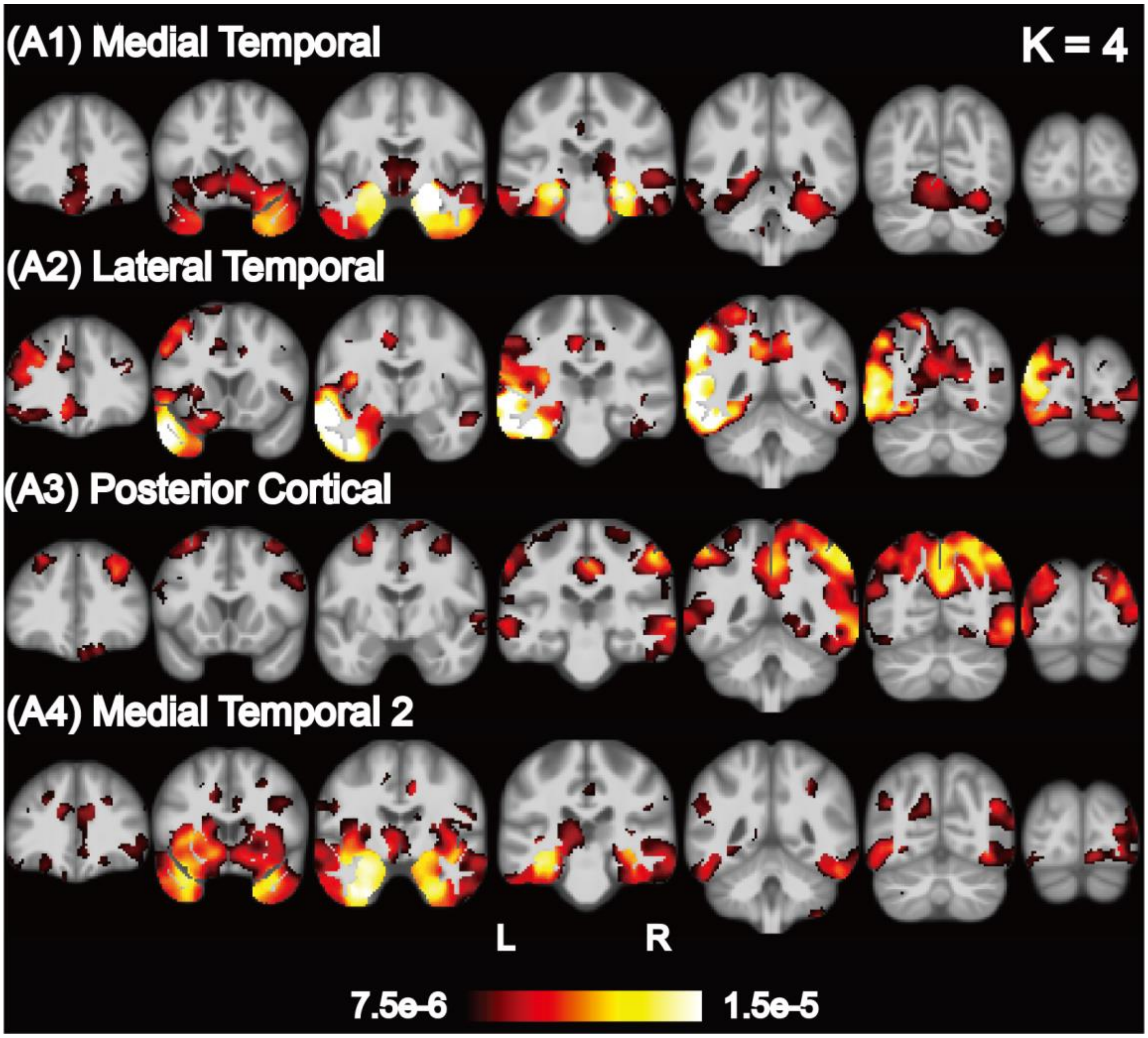


**Figure S7. Probabilistic atrophy maps for K = 4 latent factors in ADNI-GO/2.** Brighter color indicates higher probability of atrophy at that voxel for a particular factor (i.e., Pr(Voxel | Factor)).

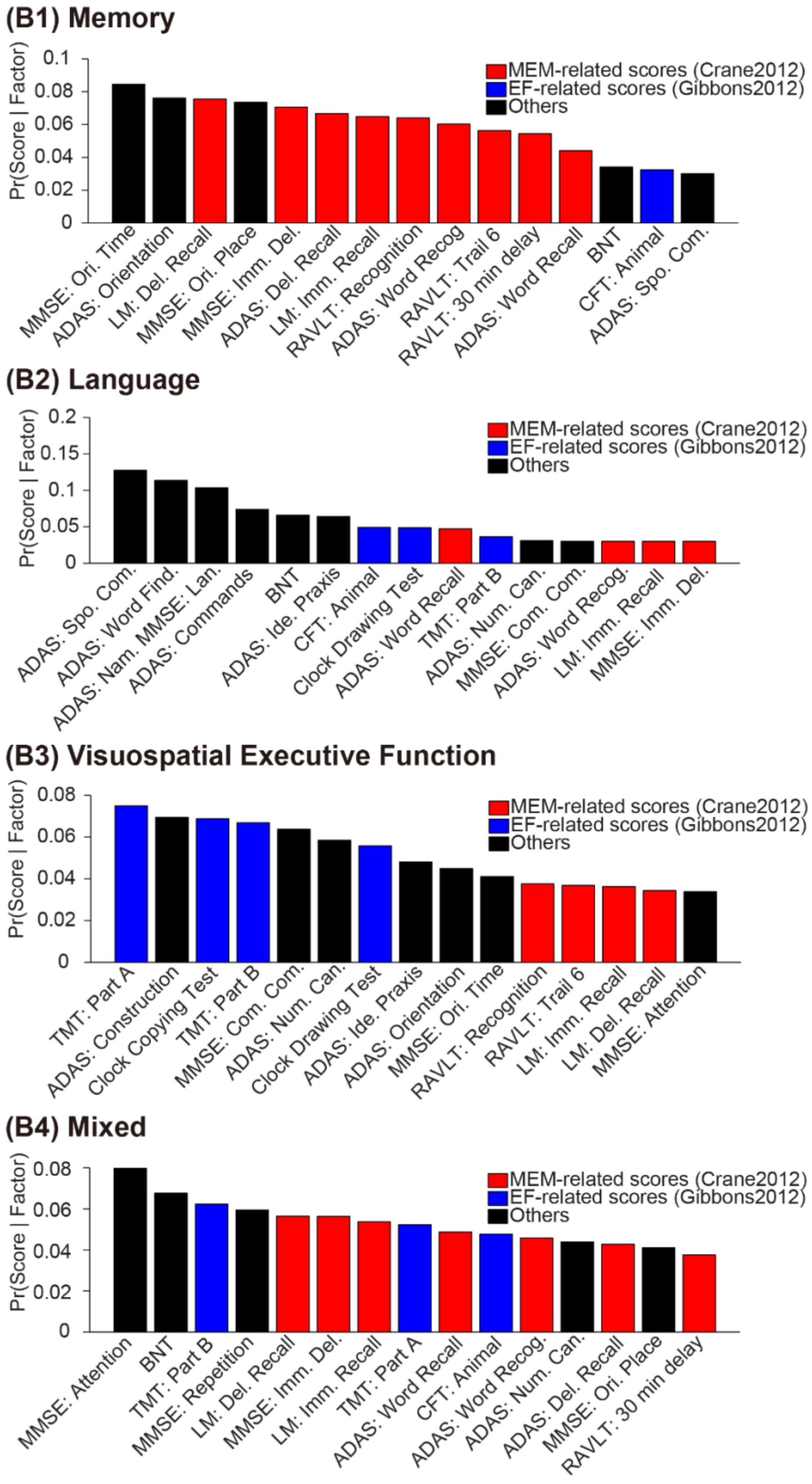


**Figure S8. Probabilistic cognitive deficits for K = 4 latent factors in ADNI-GO/2.**

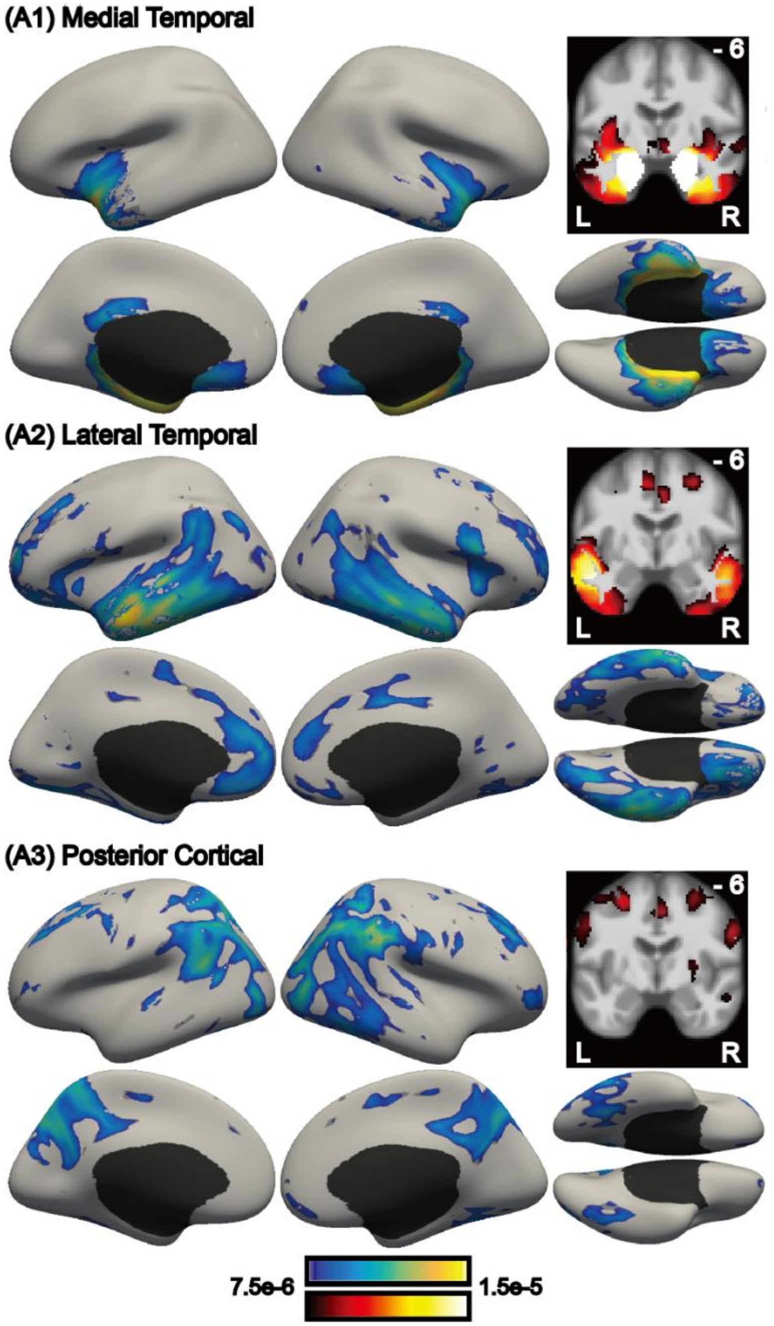


**Figure S9. Probabilistic atrophy maps for K = 3 latent factors in ADNI-1.** Brighter color indicates higher probability of atrophy at that voxel for a particular factor (i.e., Pr(Voxel | Factor)). Each of the three factors was associated with a distinct pattern of brain atrophy.

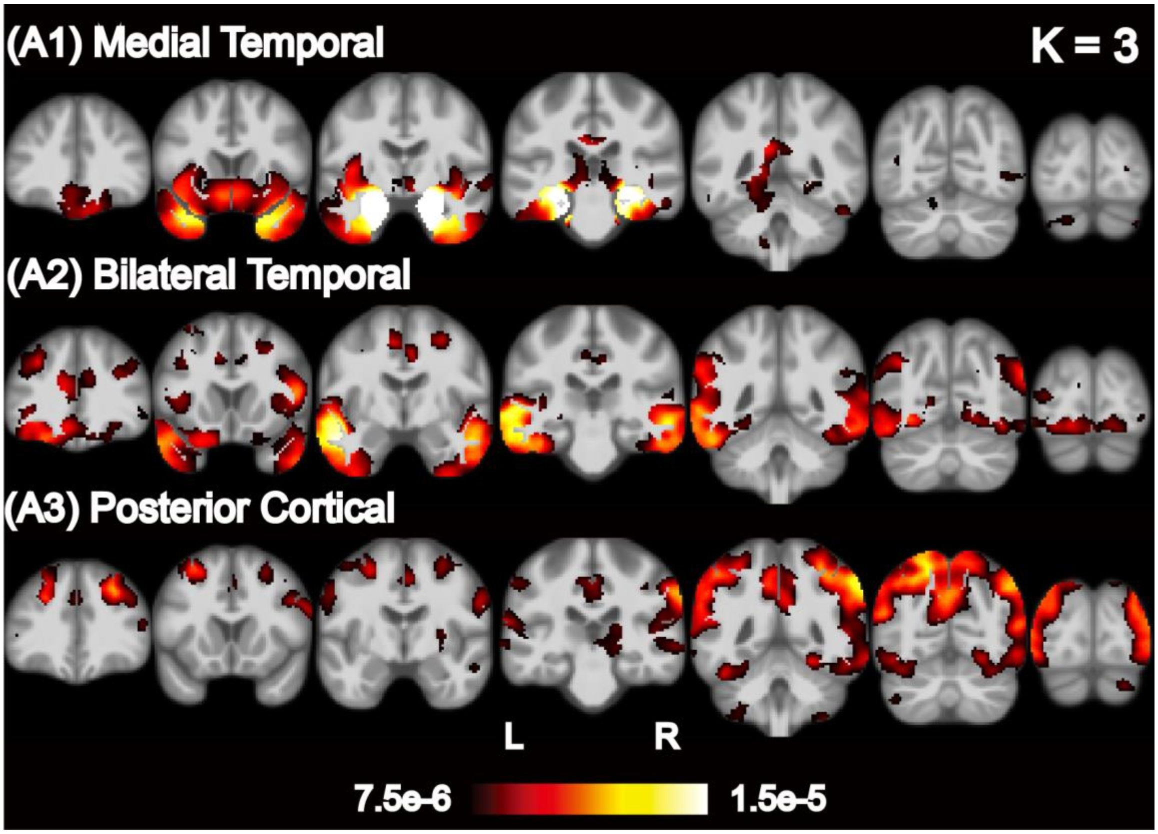


**Figure S10. Probabilistic atrophy maps for K=3 latent factors in ADNI-1.** Bright color indicates higher probability of atrophy at that spatial location for a particular atrophy factor (i.e., Pr(Voxel | Factor)). Each of the three factors was associated with a distinct pattern of brain atrophy.

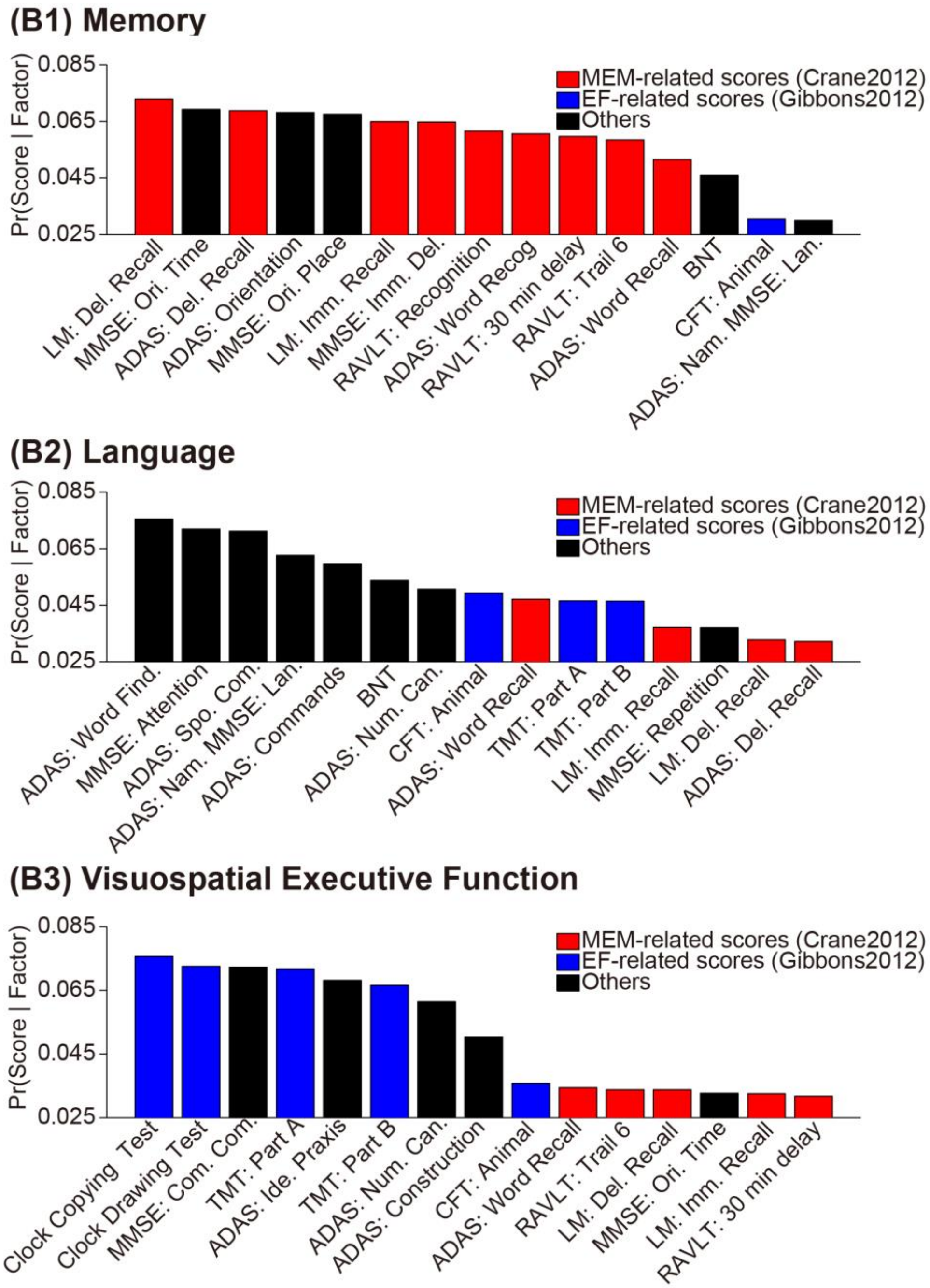


**Figure S11. Probabilistic cognitive deficits for K=3 latent factors in ADNI-1.** Red bars indicate scores associated with memory, blue bars indicate scores associated with executive function and black bars indicate scores that do not belong to either memory or executive function. The probabilities of cognitive scores associated with a factor (i.e., Pr(Score | Factor)) are sorted from high to low and only top 15/28 cognitive scores are shown in the plot.

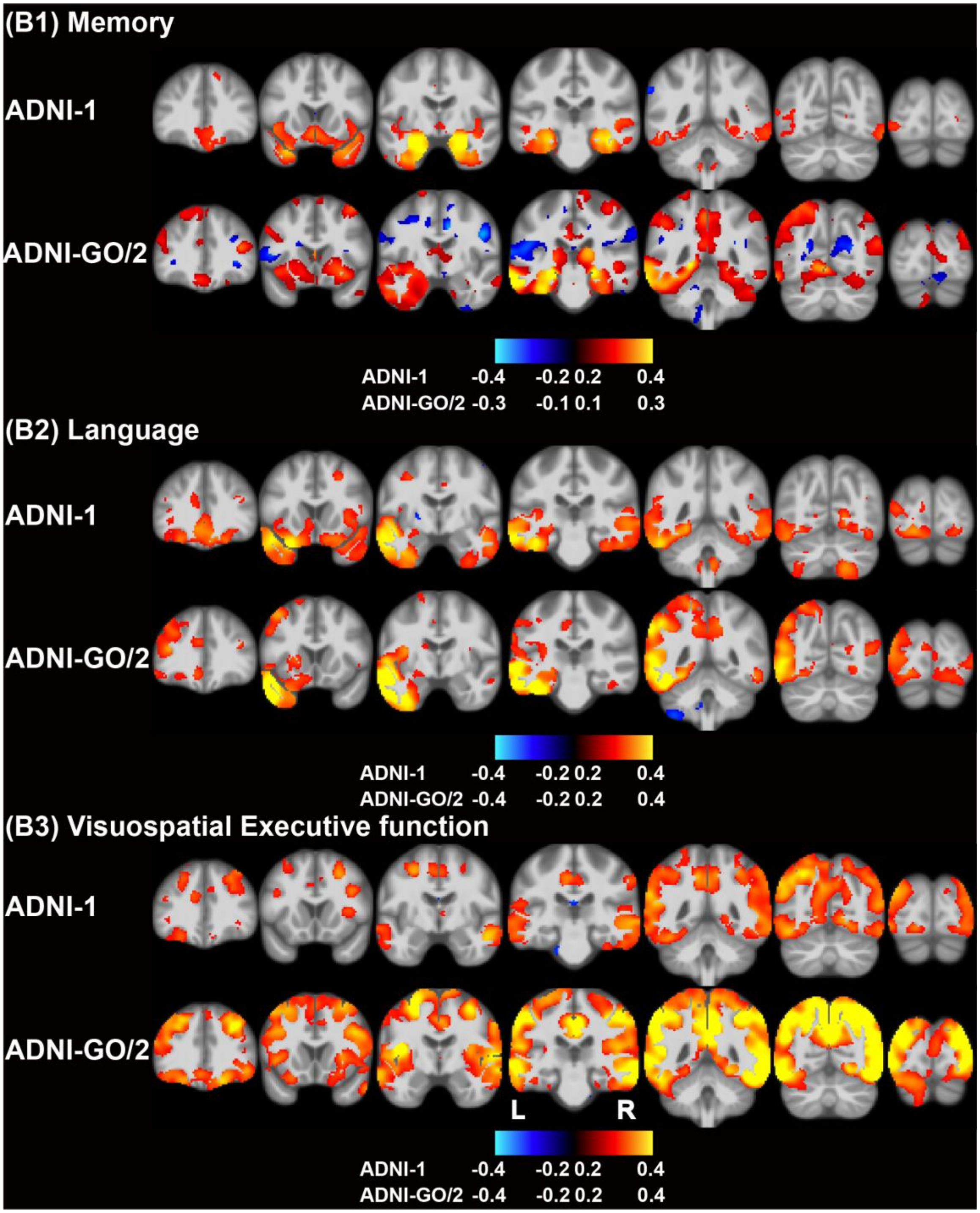


**Figure S12. Correlation maps between atrophy z score and cognitive mean z score of top 5 scores (Fig. 3) for each factor in ADNI-1 or ADNI-GO/2 AD.** Hot color indicates positive correlation and cold color indicates negative correlation.

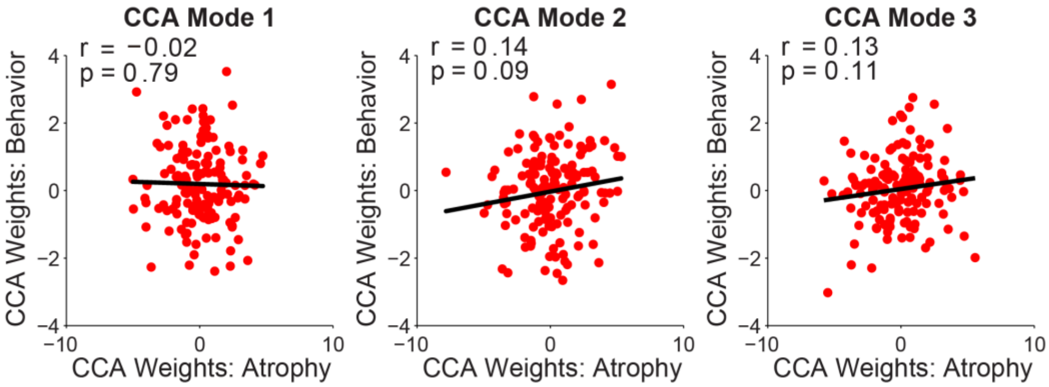


**Figure S13. 10-fold cross validation of three CCA modes.** We performed 10-fold cross-validation among the AD dementia patients in ADNI-GO/2. Each red dot represents an AD patient.

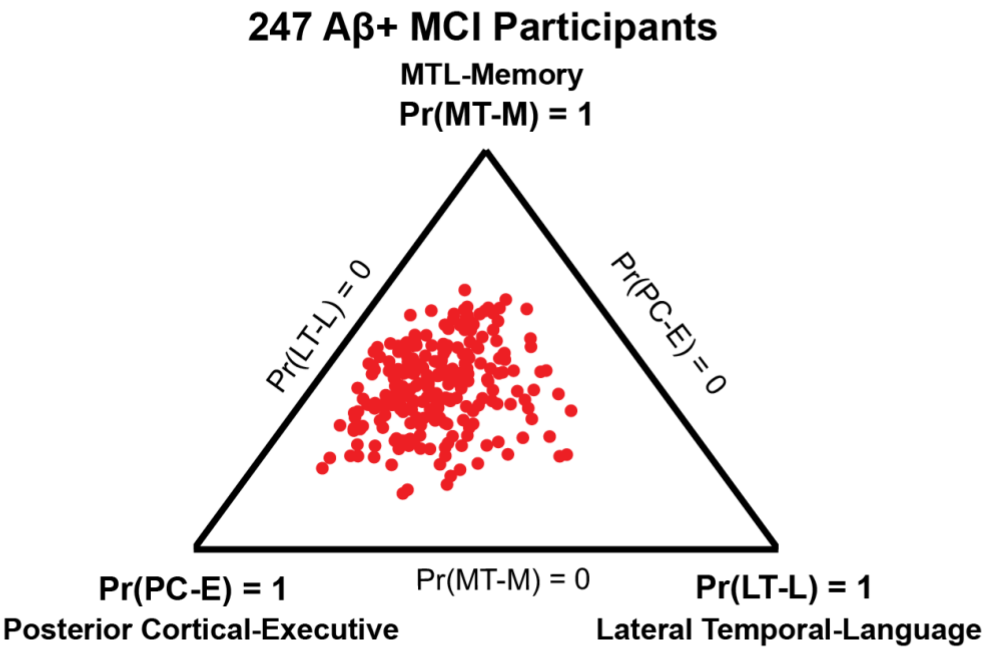


**Figure S14. Factor loadings of 247 Aβ+MCI participants for K = 3 factors.** Each participant corresponds to a dot, whose location (in barycentric coordinates) represents the factor composition. Corners of the triangle represent "pure factors"; closer distance to the respective corners indicates higher probabilities for the respective factors. Most dots are far from the corners, suggesting that most participants expressed multiple factors. "MT-M" indicates the MTL-Memory factor. "PC-E" indicates the Posterior Cortical-Executive factor. "LT-L" indicates the Lateral Temporal-Language factor.

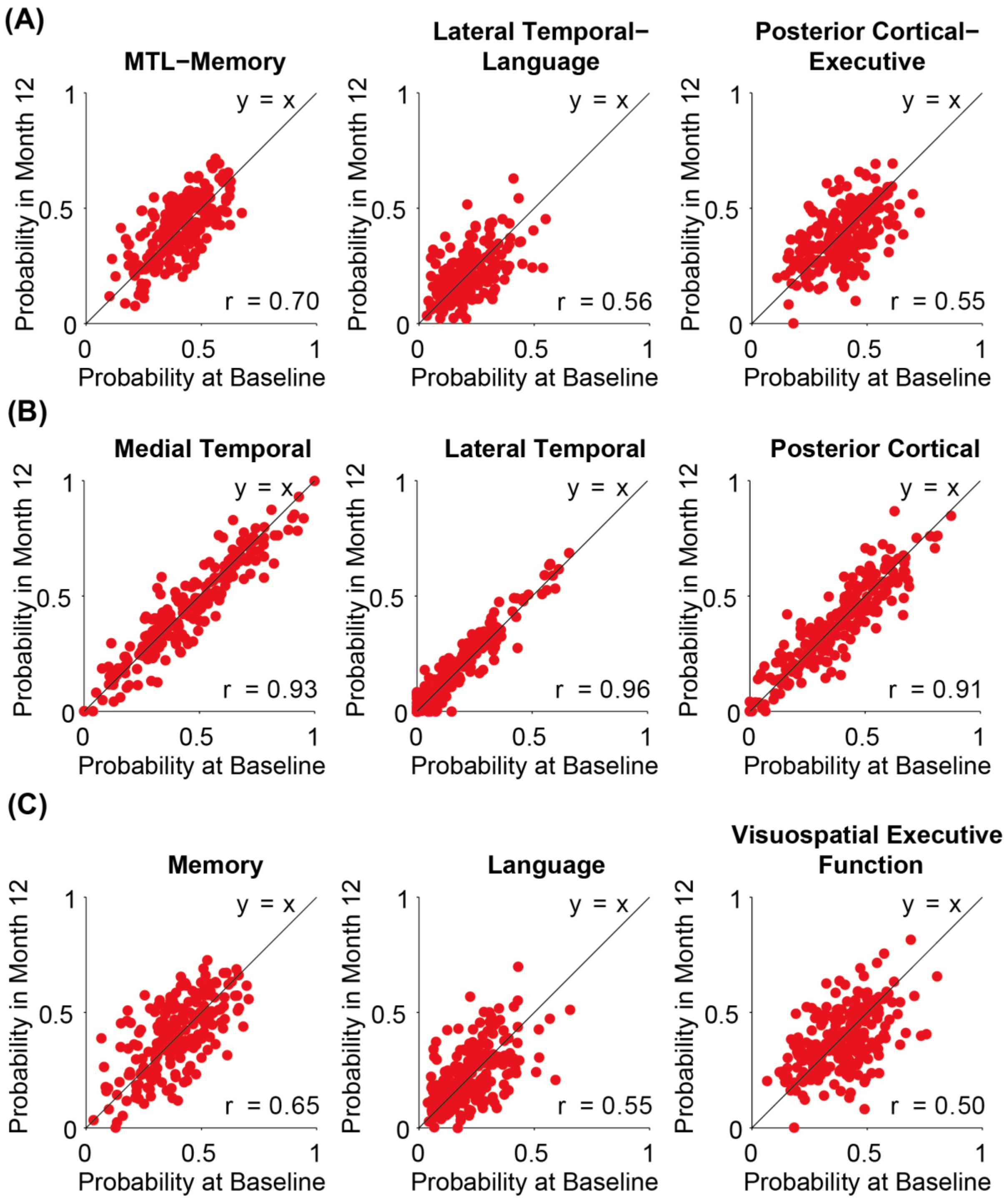


**Figure S15. Stability of factor loadings of Aβ+MCI participants after one year.** Each red dot corresponds to an Aβ+MCI participant. The x axis represents the probability of each factor at baseline and the y axis represents the probability of each factor after one year. (A) Atrophy-cognitive loading estimated using both structural MRI and cognitive scores. (B) Atrophy loading estimated using only structural MRI. (C) Cognitive loading estimated using cognitive scores.

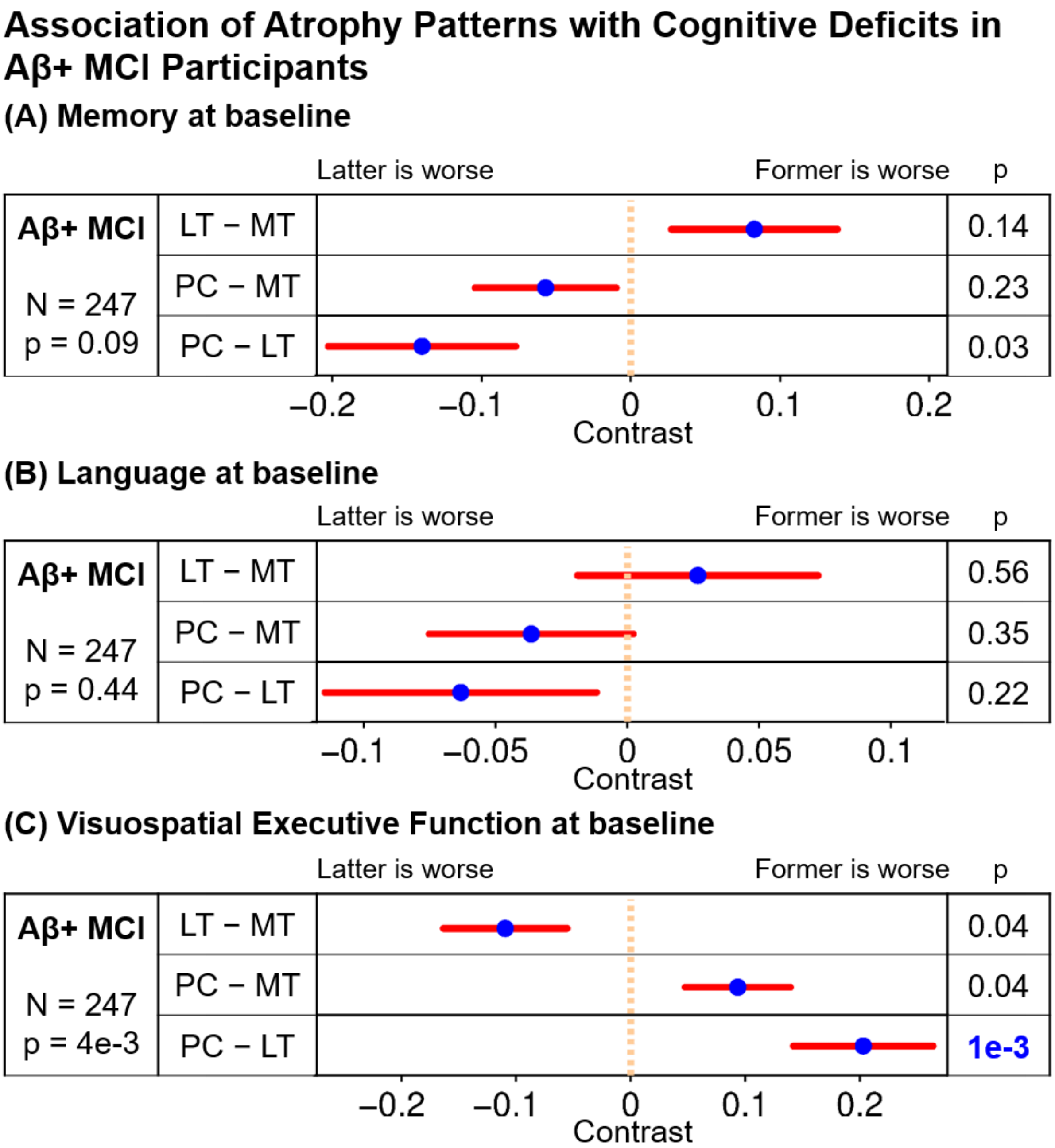


**Figure S16. Associations between atrophy loadings and cognitive loadings among Aβ+ MCI participants.** All p values remaining significant after FDR (q = 0.05) control are highlighted in blue. MT, LT, and PC indicate medial temporal lobe, lateral temporal, and cortical atrophy patterns, respectively. Blue dots are estimated differences in regression coefficients between atrophy loadings and red bars show standard errors.

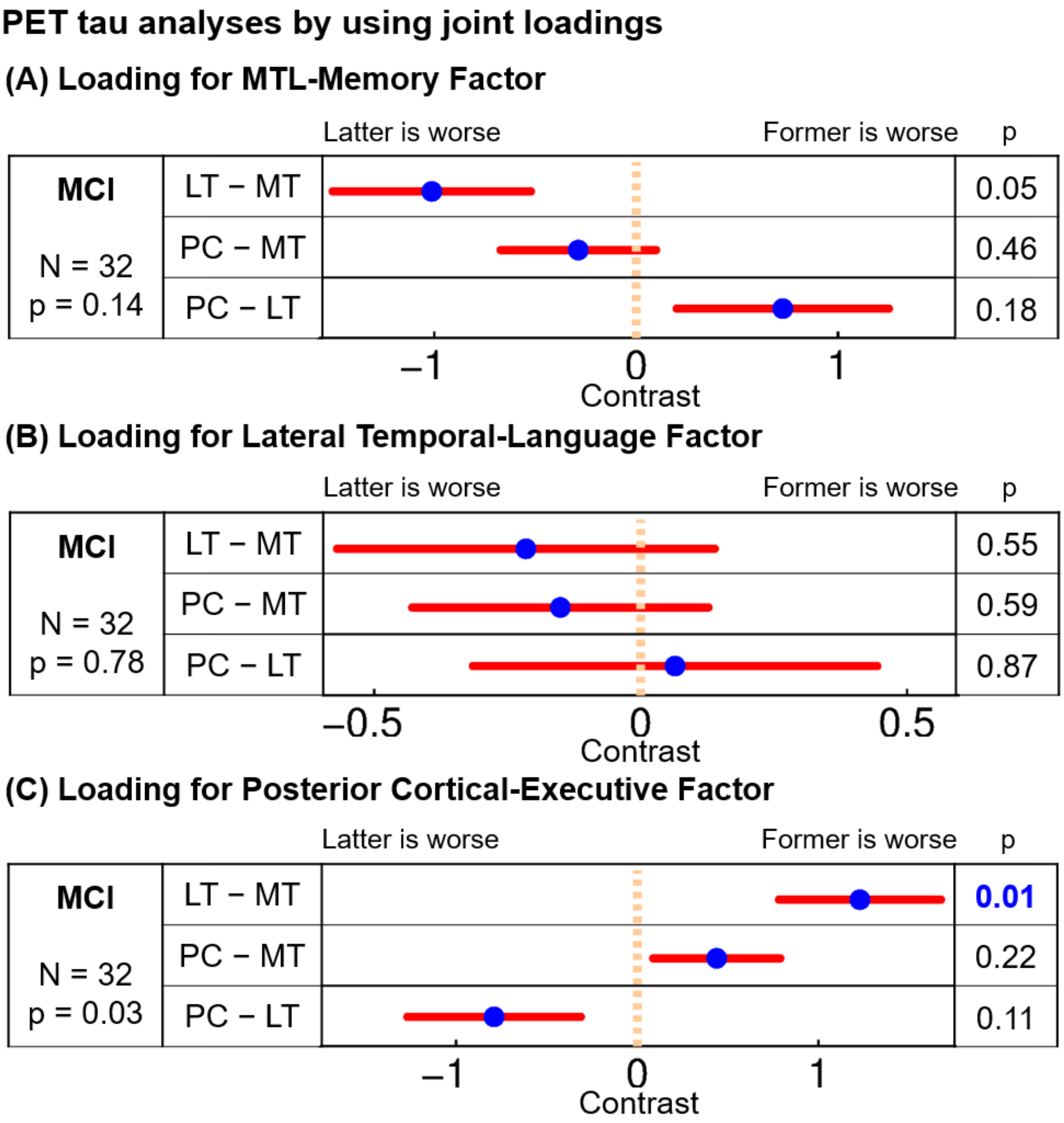


**Figure S17. Associations between factor loadings and Tau deposits in atrophied regions among ADNI-2/3 A+MCI participants.** All p values remaining significant after FDR ($q < 0.05$) are highlighted in blue. MT, LT, and PC indicate Tau deposits in the medial temporal lobe, lateral temporal, and posterior cortical atrophied regions respectively. Blue dots are estimated differences in regression coefficients between Tau deposits in different atrophied regions and red bars show standard errors.

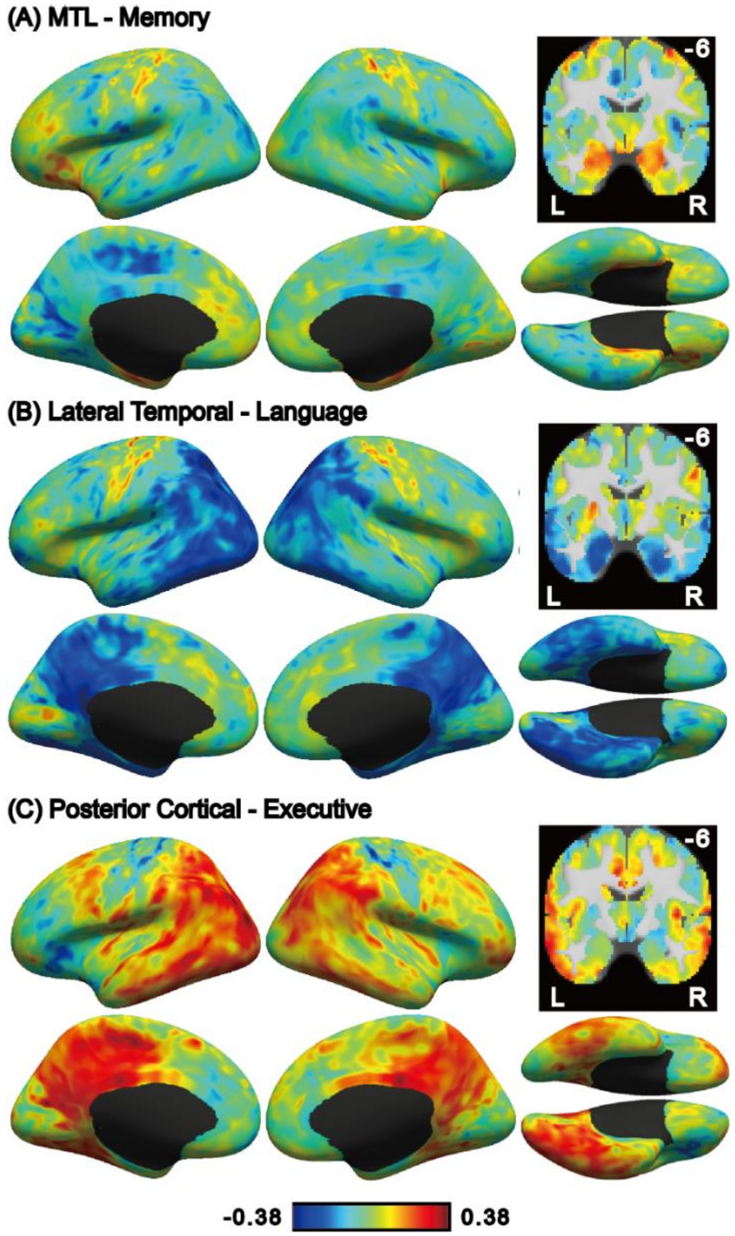


**Figure S18. Correlation between atrophy-cognitive factor loading and Tau deposition (at each voxel) across participants.** (A) Correlation between Tau deposition and MTL-Memory factor loading. (B) Correlation between Tau deposition and Lateral Temporal-Language factor loading. The spatial pattern is similar to the posterior cortical atrophy pattern ($r = -0.40$; $p = 0.013$; significant after FDR correction), suggesting that higher Tau deposition in posterior cortical regions is associated with lower Lateral Temporal-Language factor loading (C) Correlation between Tau deposition and Posterior Cortical-Executive factor loading. The spatial pattern is similar to the posterior cortical atrophy pattern ($r = 0.42$; $p = 0.017$; not significant after FDR correction), suggesting that higher Tau deposition in posterior cortical regions is associated with higher Posterior Cortical-Executive factor loading. We note that the above maps were meant for visualization and in fact, voxel-wise statistical tests were not significant when corrected for multiple comparisons.

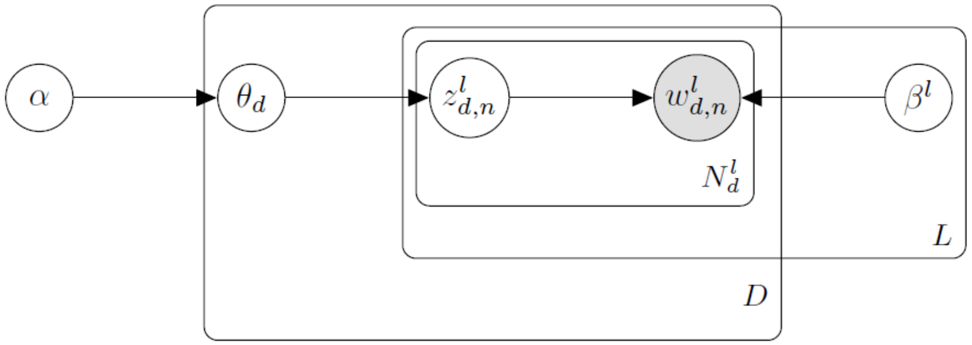


**Figure S19. Graphical model representation of the MMLDA model.** The circled variables represent random variables, while the shaded node represents observed variables. The edges represent statistical dependencies. The variables at the corner of the rectangles indicate the number of times the variables inside the rectangles were replicated. Therefore $\theta_d$ was replicated $D$ times, once for each participant. The variables $z^l_{d,n}$, $w^l_{d,n}$ and $\beta^l$ were replicated $L$ times, once for each modality. For the $d$-th participant, the variables $z^l_{d,n}$ and $w^l_{d,n}$ were replicated $N^l_d$ times, once for one unit in an atrophy voxel or a cognitive score. $\alpha$ is hyperparameter parameterizing the Dirichlet prior on $\theta$. $\theta$ denote Pr(Factor | Participant), $\beta^1$ denote Pr(Voxel | Factor) and $\beta^2$ denote Pr(Score | Factor) . Therefore, $\theta$, $\beta^1$ and $\beta^2$ are matrices, where each row is a categorical distribution summing to one. $z^l_{d,n}$ denote the factor that the voxel or score belongs to and $w^l_{d,n}$ denote the location of observed atrophy voxel or cognitive score.

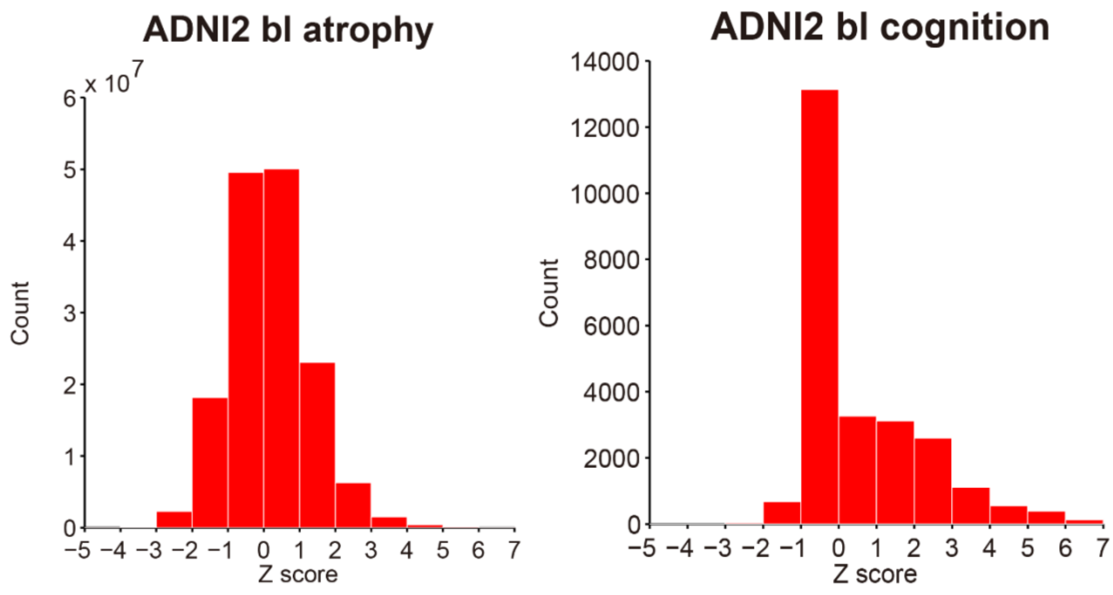


**Figure S20. Histogram of atrophy z score and cognitive z score in ADNI-2. The larger z score corresponds to worse performance.**
